# Modeling neuron-astrocyte interactions in neural networks using distributed simulation

**DOI:** 10.1101/2024.11.11.622953

**Authors:** Han-Jia Jiang, Jugoslava Aćimović, Tiina Manninen, Iiro Ahokainen, Jonas Stapmanns, Mikko Lehtimäki, Markus Diesmann, Sacha J. van Albada, Hans Ekkehard Plesser, Marja-Leena Linne

**Author notes:** These authors contributed equally to this work (Han-Jia Jiang, Jugoslava Aćimović, Hans Ekkehard Plesser, Marja-Leena Linne).

## Abstract

Astrocytes engage in local interactions with neurons, synapses, other glial cell types, and the vasculature through intricate cellular and molecular processes, playing an important role in brain information processing, plasticity, cognition, and behavior. This study advances understanding of local interactions and self-organization of neuron-astrocyte networks and contributes to the broader investigation of their potential relationship with global activity regimes and overall brain function. We present six new contributions: (1) the development of a new model-building framework for neuron-astrocyte networks, (2) the introduction of connectivity concepts for tripartite neuron-astrocyte interactions in biological neural networks, (3) the design of a scalable architecture capable of simulating networks with up to a million cells, (4) a formalized description of neuron-astrocyte modeling that facilitates reproducibility, (5) the integration of experimental data to a greater extent than existing studies, and (6) simulation results demonstrating how neuron-astrocyte interactions drive the emergence of synchronization in local neuronal groups. Specifically, we develop a new technology for representing astrocytes and their interactions with neurons in distributed simulation code for large-scale spiking neuronal networks. This includes an astrocyte model with calcium dynamics, an extended neuron model receiving calcium-dependent signals from astrocytes, and a parallelized connectivity generation scheme for tripartite interactions between pre- and postsynaptic neurons and astrocytes. We verify the efficiency of our reference implementation through benchmarks varying in computing resources and network sizes. Our *in silico* experiments reproduce experimental data on astrocytic effects on neuronal synchronization, demonstrating that astrocytes consistently induce local synchronization in groups of neurons across various connectivity schemes and global activity regimes. By adjusting the strength of neuron-astrocyte interactions, we can switch the global activity regime from asynchronous to network-wide synchronization. This work represents an advancement in neuron-astrocyte modeling, introducing a novel framework that enables large-scale simulations of astrocytic influence on neuronal networks.

**Author summary:** Astrocytes play an important role in regulating synapses, neuronal networks, and cognitive functions. However, models that include both neurons and astrocytes are underutilized compared to models with only neurons in theoretical and computational studies. We address this issue by developing theoretical concepts for representing astrocytic connectivity and interactions and provide a reference implementation supporting distributed parallel computing in the spiking neural network simulator NEST. Using these capabilities, we show how astrocytes help to synchronize neural networks under various connection patterns and activity levels. The new technology makes it easier to include astrocytes in simulations of neural systems, promoting the construction of more realistic, relevant, and reproducible models.

## Introduction

Computational modeling provides methodology and tools to integrate the knowledge about complex molecular and cellular machinery, cell morphology, and spatial organization of astrocytic domains with electrophysiological and imaging data from neurons and astrocytes into models of large-scale neuron-astrocyte circuits. Such circuit-level models can be used to examine how various molecular and cellular mechanisms affect global dynamical regimes and fundamental functions of brain circuits such as sensory processing, learning, and memory. Furthermore, biologically detailed and data-driven models can be used to probe how specific disease-related dysfunctions on genetic, molecular and cellular levels alter global dynamical regimes in brain circuits and contribute to brain disorders. Until recently, network modeling efforts have focused on networks of neurons connected by bipartite chemical synapses [1], occasionally including gap junctions as well [2]. Including astrocytes in large network models creates new challenges for model specification and construction, because they typically form tripartite synapses involving two neurons and an astrocyte. Similar tripartite connectivity is also found elsewhere in the brain, e.g., in the triadic circuitry of the lateral geniculate nucleus [3]. In the spirit of Senk and colleagues [4], we propose here a generic tripartite connection rule, provide an efficient parallel and generalizable reference implementation and demonstrate its capabilities by investigating the effect of astrocytic interactions on synchrony in a neuronal network.

Astrocytes, a prevalent glial cell subtype within the brain, fulfill many roles encompassing the facilitation of synaptic pruning and maturation, regulation of extracellular ionic concentrations, and active participation in metabolic processes as well as the maintenance of the blood-brain barrier functions. *In vivo*, astrocytes are organized into three-dimensional spatially separated domains with little overlap between domains [5–11]. Each astrocytic domain covers tens to hundreds of thousands of synapses in rodents and even up to millions of synapses in humans [5, 6, 9]. Within their domains, astrocytes ramify into complex morphologies containing multiple processes that are positioned close to synapses (see, e.g., [9, 11]) and maintain bidirectional interactions with synapses [12]. Astrocytes react to released neurotransmitters, including glutamate and gamma-aminobutyric acid (GABA), by elevating their intracellular inositol trisphosphate (IP_3_) and calcium levels [13]. Elevated astrocytic calcium levels have been shown to stimulate the release of gliotransmitters, such as glutamate, D-serine, and adenosine triphosphate (ATP) via exocytosis [12–15]. The released gliotransmitters can affect neuronal excitability, for example, the release of glutamate has been shown to induce neuronal extrasynaptic N-methyl-D-aspartate receptor (NMDAR)-mediated slow inward currents (SICs). SICs were first shown in hippocampal neuron-astrocyte co-cultures two decades ago [16], but have since been documented in brain slices of hippocampus and other brain areas as well [17–30], and in humans [31] in addition to rodents.

In the past two decades, the interest in computational modeling of neuron-astrocyte circuits has been steadily increasing (for review and analysis of models see, e.g., [32–36]). However, published models consider only small networks compared to the extent of the biological interaction and are rarely implemented using standard open-access simulation tools or supplemented with code and extensive documentation in public repositories.

This hinders reuse of models and reproduction of the reported results. To address these challenges, we propose here connectivity concepts for the representation of tripartite neuron-astrocyte interaction in large-scale models in the spirit of Nordlie and Senk [4, 37]. We provide a reference implementation for the NEST simulator [38, 39]. Our work complements astrocytic implementations in neural network simulators, namely ARACHNE [40] and Brian2 [41, 42], by focusing (a) on high-level specification of tripartite connectivity patterns and (b) on simulations of large neural networks using distributed parallel computing based on Message Passing Interface (MPI). We propose efficient data structures and algorithms for network construction and simulation, combined with domain-specific language constructs supporting the definition of interactions in large heterogeneous neuron-astrocyte models.

The framework comprises a model for astrocytic calcium dynamics [43, 44], a model for neuron-astrocyte interaction [45], and a method for pairing synapses with astrocytes in neuron-astrocyte circuits. We extend the concept of a synapse, conventionally composed of a presynaptic and a postsynaptic component, with an astrocyte element and develop respective modeling language constructs for tripartite connectivity. The new language constructs allow researchers to complement probabilistic connection rules for neuron populations with specifications for connections with a third factor, such as an astrocyte population. In this framework, we implement neuron-astrocyte interactions through glutamatergic signaling and SICs, based on a previously published model [45] already used in several follow-up studies (e.g., [35, 46, 47]). Due to the long history of modeling slow inward currents and their widespread use in previous studies, we chose to implement SICs in this framework (see all models using SICs in [33, 35]). We verify the efficiency of the data structures and algorithms through a series of benchmarks of the reference implementation in NEST by evaluating the computational load associated with different steps in the simulation while systematically scaling up computing resources and model size. The benchmarks show that the implementation provides an efficient tool for large-scale modeling of neuron-astrocyte circuits.

We demonstrate the capability of the new technology to serve as a platform for *in silico* experiments on neuron-astrocyte circuits. Recent experimental evidence reveals coordinated activity of astrocytic and neuronal populations *in vitro* [48] and *in vivo* in behaviorally relevant tasks [49–52]. Here, we explore a specific mechanism, the emergence of coordinated neuronal and astrocytic activity through the release of glutamate from astrocytes, that induces SICs in postsynaptic neurons. In the central nervous system, SICs contribute to the synchronization of neurons in close proximity, thereby influencing the coordination of local network activity [18, 19, 24, 53].

Additionally, large SICs in cortical areas may hold pathophysiological implications, as they can synchronize neuronal network activity and facilitate the generation of seizures [22, 54], contributing to the pathophysiology of stroke and cerebral edema [30, 55]. Moreover, according to very recent findings, SICs are likely to play an important role in age-dependent physiological and pathological alterations of brain activity in humans, including synaptic plasticity [56]. Brainstem SICs may serve different functions, as previous research [27, 57] has not identified widespread synchronization of these events among neighboring neurons.

We note that SICs exhibit slower kinetics than spontaneous or miniature excitatory postsynaptic currents. The prevailing view is that GluN2B-containing NMDARs drive the generation of SICs [18, 19, 23, 24, 26, 53, 58]. This disparity in kinetic and dynamical properties affects the emergence of coordinated activity in neuron-astrocyte systems.

We construct a spiking neuron-astrocyte network model comparable to a selected experimental study [53, 58] and use it to show that astrocytes robustly enhance neuronal synchrony *in silico* across diverse network activity regimes and connectivities. We fit the parameters of an astrocyte model to reproduce frequency and duration of spontaneous calcium transients reported in the literature [48, 59–61]. The first set of *in silico* experiments distributes neurons across non-overlapping domains, where all neurons in the same domain interact with the same one astrocyte. When synaptic transmission between neurons is suppressed, astrocytes promote synchronized activity among neurons within their respective domains, aligning with experimental findings. In models where neurons fire at biologically realistic frequencies, astrocytes still induce synchrony within their domains, both in a low-rate asynchronous network spiking regime and in the presence of network-wide bursts. Switching from low-rate asynchronous spiking to network-wide bursting is achieved by increasing the strength of inputs from neurons to astrocytes. The bursting regime induces some degree of synchrony between the majority of neurons in the model, but the synchrony in the whole network still remains significantly below the level of synchrony within domains. In the second set of experiments, neurons typically belong to more than one astrocytic domain and interact with more than one astrocyte. Therefore each neuron can belong to more than one synchronous group. We show that, in this case as well, the synchronization within domains still exceeds overall synchronization in the network.

The structure of the present work is as follows. We first provide details of the astrocyte models and the novel tripartite connectivity concepts including their implementation (Methods). We then verify the efficiency of our reference implementation, confirming that it can serve as a platform for the study of large-scale neuron-astrocyte circuits (Results, first subsection). Finally, we demonstrate how the new capabilities can be used to reproduce and extend experimental studies (Results, second subsection).

Preliminary results of this study have appeared in abstract form [62, 63]. The technology described in the present article has been released as open source with several major releases of NEST. The conceptual and algorithmic work is a module in our long-term collaborative project to provide the technology for neural system simulations [39].

## Methods

### Model of neuron-astrocyte interaction

The model of neuron-astrocyte interaction we consider in this work has four components: **(a)** a Li-Rinzel astrocyte model (astrocyte_lr_1994) with calcium dynamics [43, 44], and a mechanism that generates a calcium-dependent SIC [45], **(b)** an extended AdEx neuron model (`aeif_cond_alpha_astro`) capable of receiving the SIC elicited by astrocytes, **(c)** a sic_connection model for transmitting time-continuous SIC signals efficiently from astrocytes to neurons also in parallel simulations, and **(d)** a tripartite connection rule (third_factor_bernoulli_with_pool) extending the standard notion of binary connectivity in neuronal circuits by allowing to establish interactions between triplets of elements facilitating parallel construction of networks including astrocytes. The biological mechanisms modeled with this implementation are illustrated in Fig 1.

**Fig 1.**
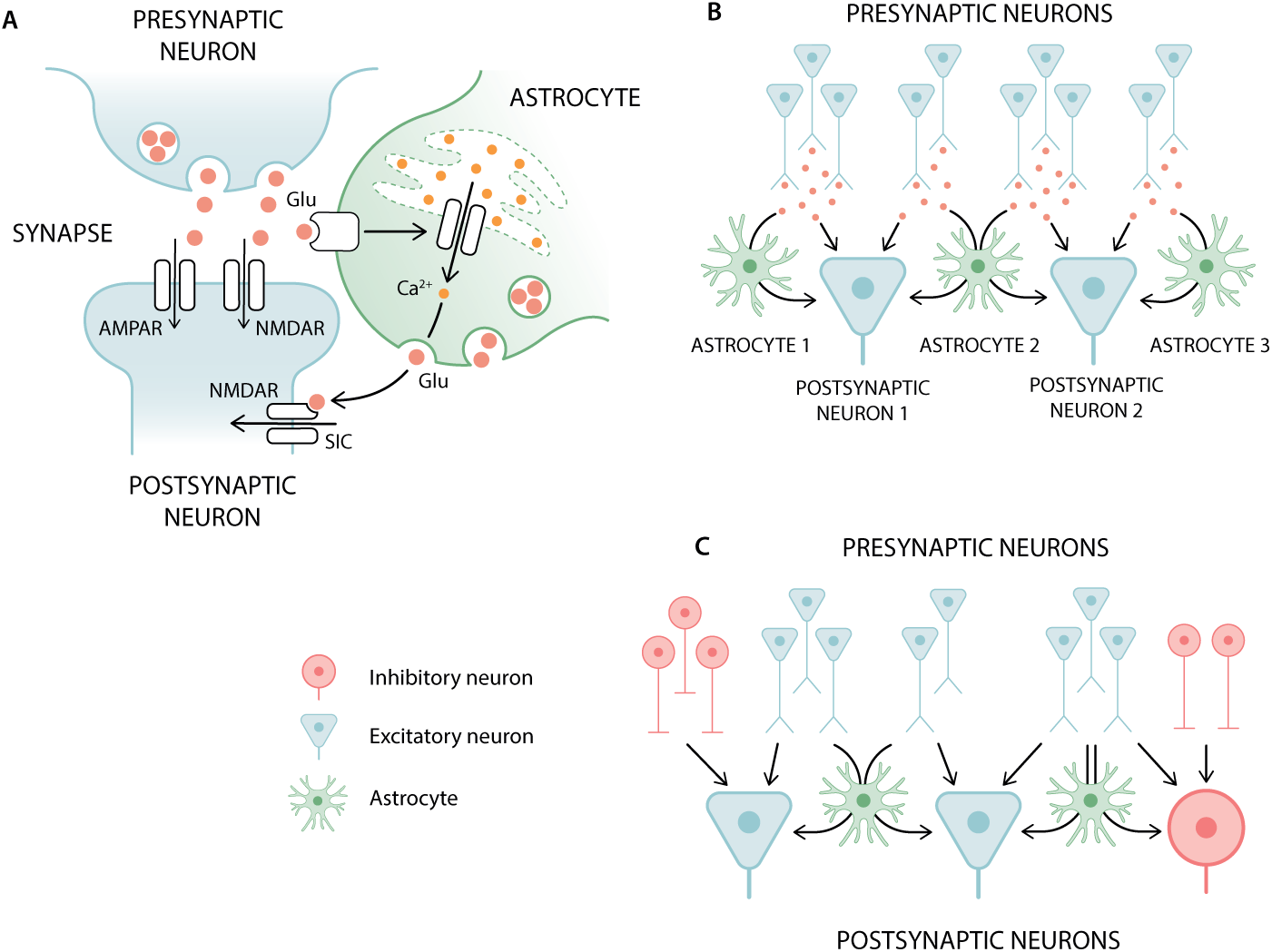
Illustration of the biological mechanisms captured by the model framework for astrocytes. (A) Feed-forward interaction from the presynaptic neuron to the postsynaptic neuron: direct synaptic exchange and an indirect pathway through a neighboring astrocyte. This interaction with an astrocyte is modeled only for the glutamatergic synapses while GABAergic synapses include only interaction between the two neurons. (B) Interaction scheme between astrocytes and excitatory neurons. An astrocyte receives glutamate from all its presynaptic contacts and induces slow inward currents (SICs) only in those postsynaptic neurons that form synapses with its presynaptic contacts. (C) Extension of the interaction scheme in B by inhibitory neurons. Astrocytes are only involved in excitatory-excitatory and excitatory-inhibitory connections.

Reference implementations of the astrocyte and neuron models as well as the SIC interaction were released with NEST 3.6 [64]. A preliminary version of the tripartite connectivity concepts was provided in NEST 3.7 [65], while the definite version of the tripartite connectivity support discussed here was released with NEST 3.8 [38]. A fourth component solely for the purpose of benchmarks presented here is a simplified variant of the astrocyte model, astrocyte_surrogate, which is not integrated into the NEST code base but is provided as an extension module on GitHub [66]. Model documentation and example scripts are available through Read the Docs. Nordlie tables [37] summarize the model equations in S1 Appendix.

The present work requires a two-step mapping of notation. In the first, the mathematical notation of a range of studies needs to be unified to interrelate the components of the network models. In the second, the unified notation needs to be mapped to a domain-specific programming language for expressing the models in executable descriptions. An ontology of variables and parameters together with their default values is given in S2 Appendix. The phenomenological models of the present study, based on phenomenological description of biological mechanisms and the abstractions of the anatomy, require parameter fitting to the experimental data from the literature. These procedures are detailed in S3 Appendix.

### Astrocyte model

We implement an astrocyte model astrocyte_lr_1994 according to the Li-Rinzel reduction [44] of the De Young-Keizer model [43] for intracellular calcium dynamics in nonexcitable cells. The Nadkarni-Jung model extends the Li-Rinzel model by adding astrocytic input and output mechanisms [45]. Our implementation follows the published equations [44, 45] with further adaptations of inputs and outputs. All parameters and their default values are given in S2 Appendix. Here, we provide an overview of the model structure.

Input to the astrocyte induces the production of astrocytic IP_3_, and the change in IP_3_ concentration ([IP_3_]) affects the flux of calcium from the endoplasmic reticulum (ER) to the cytosol through the IP_3_ receptor (IP_3_R) channel. The cytosolic calcium concentration ([Ca^2+^]) determines the output of the astrocyte, which is a SIC to its target neurons. These mechanisms are illustrated in Fig 1A. The equations for the astrocytic dynamics are:

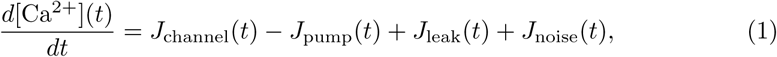

where

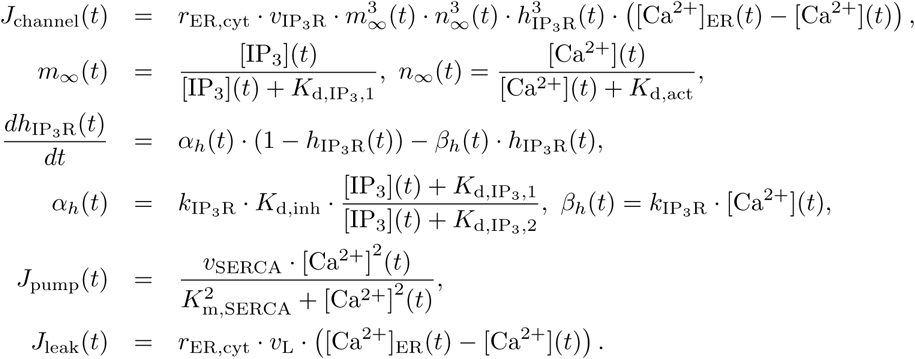

*J*_channel_, *J*_leak_, and *J*_pump_ are the three terms determining the flux of calcium between cytosol and ER. *J*_channel_ is the IP_3_R-mediated release of calcium from the ER to the cytosol, *J*_pump_ refers to the ATP-dependent transport of calcium to the ER via the sarco/ER calcium-ATPase (SERCA), and *J*_leak_ is the leak of calcium between the ER and cytosol. In addition to these standard terms, we include the term *J*_noise_(*t*) to account for natural fluctuations in calcium dynamics.

The parameters and variables involved in Eq (1) are as follows. *r*_ER,cyt_ is the ratio between astrocytic ER and cytosol volumes. *v*_IP3 R_ is the maximum rate of calcium release via astrocytic IP_3_Rs. *K*_d,IP3, 1_ and *K*_d,IP3,2_ are the first and second astrocytic IP_3_R dissociation constants of IP_3_. *K*_d,act_ and *K*_d,inh_ are the astrocytic IP_3_R dissociation constants of calcium (for activation and inhibition, respectively). *k*_IP3 R_ is the astrocytic IP_3_R binding constant for calcium inhibition. *v*_SERCA_ is the maximum rate of calcium transport by astrocytic SERCA pumps. *K*_m,SERCA_ is the half-activation constant of astrocytic SERCA pumps. *v*_L_ is the rate constant of calcium leak from astrocytic ER to cytosol. [Ca^2+^]_ER_ is the calcium concentration in the ER. *m_∞_*(*t*) and *n_∞_*(*t*) are steady-state values for two gating variables of IP_3_Rs. *h*_IP3 R_(*t*) is the fraction of IP_3_Rs that are not yet inactivated by calcium. *α_h_*(*t*) and *β_h_*(*t*) are two rates of the exchange between activated and inactivated IP_3_Rs.

The calcium conservation is enforced by calculating the ER calcium concentration according to the cytosolic calcium concentration at every simulation time step, similarly to [44, 45]:

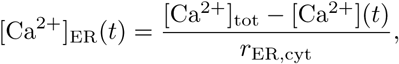

where [Ca^2+^]_tot_ is the parameter determining the maximal cytosolic calcium concentration, i.e., the cytosolic calcium concentration given that all the calcium ions are in the cytosol (see [43] for details). Thus, the total amount of calcium in the astrocyte (cytosol + ER), is kept constant. In our astrocyte model, the cytosolic calcium concentration is forced to stay within the range between zero and [Ca^2+^]_tot_ at each simulation step. Therefore, the given noise *J*_noise_(*t*) does not produce negative calcium concentration.

The astrocyte receives presynaptic spike events as excitatory inputs, similarly to previous studies [40, 42], which initiate IP_3_ production given by

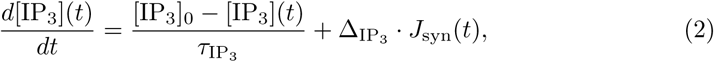

where [IP_3_]_0_ is the baseline value of [IP_3_], *τ*_IP3_ is the time constant of the exponential decay of [IP_3_], Δ_IP3_ is the parameter determining the increase in [IP_3_] induced by the input, and *J*_syn_(*t*) is the sum of all excitatory synaptic inputs (in the form of delta functions) that the astrocyte receives at time *t*.

The output of the astrocyte is a SIC to its targets. This current is determined by [Ca^2+^] with the phenomenological expression [45], given by

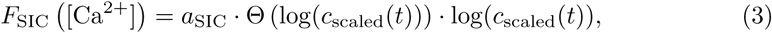

where

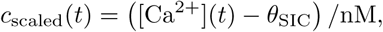

Θ(*·*) is the Heaviside function, and SIC is generated (*F*_SIC_ *>* 0) when the calcium concentration is larger than *θ*_SIC_ + 1 nM. *F*_SIC_ [Ca^2+^] is unitless, and the correct unit (pA) for the summed astrocyte-induced SIC in a neuron (*I*_SIC_; see SIC-receiving neuron model) enters when *F*_SIC_ [Ca^2+^] is multiplied by the weight of astrocyte-to-neuron connections (*w*_astro to post_) which has a unit of pA. Note that due to the Heaviside function, *F*_SIC_ *≥* 0 everywhere and does not diverge to *−∞* for small *c*_scaled_. Our implementation allows the users to change *θ*_SIC_ in this equation, and specify a factor *a*_SIC_ to scale the generated SIC. These implementation choices add flexibility to Eq (9) in the original publication [45], where the parameters *θ*_SIC_ and *a*_SIC_ are fixed. Here, we use the notation with /nM to indicate that *c*_scaled_ is the dimensionless version of the given concentration in nM.

### SIC-receiving neuron model

Next to a mathematical model of an astrocyte, we provide a model of single neuron dynamics susceptible to the SICs astrocytes induce in a neuron. For the purpose of the present work, our starting point is the AdEx neuron model with conductance-based synapses [67, 68]. An additional current (*I*_SIC_) in the differential equation describing the membrane potential represents the summed input from all connected astrocytes. The adapted equation of the combined model (`aeif_cond_alpha_astro`) reads

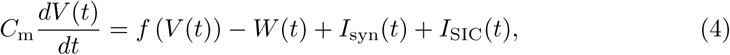

where *C*_m_ is the membrane capacitance, *V* (*t*) is the membrane potential, *f* (*V* (*t*)) defines the passive properties and the spiking mechanism, *W* (*t*) is the adaptation variable, *I*_syn_(*t*) is the summed synaptic current to the neuron, and *I*_SIC_(*t*) is the summed SIC induced by all connected astrocytes. *f* (*V* (*t*)) and *W* (*t*) follow Eqs (2) and (3) in [67] respectively, and *I*_syn_(*t*) is given by

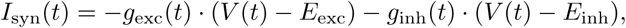

where *g*_exc_ and *g*_inh_ are the summed excitatory and inhibitory synaptic conductances, and *E*_exc_ and *E*_inh_ are the excitatory and inhibitory reversal potentials.

When *V* (*t*) reaches the spike detection threshold *V*_peak_, the reset condition is applied:

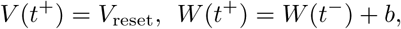

where *V*_reset_ is the reset value for *V* after a spike, and *b* is the spike-triggered adaptation. The membrane potential *V* (*t*) is then held at the reset value *V*_reset_ for a refractory period *t*_ref_.

Fig 1B illustrates a situation where an excitatory neuron receives a SIC from two astrocytes, while astrocytes receive inputs from several excitatory presynaptic neurons. Fig 1C extends Fig 1B by adding also inhibitory cells.

### SIC connection model

The SIC connection is implemented as a unidirectional interaction from an astrocyte to a target neuron. The connection is dependent on two parameters, the weight that defines the strength of interaction and the connection delay. The interaction is continuous and dependent on the state of the source astrocyte, but independent of the state of the target neuron. This combination of features distinguishes SIC connections from other cell-to-cell interactions available in NEST, such as synaptic connections or gap junctions. Therefore, simulation codes for neuron-astrocyte networks require specific software support for the interaction through SIC, and mathematical models of neuronal networks need to specify this type of interaction if desired. In the present work this interaction is specified as `sic_connection`.

### Concepts for tripartite connectivity

The recent review of connectivity concepts in the computational neuroscience literature by Senk et al. [4] concentrates on the connection patterns between two populations of neurons and presents an ontology. Neuron-astrocyte networks, however, establish a three-party connectivity. This suggests an interdependence in the connections among three network nodes that goes beyond pairwise probabilities, highlighting innate network motifs in this type of connectivity. A motif is a common graph-theoretic concept extensively used to describe networks with complex connectivity, including neuronal networks [35, 69–72].

Details of the anatomical structure of neuron-astrocyte circuits are still a matter of debate in spite of experimental progress [8–11]. We therefore here develop a flexible approach for generating tripartite connectivity. First, our approach supports a wide range of primary connection rules [4]. Second, we introduce a third-factor connection rule, the *third-factor Bernoulli with pool* rule (TBP rule), which in itself supports three alternative statistical descriptions of the structure of neuron-astrocyte connectivity.

We propose to specify tripartite connectivity as follows. Given source and target neuron populations, a *primary connection rule* selects specific source-target neuron pairs and establishes primary connections between the source and target neuron. For each source-target pair selected, the *third-factor connectivity rule* then determines whether and which element of a third-factor population (astrocytes in our case) to attach to the primary connection. Attachment means that further connections are created from the primary source neuron to the third-factor element and from the third-factor element to the primary target neuron. For the sake of clarity, we will refer the third factor as astrocytes in what follows, but point out that the rules and their implementation are entirely general: The only requirement on the elements of the third-factor population is that they can receive the type of output provided by the primary source neurons and emit output that can be received by the primary target neurons.

In the TBP rule, the selection of the specific astrocyte to attach to a given source-target pair is based on *pools*, i.e., subsets of the astrocyte population. Fig 2 illustrates alternative pool types and resulting connection patterns. Each neuron in the target population is assigned a fixed pool of a user-defined pool size *S*_pool_. We distinguish between two different pool types: random and block pools. For random pools, the members of the pool are selected at random from the entire astrocyte population without replacement. Any given astrocyte can be part of the pools of multiple target neurons and pools assigned to different target neurons typically share some members as illustrated in Fig 2A. This random pool formation corresponds to the *random fixed in-degree without multapses connection rule* as defined by Senk et al. [4], albeit for potential, not actual connections. For example, in Fig 2A each target neuron could, but does not necessarily, connect to two astrocytes.

**Fig 2.**
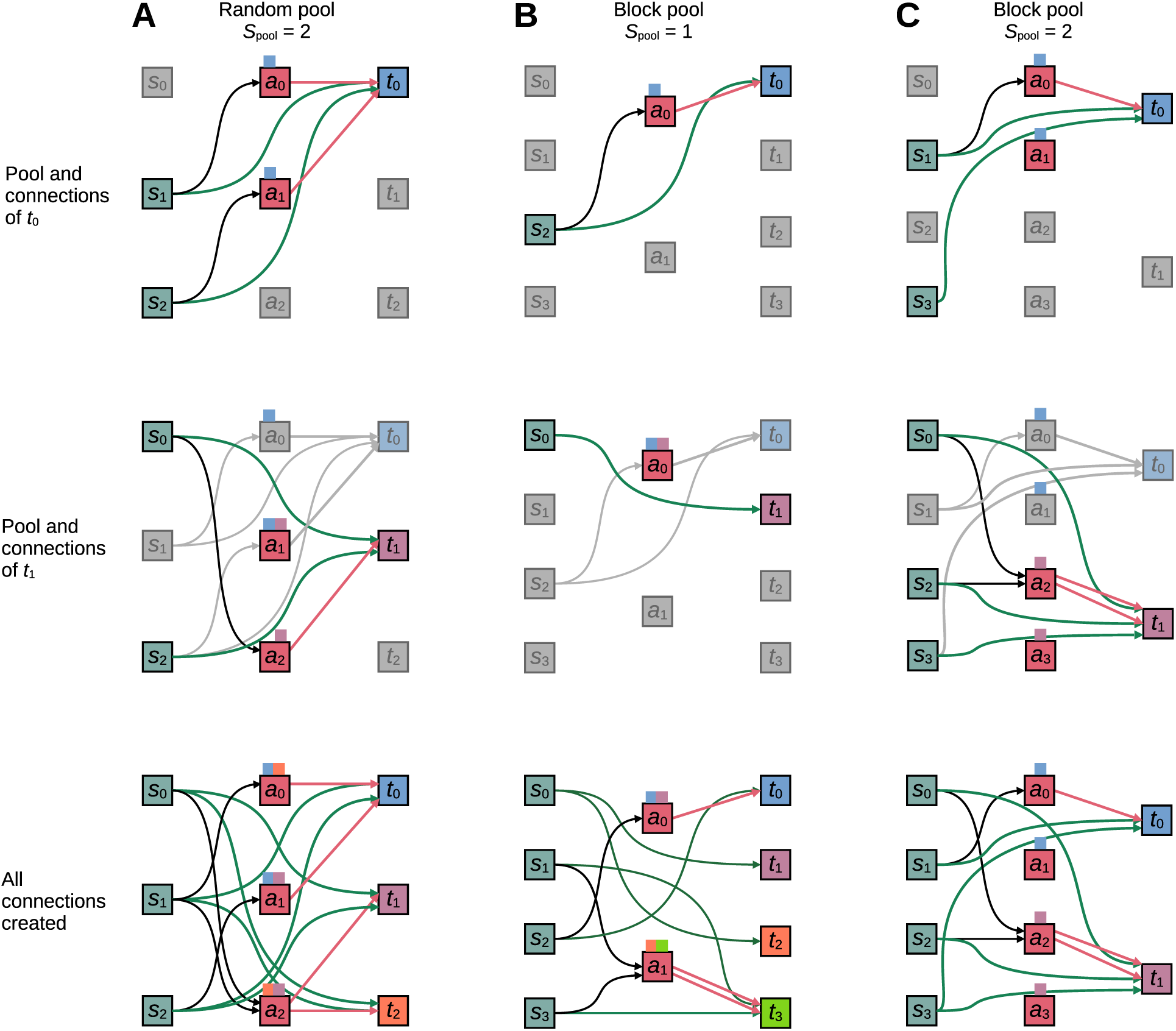
Examples of concept of tripartite neuron-astrocyte connectivity. Green arrows between source (*s_j_* squares) and target (*t_i_* squares) neurons indicate synapses. In the tripartite concept, a synapse can be attached to an astrocyte (*a_k_*) with a certain probability, e.g., *t*_1_ and *t*_2_ in (A) have both attached and unattached synapses. Black and magenta arrows indicate astrocyte attachment (i.e., neuron-to-astrocyte and astrocyte-to-neuron connections) of the synapses. Colored squares on top of astrocyte symbols indicate which astrocyte pool(s) they belong to, but do not determine actual connections. (A) Random pools. Each target neuron interacts with up to *S*_pool_ randomly selected astrocytes. (B) Block pools with *S*_pool_ = 1. Each target neuron interacts with up to one astrocyte. The number of target neurons is a multiple of the number of astrocytes. (C) Block pools with *S*_pool_ *>* 1. Each target neuron interacts with up to *S*_pool_ astrocytes. The number of astrocytes is *S*_pool_ times the number of target neurons. *s*_0_*, s*_1_*, …*: source (presynaptic) neurons; *t*_0_*, t*_1_*, …*: target (postsynaptic) neurons; *a*_0_*, a*_1_*, …*: astrocytes. From (A) to (C), top two rows show the astrocyte pools and connections associated with the first two target neurons. The bottom row shows the total connectivity of an example instance. The diagram and notation follow [4].

Block pools map astrocytes to target neurons deterministically in two possible ways. For pool size *S*_pool_ = 1, exactly one astrocyte is assigned to each target neuron, while each astrocyte can connect to multiple target neurons as shown in Fig 2B. In this case, the number of target neurons must be a multiple of the number of astrocytes. For block pools of size *S*_pool_ *>* 1, each target neuron selects exactly *S*_pool_ astrocytes; these pools do not overlap. The total population of astrocytes needs to be *S*_pool_ times larger than the number of target neurons. Fig 2C shows an example for *S*_pool_ = 2. As for the random pool, the block pool also defines potential instead of actual connections.

Note that the TBP rule allows multiple connections between a given astrocyte-target neuron pair. If a target neuron receives input from multiple source neurons, an astrocyte can be attached to each of those connections and the same astrocyte may be selected for each of them. Fig 2B provides an example: Target *t*_3_ receives input from sources *s*_1_ and *s*_3_ and astrocyte *a*_1_ is attached to both connections, resulting in two connections from *a*_1_ to *t*_3_; Fig 2C shows another example. In both cases, a neuron interacts with the same astrocyte at multiple synapses, as commonly observed in nature [73]. For further details on connection probabilities, see S3 Appendix.

The examples in Fig 2 show that the TBP rule alone can give rise to a wide range of tripartite connection patterns, in particular as the TBP rule can be combined with various binary connection rules for the primary connections. We therefore restrict ourselves to exploring the consequences of this rule in the remainder of this work.

### Efficient parallel instantiation of a tripartite connection rule

Efficient parallel instantiation of network connectivity from rules, as needed for large-scale network simulations, benefits from simulation software that explicitly implements high-level rules in low-level code to optimally exploit parallelism. While some neuronal network simulation software provides such support for pairwise connectivity [4], there has been little support for tripartite connection patterns. We show here how such support can be integrated into parallel instantiation of high-level connection rules in the NEST simulator.

Our approach has the following requirements: (i) a primary connection rule specifying connections between source and target neurons; (ii) a third-factor connection rule specifying if and how to attach a third-factor element to a given primary connection; (iii) the third-factor element is attaching to a primary connection by forming two additional connections, one from the primary source to the third-factor element and another from the third-factor element to the target. Beyond this, our approach is entirely general.

To achieve memory- and time-efficient representation and creation of connectivity, NEST generates and stores connectivity on the target (postsynaptic) side, i.e., on the virtual process (VP) managing the target neuron of a connection [74, 75]. VPs can be mapped arbitrarily to OpenMP threads and MPI processes [76]. Creating and storing connections on the VP of the connection target causes complications when creating tripartite connectivity, since the third-factor element (astrocyte) selected may be managed by a different VP than the primary target. We solve this as follows: The VP managing the target neuron first applies the primary connection rule to decide which primary connection to create next. For the selected source-target pair, it then invokes the third-factor connection rule to decide whether to create a third-factor connection, and if so, which third-factor element to attach to the primary connection. The VP then creates a “third out” connection from the third-factor element to the primary target immediately. It further stores the element IDs of the primary source and the selected third-factor element in a buffer local to the VP. Once all primary and “third out” connections have been created, the VPs exchange the buffered primary-source–third-factor pairs and each VP then creates the “third in” connections from the selected primary sources to the third-factor elements it manages. This approach allows for a flexible combination of primary and third-factor connection rules and preserves the scaling properties of the primary connection rule. The algorithm adds an overhead for tripartite connectivity that is linear in the number of primary connections and incurs only a single round of interprocess communication for the exchange of the primary-source–third-factor pairs. Implementation details are described in S4 Appendix.

We would like to point out that the approach as described is not restricted to the TBP rule: it applies to any tripartite connection rule fulfilling the mild requirements mentioned above. Indeed, while we so far only support the TBP rule as part of our reference implementation in the NEST simulator, users could add new third-factor rules without modifying existing NEST source code by implementing a corresponding C++ class in a NEST extension module.

Fig 3 illustrates how tripartite connectivity can be created in practice in NEST 3.8 and later. A key feature of our approach is that the specification of the primary and the third-factor connectivity are independent of each other. To create tripartite connectivity with a different primary rule, one just needs to specify the desired rule and its parameters as the conn_spec parameter. The third_factor_conn_spec parameter must at present always select the TBP rule, as no other rules are available yet, but the details of the rule, i.e. attachment probability, pool type, and pool size, can be varied by the user. The synapse specifications through parameter syn_specs are in turn entirely independent of the connectivity specifications and also of each other. The only requirement on the synapse models specified is that they are compatible with the neurons and third-factor elements to be connected. New rules for third-factor connectivity fit seamlessly into this specification scheme: The user only needs to provide the name of the new rule and its parameters in the third_factor_conn_spec parameter.

**Fig 3.**
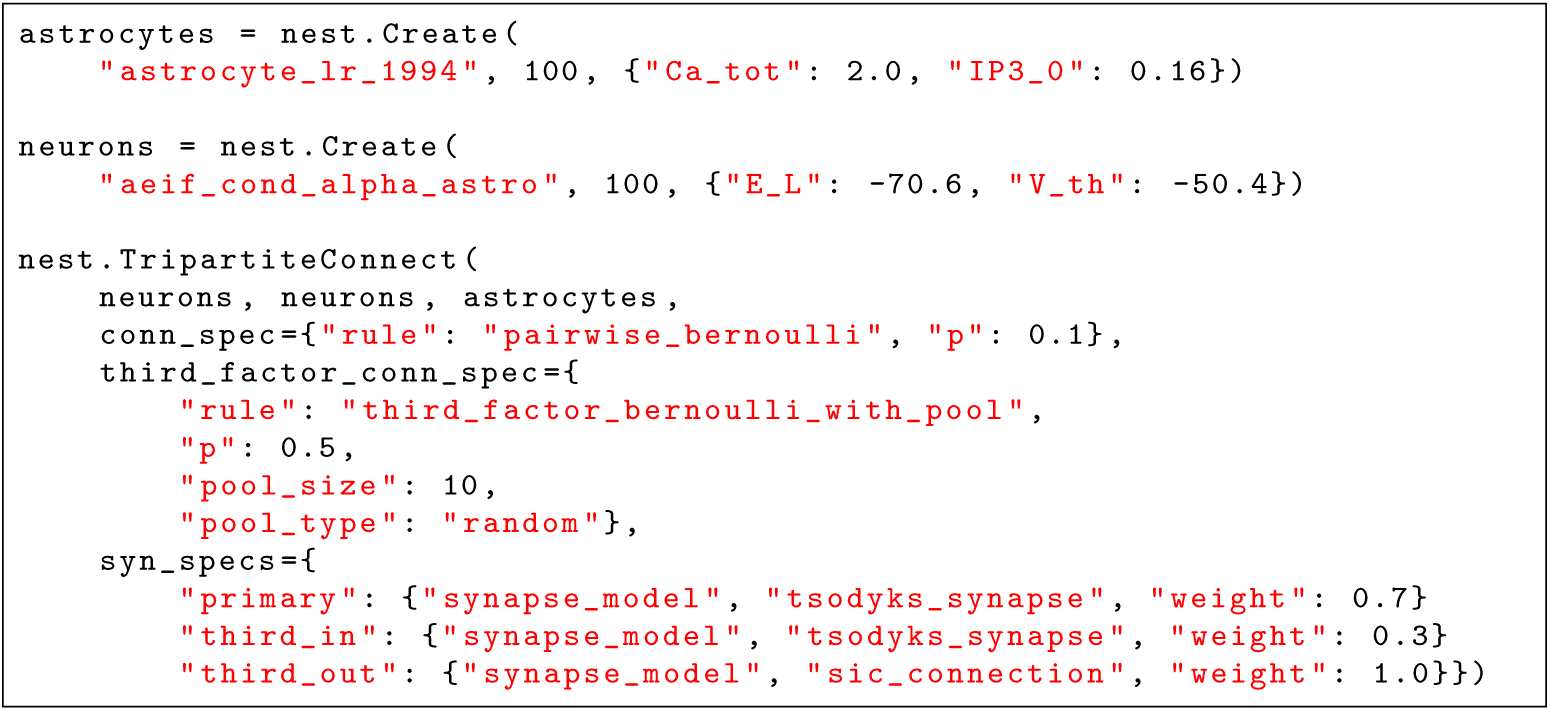
Example of tripartite connectivity in NEST. The executable description specifies a network of 100 astrocytes and 100 neurons and tripartite connectivity. The conn spec parameter specifies the primary connection rule, here pairwise Bernoulli connectivity with 10% connection probability. The `third factor conn spec` parameter specifies the rule for attaching third-factor elements, in this case the TBP rule with 50% probability to attach an astrocyte to a given primary connection. The astrocyte to be attached will be drawn with equal probability from a random pool of 10 astrocytes. Each target neuron has its own, random but fixed astrocyte pool to choose from. The syn specs parameter specifies the properties of the actual synapses to be created.

### Benchmark methods

We evaluate the performance of our NEST reference implementation for astrocytes by simulating neuron-astrocyte network models using the beNNch framework for reproducible benchmarks of neuronal network simulations [77]. The metrics for performance is the wall-clock time (*T*_wall_) spent on network creation, network connection, and state propagation. Network creation and network connection capture the time spent on creating and connecting cells and recording devices, respectively.

State propagation captures the time required to advance the dynamical state of the network by the requested span of model time [77]. Network creation time in our benchmarks is always so short (*<* 50 ms) that we do not include it in the results for the sake of conciseness.

Three design goals guide the choice of network model used for benchmarks: It should be indicative of performance for large-scale models, it needs to be scalable to arbitrary network size with minimal changes in firing rates, and it should capture an average to worst case scenario. In accordance with widely established practice, we choose a ratio of excitatory to inhibitory neuron numbers of 4:1 [78, 79] and equal numbers of neurons and astrocytes (see, e.g., [80] referring to astrocyte-to-neuron ratios of 1:3 to 1.4:1 in the mammalian cortex). Sparse random connectivity with average in-degrees held constant ensures comparable firing rates independent of network size (but see [81]). Unless otherwise specified, the primary connectivity follows the pairwise Bernoulli rule [4]. The TBP rule with random pools adds astrocytic connections with fixed probability to all excitatory connections. Excitatory Poisson spike trains with fixed firing rate provide an external drive for neurons. Parameters for the astrocyte model are based on [43–45], who investigate models with a rather high level of excitability. These astrocyte parameter values are also the default values in NEST 3.8. Overall, we consider this network configuration to represent a high but realistic computational load for large-scale neuron-astrocyte models. The model structure is described in detail in S1 Appendix, Table A, and parameters are given in Table B. To assess the effect of different network dynamics on simulation runtime, we use two model variants which only differ in delay and synaptic time constant for inhibitory connections (see S1 Appendix, Table B): Default parameter values give rise to sparse spiking activity (“Sparse” model), while modified parameter values lead to synchronous activity (“Synchronous” model). Fig 4 shows the dynamics of the two benchmark models.

**Fig 4.**
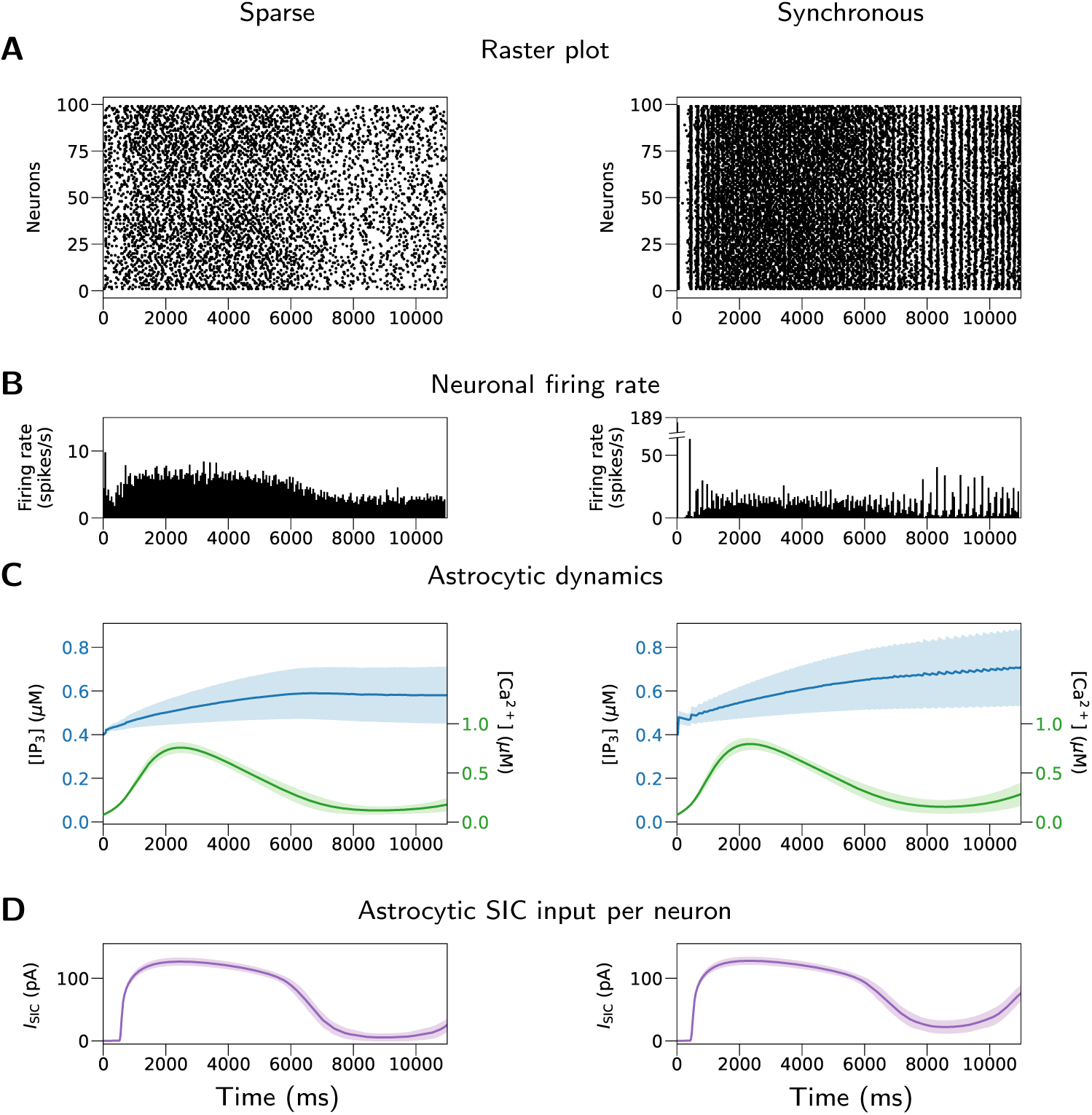
Dynamics of the “Sparse” and “Synchronous” models used for benchmarks. (A) Raster plot of neuronal firings. (B) Neuronal firing rate. (C) Astrocytic [IP_3_] and [Ca^2+^]. (D) Astrocytic SIC input (*I*_SIC_) per neuron. Shaded areas: standard deviations across cells. The left column shows a simulation with the “Sparse” model, where the mean neuronal firing rate is 4.74 spikes/s and the mean pairwise spike count correlation of sampled neurons is 0.014. The right column shows a simulation with the “Synchronous” model, where the mean neuronal firing rate is 12.0 spikes/s and the mean pairwise spike count correlation of sampled neurons is 0.072. Data from *t* = 1 to *t* = 11 s of the simulation are used for the calculation of firing rate and correlation. For the specifications of the models, see Tables A and B in S1 Appendix. The examples shown here are for scale one, i.e., 20,000 cells in total (neurons and astrocytes). The data shown are (A) the first 100 neurons, (B) all neurons, and (C,D) 100 randomly sampled neurons or astrocytes. Firing rates were averaged over all 10,000 neurons and correlations computed across 100 neurons with 10 ms bin width.

In order to measure the cost of integrating complex astrocyte dynamics given by Eqs (1)–(3) on the run time of a simulation, we define an astrocyte_surrogate model in a NEST extension module [66]. The surrogate model is a copy of the astrocyte_lr_1994 model, however it neither includes astrocytic dynamics nor processes incoming spikes. It only emits a continuous, user-configurable and constant SIC to its targets. Therefore, it serves as a surrogate that gives the same type of output as the astrocyte but costs minimum computing resources. We create a benchmark model called “Surrogate” that has the same parameters as the “Sparse” model but replaces astrocyte_lr_1994 with astrocyte_surrogate. We compare the benchmark results of the “Surrogate” model with the “Sparse” model in order to reveal the cost for the calculation of astrocytic dynamics in the network. In the “Surrogate” model, the pre-defined SIC output of astrocyte_surrogate is set to such value that the overall neuronal firing rate is close to the “Sparse” model.

The second set of benchmarks compares the performance across four connectivity rules: pairwise Bernoulli, fixed in-degree, fixed out-degree, and fixed total number; for details on these rules see [4]. We choose the parameters for these rules so that the expected number of primary connections in the network is the same as for pairwise Bernoulli connectivity in the “Sparse” model; synapse parameters are unchanged.

The third set of benchmarks compares the performance for different astrocytic pool sizes, i.e., upper bounds for the number of astrocytes that each neuron can receive input from. The models used in this set of benchmarks are versions of the “Sparse” model with *S*_pool_ = 10, 100, 1,000, 10,000.

The fourth set of benchmarks test the performance when the cell number is further scaled up to one million. In this set of benchmarks, we use the fixed in-degree rule for primary connectivity and use pool sizes of 10 and 1,000.

As in [77], we perform strong and weak-scaling benchmarks for the described models. Strong scaling shows how fast a solution can be achieved if increasingly more resources are invested, whereas weak scaling shows how time to solution changes if resources are increased proportional to problem size, i.e., scale of the model. If the simulation performance is perfect, the time to solution should decrease proportionally to the amount of resources invested in strong scaling, and should remain approximately the same in weak scaling, due to the distribution of workload. In this sense, in our study the strong-scaling benchmarks measure the simulation time of a model while up-scaling the number of compute nodes, and the weak-scaling benchmarks measure the simulation time while the model size is up-scaled proportionally to the number of compute nodes. The performance can thus be evaluated by the simulation time measured at different scales. In weak scaling, in order to keep the same level of model activity, the cell number is scaled but the expected number of connections per cell is not, i.e., the expected number of primary (neuron-to-neuron) and third-factor (neuron-to-astrocyte and astrocyte-to-neuron) connections per cell is preserved.

### Use case: Constructing neuron-astrocyte network model based on the experimental literature

As a concrete scientific use case for the technology presented here, we create a model mimicking experimental work by Pirttimaki et al. [48] in slice. We use the same network structure as for the benchmarks (see S1 Appendix, Table A), but with modified parameters given in S1 Appendix, Table C. While some model parameters are fixed, others are fitted to experimental data as described below. Specifically, our use case consists of 500 neurons and 100 astrocytes; the neuron to astrocyte ratio is consistent with experimental preparation used in [48], brain slices from rats at P10-P16 and P19-P21. Pairwise Bernoulli and TBP rules, respectively, are used to establish primary and third-factor connections. TBP rule was used with either block or random pool in different versions of the model; the selected astrocyte-to-neuron ratio allows for both pool types. Flexibility of these rules, illustrated in Fig 2, supports analysis of the role of complex neuron-to-astrocyte connectivity, hypothesized in [48]. Poisson trains with fixed rate provide external drives for neurons and astrocytes. Gaussian noise is applied to neurons and astrocytes in the form of electrical current and calcium fluctuation, respectively, as sources of variation in dynamics. S3 Appendix contains additional details of the model construction procedure and supplementary results.

### Experimental data from the literature

We first collect experimental measures of spontaneous calcium activity in astrocytes [17, 48, 59–61] as well as neuronal spiking activity under several experimental conditions and in different brain regions [82, 83]. These experimental measures are compiled in S3 Appendix. They are used to fit some of the model parameters, and to guide selection of other parameters.

### Fitting astrocyte parameters

Astrocyte parameters are selected by simulating Eqs (1) and (2) for 10^5^ parameter sets and evaluating the resulting dynamics. Astrocyte dynamics are driven by Poisson inputs through IP_3_Rs (through *J*_syn_ in Eq (2)). The 10^5^ evaluated parameter sets differ in frequency of the Poisson events (*λ*_Poiss,A_), the intensity with which they affect IP_3_Rs (Δ_IP3_), the total calcium concentration in terms of the cytosolic volume ([Ca^2+^]_tot_), the steady state value for IP_3_ ([IP_3_]_0_), and the time constant that determines the speed of convergence towards the steady state (*τ*_IP3_). These five parameters are randomly sampled from the predefined intervals (see S3 Appendix) while other parameters are kept constant at NEST default values. From the initial 10^5^ parameter sets we select those models that exhibit calcium transients with the experimentally plausible period, duration, and peak values (see Table A in S3 Appendix). Finally, we test sensitivity by varying each optimized parameter in each model for *±*1%, *±*5% and *±*10% and testing if the results remain close to experimental when running the model for 5 different seeds of the random number generator. To create an astrocyte population, five fitted parameters are randomly sampled from a Gaussian distribution. Mean, variance, and cutoff values of the Gaussian distribution are based on conclusions from sensitivity analysis. The selected parameter sets are described in Table C of S1 Appendix.

### Selecting neuron model parameters

Neurons are represented by the AdEx model [68, 84] as implemented in the NEST simulator. Excitatory and inhibitory neurons are respectively adapted from initial bursting and regular spiking neurons from Table 1 in [68]. The parameters are selected in such a way to ensure that the neurons respond to a SIC-mimicking current with a burst of spikes that fit the experimental data (details described in S3 Appendix; based on the SIC as illustrated in Fig 5 in [48]). The parameters of inhibitory neurons are further selected to produce a higher firing rate than the excitatory neurons, which is consistent with the literature (see Table B in S3 Appendix). Sensitivity to parameter change is tested by perturbing *V*_reset_ and *b*, two parameters that strongly affect spiking regime of AdEx neurons. To generate neuronal population, *b* and *V*_reset_ are drawn from Gaussian distribution with mean, variance, and cutoff values determined through sensitivity testing. The selected neuronal parameters are listed in Table C of S1 Appendix.

**Fig 5.**
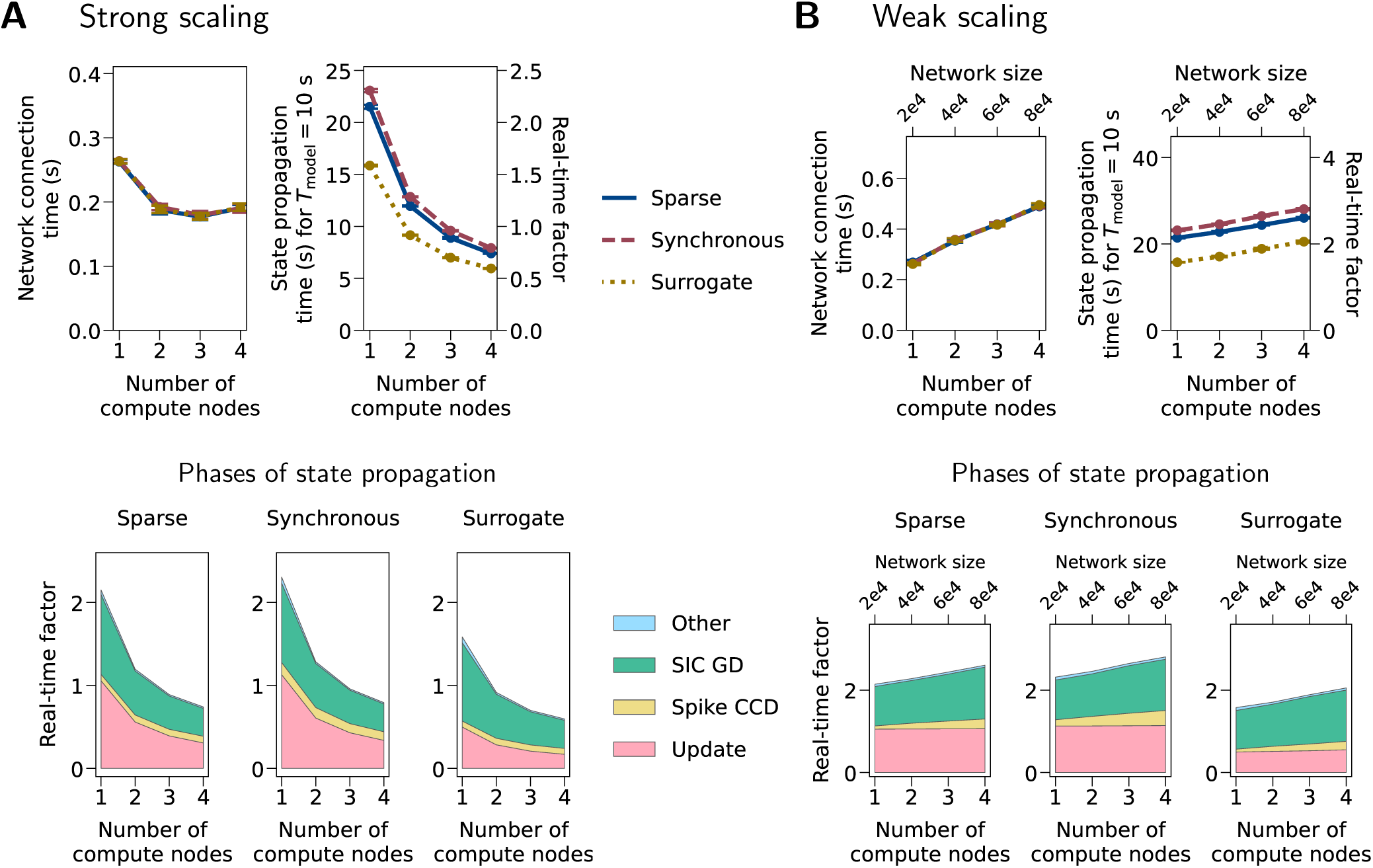
Benchmarks with neuron-astrocyte network models. (A) Strong scaling. In this type of benchmark, the network size stays constant (20,000 cells), but the number of compute nodes used is scaled. (B) Weak scaling. In this type of benchmark, the model size and the number of compute nodes are scaled together. The first row in (A) and (B) shows network connection time and state propagation time for a model time of *T*_model_ = 10 s and its real-time factor (*T*_wall_/*T*_model_). Error bars indicate standard deviations across nine simulations. In some cases the error bar is very small and hence not visible. The second row shows the phases of state propagation in terms of average real-time factor. “Update”: advancing the dynamical states in each cell and device; “Spike CCD”: collocation, communication, and delivery of spikes; “SIC GD”: gathering and delivery of SIC events; “Other”: the part of time that cannot be attributed to the three phases above. “Sparse”, “Synchronous”: models with sparse and synchronous activities, respectively. “Surrogate”: Model using astrocyte_surrogate (no astrocytic dynamics). For specifications of these models, see Tables A and B in S1 Appendix. For the dynamics of the “Sparse” and “Synchronous” models, see Fig 4.

### Selecting parameters of the neuronal network

We first select parameters that determine activity of the neuronal part only, the synaptic weights *w*_exc_, *w*_inh_ and rates of Poisson inputs to neurons *λ*_Poiss,E_, *λ*_Poiss,I_. The selected parameters ensure low-frequency asynchronous activity regime for the neuronal part of the model such that spiking frequency of excitatory and inhibitory cells falls within the experimentally plausible range of values (see values from the literature compiled in Table B of S3 Appendix). The neuronal spiking rates are set to a lower end of the biologically plausible intervals to accommodate for the additional increase in the spiking rate due to SIC inputs. The following parameter values are used in simulations unless otherwise stated: *λ*_Poiss,E_ = 2700 spikes*/*s, *λ*_Poiss,I_ = 2500 spikes*/*s, *w*_exc_ = 5 nS, and *w*_inh_ = *−*5 nS. The Poisson rate is equal for all neurons of the same type, the synaptic weights are identical for all synapses of the same type. Synaptic weights and delays are supplied to the TBP connection rule in NEST as part of the primary synapse specification, cf. Fig 3.

### Strength of neuron-astrocyte interactions, selection of *w*_pre to astro_ and *w*_astro to post_

We first determine the values for *w*_pre to astro_ sufficient to induce astrocytic calcium transients in the absence of Poisson input to astrocytes (*λ*_Poiss,A_ = 0) and in the absence of astrocyte feedback to neurons (*w*_astro to post_ = 0). Similarly, we determine the values for *w*_astro to post_ sufficient to induce SIC in the postsynaptic neurons in the absence of other (noise or synaptic) inputs. Next, we tune both parameters together to find the correct activity regime for each of the models shown in Results and S3 Appendix. In order to maintain dynamical regime of astrocytes in the presence of synaptic inputs, the rate of the astrocytic Poisson input is decreased to 70% of its value obtained from fitting single astrocytes. Weights and delays for the neuron-to-astrocyte connections are supplied to the TBP connection rule in NEST as part of the third in synapse specification, while weights for the astrocyte-to-neuron connection are supplied via the third out synapse specification.

### Probability of interaction between a synapse and an astrocyte

In order to test synchronization in neuronal groups resulting from neuron-astrocyte interaction, it is crucial to control the number of postsynaptic neurons interacting with the same astrocyte, and the number of astrocytes sending SIC to the same neuron. With our network parameters, these connected cell numbers per astrocyte or neuron are fixed across individuals if the block pool type is used, due to the nature of the block pool connectivity. In the random pool type, the number of postsynaptic neurons per astrocyte depends on *p*_third if primary_ and differs between astrocytes (see Figs 1 and 2 in S3 Appendix). The number of astrocytes that send SIC to the same neuron has an upper limit defined by *S*_pool_, but it is also determined by *p*_third if primary_. Thus, for block pools, the choice of *p*_third if primary_ is less important and we set it to be the same as the primary connection probability *p*_primary_ between pairs of neurons. However, for random pools the selected values for *p*_third if primary_ have to be much smaller in order to support both asynchronous activity and network bursting. A smaller probability of connection leads to fewer individual tripartite synapses which has to be compensated by an increase of *w*_pre to astro_ and *w*_astro to post_. Pool type and size and the probability of connecting to an astrocyte are passed to the TBP connection rule as part of the `third factor conn spec` specification.

### Hardware and software configurations for simulations

For the assessment of performance, the study characterizes an implementation of the astrocyte framework on the basis of NEST 3.8 on 128-core compute nodes of the supercomputer JURECA [85] at Jülich Research Centre (Rocky Linux 8, AMD EPYC 7742). Each compute node hosts two MPI processes and each benchmark consists of a total of nine simulations composed of three repetitions of three different seeds. The simulations reproducing experimental data employ NEST 3.8 on a personal computer (Debian Linux subsystem under Windows 10, Intel Core i7-8650U) and fit the astrocyte parameters to the values from the literature with NEST 3.6 running on 48-core compute nodes of a cluster (Debian 11, Intel Xeon E5-2680v3) at Jülich Research Centre. All simulations use the Runge-Kutta-Fehlberg-45 adaptive step size method, bounded from above by the overall simulation time grid of 0.01, 0.1, or 0.2 ms model time.

### Data analysis

#### Mean neuronal firing rate

The mean neuronal firing rates of the network models used in this study are calculated with spike counts during the sampled window of the simulations:

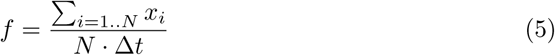

where *x_i_* is the spike count of neuron *i* during the sampled window, *N* is the number of neurons, and Δ*t* is the duration of the sampled window.

#### Pairwise spike count correlation as a measure of synchrony

Pairwise spike count correlation is used as a measure of overall similarity of spike trains in two neurons, reflecting their underlying synchrony. For the benchmark and the model based on the experimental literature, the same approach is used for the calculation of correlation coefficient, i.e., Pearson’s *r*, but different approaches are used for binning spikes due to different model activities. For the benchmark models (Fig 4), spike counts in 10 ms non-overlapping bins are obtained from *t* = 1 to *t* = 11 s of the simulation for all sampled neurons (*N* = 100), and pairwise spike count correlation is then calculated as:

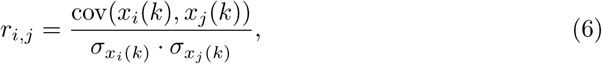

where *x_i_*(*k*)*, x_j_*(*k*) are spike counts of the *i*th and *j*th sampled neurons in the *k*th bin. cov(*x_i_*(*k*)*, x_j_*(*k*)) and *σ_xi_*_(*k*)_ and *σ_xj_* _(*k*)_ are the covariance and standard deviations of *x_i_*(*k*) and *x_j_*(*k*). The mean correlation *r* across all neuron pairs is then calculated as:

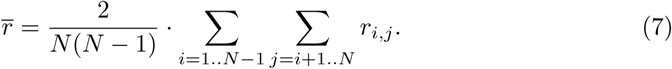

For the model based on experimental literature, spike trains of a neuron are binned using a sliding window of length 2 s which is shifted for 4 ms in each step. Binned spike trains for two neurons, *x_i_*(*t*) and *x_j_*(*t*), are used to compute the correlation coefficient given by Eq (6). Distribution of correlation coefficients is shown in Figs 9-12D.

#### Single-neuron burst detection

To test the impact of neuron-astrocyte interaction, we evaluate SIC-induced single-neuron bursts. A neuron burst can appear between SIC onset and 2 s (for excitatory neurons) or 400 ms (for inhibitory neurons) after the SIC offset. Spikes belong to the same burst if they are closer in time than 2 s for excitatory or 400 ms for inhibitory neurons. Details of neuron burst detection are given in S3 Appendix. Parameters of the burst detection algorithm are listed in Table C of S1 Appendix. From detected bursts we record burst onset, duration, frequency of bursts, and number of spikes per burst for each neuron. Burst onsets are used to evaluate SIC-induced synchronization in pairs of neurons, the results are reported in Figs 9–12.

#### Detection of astrocyte transients

Calcium transients are detected when calcium levels cross the same threshold used to trigger SIC current. Noise in calcium recordings might induce multiple threshold crossings around the transient onset and offset times. This is accounted for by merging crossings that are too close in time, as described in detailed transient detection algorithm in S3 Appendix. The algorithm parameters are also listed in Table C of S1 Appendix. Transient duration is measured as the interval between its onset and offset, while frequency is computed as the total number of transients divided by the span of time simulated. The latter we call model time *T*_model_ to contrast it from the wall-clock time *T*_wall_ required by a computer to carry out the simulation (Albers et al. 2022 [77]). Frequency and duration of calcium transients for all astrocytes in the model are shown in Figs 9–12A.

#### Distance of burst onsets as a measure of synchrony

Synchrony in pairs of spiking neurons is evaluated as a distance between their burst onsets in successive SIC-induced bursts. The metric is inspired by synchrony detection adopted in [48], and it also reflects phase synchrony in coupled spiking neurons. The metric is implemented as a Python function and supplied with the rest of the code.

## Results

### Benchmark results confirm efficiency of the implemented NEST support for astrocytes

In order to evaluate the performance of the implemented models and functions, we run benchmarks with neuron-astrocyte network models as described in Methods. Following the previous approach [77], we use NEST and beNNch to measure the time spent on network creation, network connection, and state propagation. The state propagation consists of multiple phases, labeled as follows: “Update” is the time needed to advance the dynamical states of all cells and devices. “Spike CCD” and “SIC GD” is the time required to exchange of information in parallel computing: “Spike CCD” is the time for collocation, communication, and delivery of spikes, and “SIC GD” the time for gathering and delivery of the SIC events. “Other” stands for the part of time that cannot be attributed to the three specific phases.

Fig 5 shows network connection and state update times for the first set of benchmarks; creating the neurons always takes less than 50 ms and this time is therefore not shown. In the strong-scaling benchmarks, the network connection time reaches a minimum for three compute nodes and then increases, while the state propagation time always decreases with the scale. In the weak-scaling benchmarks, both network connection time and state propagation time increase with the scale, except for the “Update” phase which advances the dynamical states in individual cells and hence can be highly parallelized. All models spend less than 1 s on network construction (creation + connection), and the real-time factor (*T*_wall_/*T*_model_) for state propagation is always less than three.

Consistent with Fig 4, the “Synchronous” model always produces many more spikes than the “Sparse” model. Nevertheless, this only slightly affects the simulation time. The mean neuronal firing rate of the “Synchronous” model is more than twice that of the “Sparse” model (4.76 versus 11.95 spikes/s for 20,000 cells; mean of nine simulations). However, in both strong and weak-scaling benchmarks, the state propagation times are similar, with a difference of 7.0% to 8.6% compared to the “Sparse” model (Fig 5). This limited effect of higher firing rate on state propagation time reflects the small differences in each of the phases (Fig 5). This shows a good efficiency in the communication and processing of spikes between and within the cells, and the fact that sic_connections send continuous SIC signals regardless of the level of model activity.

The “Surrogate” model has a similar mean neuronal firing rate as the “Sparse” model (4.64 versus 4.76 spikes/s for 20,000 cells) but spends less time on state propagation, which is mainly due to the shorter “Update” phase (Fig 5). Since astrocyte_surrogate does not have astrocytic dynamics and sends predefined SICs instead, in the “Surrogate” model the time spent on the “Update” phase is mostly for the update of states in the neurons. The “Sparse” model spends 80.9% to 112.1% more time on “Update”, compared to the “Surrogate” model (Fig 5). This difference approximately represents the additional computational demands in the “Update” phase due to the astrocytic dynamics. On the other hand, since the connectivity function does not distinguish astrocyte_lr_1994 and astrocyte_surrogate, the network connection time of the “Surrogate” model are similar to the “Sparse” model.

The benchmark results with four primary (neuron-to-neuron) connectivity rules are shown in Fig 6. The four models have a similar performance in all aspects, except that the fixed out-degree rule shows a longer network connection time than the other three because it cannot be parallelized. Note that the “Bernoulli” model is the same as the “Sparse” model in Fig 5, where the pairwise Bernoulli rule is used for the primary connections, and the other three models have the same expected numbers of primary and third-factor connections as the “Bernoulli” model.

**Fig 6.**
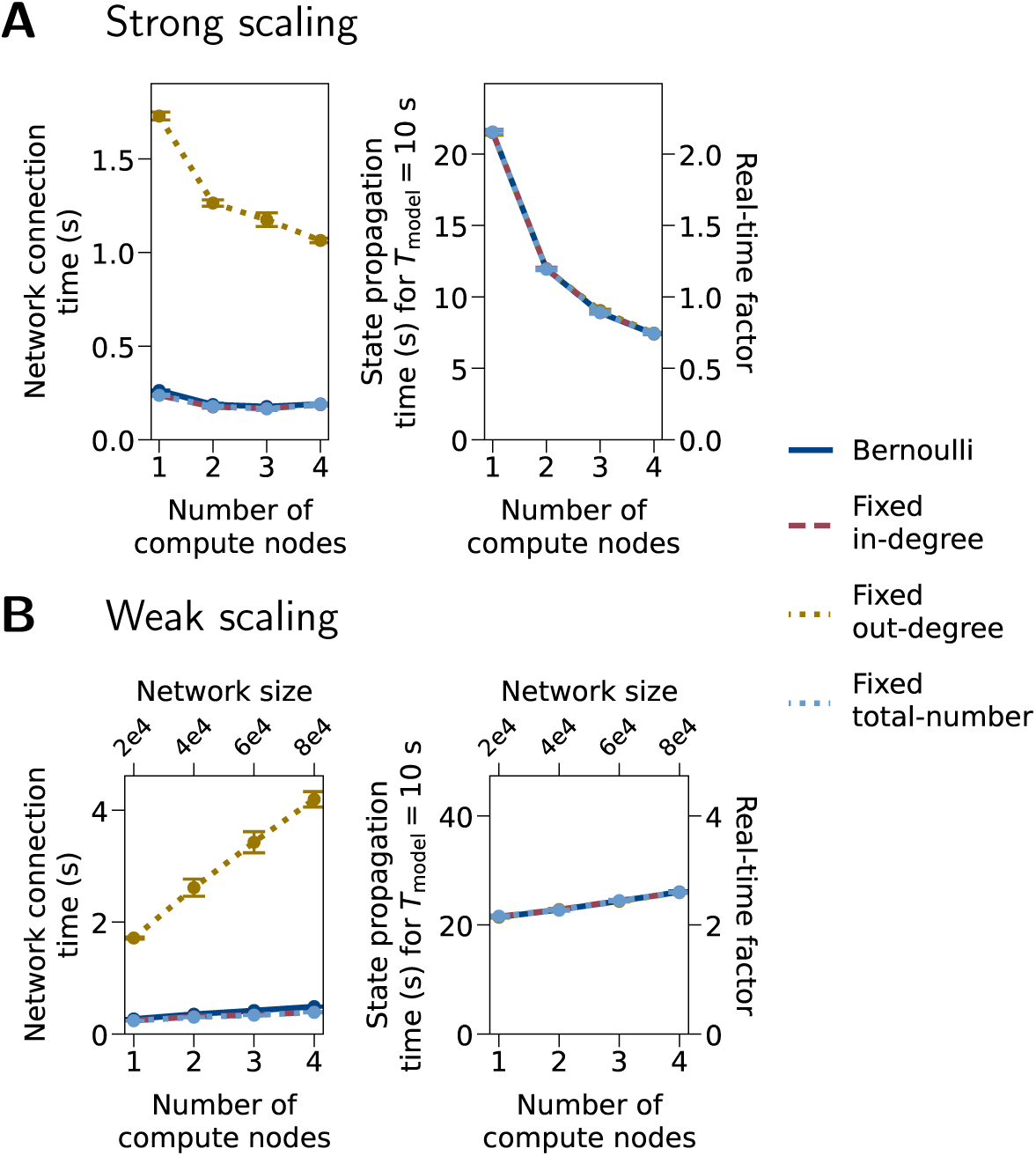
Benchmarks with different primary (neuron-to-neuron) connectivity rules. (A) Strong scaling. The network size is 20,000 cells. (B) Weak scaling. The four tested connectivity rules are pairwise Bernoulli, fixed in-degree, fixed out-degree, and fixed total number. For specifications of these rules, see [4]. Model parameters other than the connectivity rule are the same as the “Sparse” model in Fig 5. Notations and benchmark configurations are the same as in Fig 5.

Weak-scaling benchmark results for different astrocyte pool sizes are shown in Fig 7. While network connection time is only mildly affected by pool size, pool sizes of 100 and larger lead to increased state propagation times compared to pool size 10. For four compute nodes and a pool size of 100, the increase is 61.5%, for a pool size of 10,000 it is 65.6%. This increase is driven by increased spike and SIC communication times as well as an increased “Other” time (Fig 7, second row). This difference in scaling behavior appears to be due to different communication requirements. If the astrocyte pool size is well below the average number of incoming astrocytic connections per target neuron (astrocytic in-degree, 400 in our models), the number of target neurons to which any astrocyte connects is close to the pool size; for large pools, on the other hand, it is determined by the astrocytic in-degree. In particular, for a pool size of 10, each astrocyte connects to 10 target neurons on average. In this case and when using four compute nodes with eight MPI processes in total, *I*_SIC_ needs to be transmitted to all MPI processes only for fewer than 10% of all astrocytes. For a pool size of 100 and larger, on the other hand, *I*_SIC_ from all astrocytes must be communicated to all MPI processes, requiring higher bandwidths. The data for a pool size of 10 are the same as the “Sparse” model in Fig 5.

**Fig 7.**
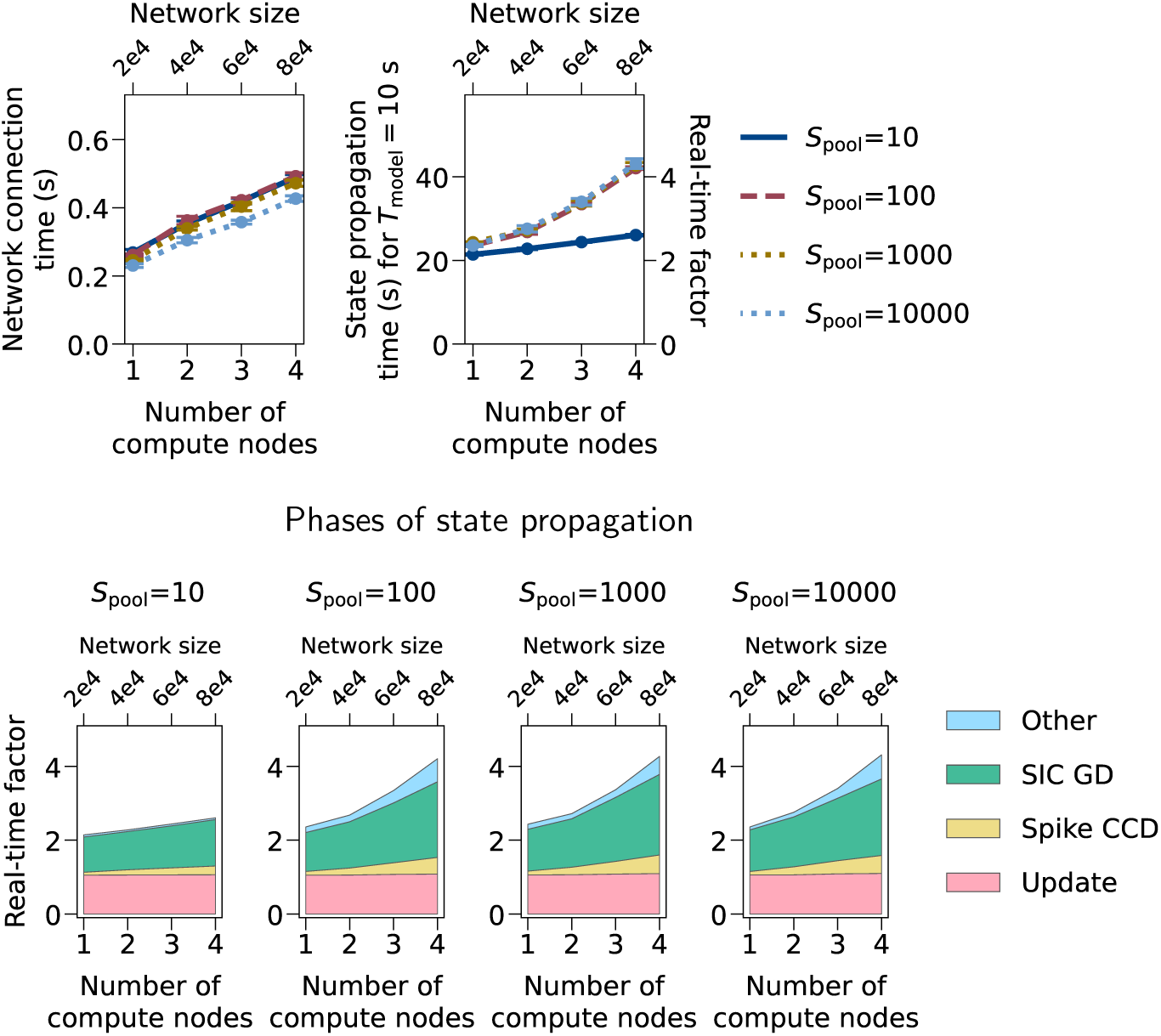
Weak-scaling benchmarks with different astrocyte pool sizes. Weak-scaling benchmark results with astrocyte pool sizes per neuron (*S*_pool_) from 10 to 10,000. Model parameters other than *S*_pool_ are the same as the “Sparse” model in Fig 5 and the “Bernoulli” model in Fig 6. Notations and benchmark configurations are the same as in Figs 5 and 6.

To demonstrate the performance of our approach for very large network models, we repeat the benchmarks for tenfold larger networks, i.e., for 200,000 instead of 20,000 cells per compute node. The largest network simulated has one million cells in total. Fig 8 shows weak scaling behavior as network size and compute resources grow by a factor of 40: While network connection time increases by a factor of 6.8, it remains much shorter than simulation time (6.2 s vs. 293 s for *S*_pool_ = 10 on 5 compute nodes, 10 s simulated time) and is thus unproblematic in practice. For small pools (*S*_pool_ = 10), state propagation (simulation) time increases only by a factor of 1.8, i.e, scaling is good if not perfect. Furthermore, the real-time factor is about ten times larger than for the smaller models, which indicates that the simulation algorithm scales linearly in the number of cells per compute core. For larger pools (*S*_pool_ = 1,000), simulation times increases markedly for more than two compute nodes (factor 3.2 from ^1^ to 5 nodes) in the same way and for the same reasons as for the smaller networks.

**Fig 8.**
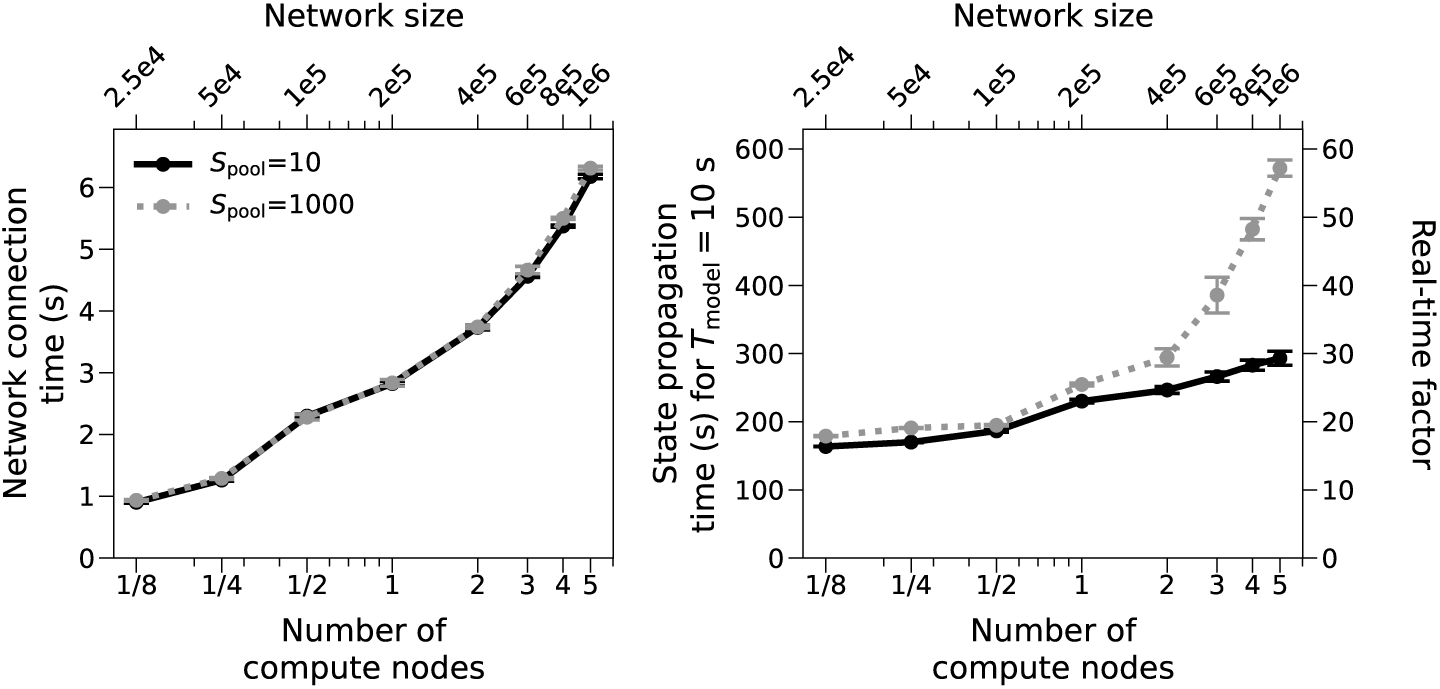
Weak-scaling benchmarks for very large models. Network building and simulation times for the same models and parameters as the “fixed in-degree” model in Fig 6 but with 200,000 instead of 20,000 cells per compute node and for *S*_pool_ = 10 and *S*_pool_ = 1,000. Fractions in the x-axis indicate that only a part of a compute node is used (16, 32 or 64 out of 128 physical cores).

### Use case: Astrocytes promote neuronal synchronization *in silico* through neuron-astrocyte interaction

In this use case, we demonstrate how the newly developed NEST support for astrocytes can complement a selected experimental study described in [48, 53]. Recording from rat cortical, hippocampal, and thalamic slices, the authors of [48, 53] provided evidence for synaptic glutamate release, astrocyte activation, SIC generation mediated by extrasynaptic NMDARs, and subsequent slow calcium responses in neurons. They demonstrate that SICs induce synchrony in groups of neurons and hypothesize that this synchronization results from neuron-astrocyte interactions. In what follows, we explore synchrony in the computational model of neuron-astrocyte network.

First, we reproduce the duration and frequency of spontaneous calcium transients in astrocytes reported in experimental studies. The frequency of SIC occurrences (and calcium transients that induce it) is estimated to be 1.26 *±* 0.2 SIC/min in the slices pre-exposed to glutamate and 0.07 *±* 0.01 SIC/min in control [48]. Other studies, that examine calcium dynamics in astrocytes but do not consider SIC, report frequency of calcium transients to be 1.2 transients/min [59], 4 transients/min [61], and 0.5 transients/min [60]. Duration of calcium transients is found to be between 1–20 s [61] and 9–18 s [60]. These values collected from experimental studies are systematically listed in S3 Appendix. We fit a model of an isolated astrocyte, and specifically the parameters [Ca^2+^]_tot_, [IP_3_]_0_, Δ_IP3_, and *τ*_IP3_ as well as the rate of the input Poisson noise *λ*_Poiss,A_ to reproduce these values (see Methods and S3 Appendix). The selected rate of the Poisson noise, together with other parameters, ensures that astrocytes exhibit biologically plausible dynamics of calcium transients. In addition, Gaussian noise is added to the calcium equation of each astrocyte to model random fluctuations typically seen in calcium traces. Noise sources are stationary and independent for each cell to guarantee uncorrelated astrocyte activity in the absence of interaction with neurons. Thus any emergent synchrony in the model results from neuron-astrocyte interactions. Finally, the results of this study do not depend on specific astrocyte or noise parameters. Any model that can reproduce the same spontaneous calcium dynamics in astrocytes could lead to the same results. All model parameters are listed in Table C of S1 Appendix and also in a JSON file supplied with the code for model implementation. The obtained model generates transients with frequency of about 0.5–1.5 per minute and duration of between 1–5 s as shown in Fig 9A.

**Fig 9.**
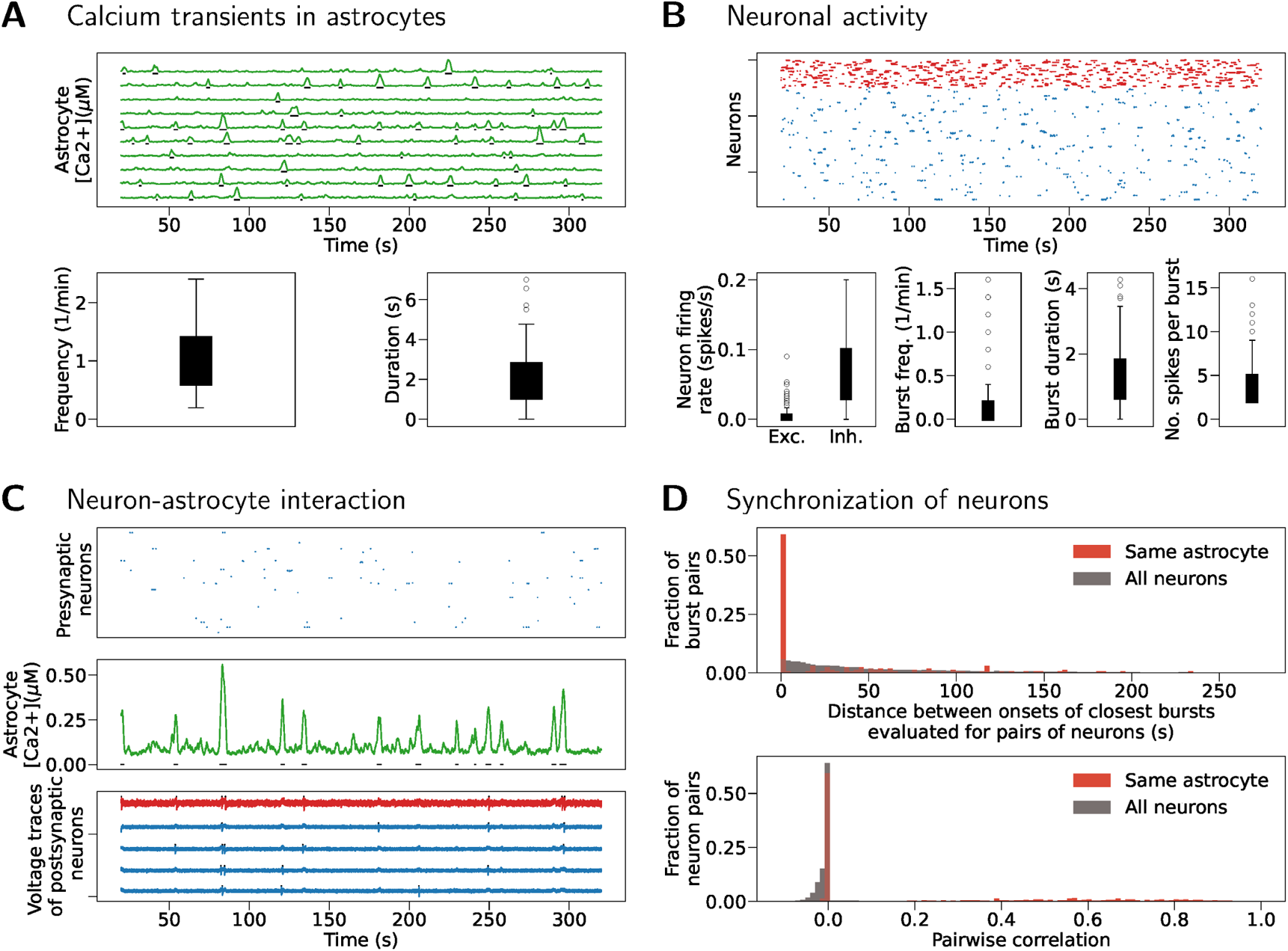
Reproducing experiment with TTX blocking - neurons do not spike without SIC input and all synaptic weights are set to zero except *w*_astro to post_. (A) Top: Spontaneous calcium transients in ten randomly selected astrocytes. Bottom: Frequency and duration of calcium transients across all astrocytes in the model. (B) Top: Rasterplot showing activity of all excitatory (blue) and inhibitory (red) neurons in the model. Bottom left: Spiking frequency for excitatory (Exc.) and inhibitory (Inh.) neurons. Bottom right: Single neuron burst frequency, duration and number of spikes per burst. Bursts are evoked solely by SIC inputs in this simulation. (C) Single astrocyte (middle panel) is depicted together with its 5 postsynaptic neurons (bottom panel, blue voltage traces - excitatory neurons, red traces - inhibitory neurons) and all of its (excitatory) presynaptic neurons (top panel). Synchronization induced by calcium transients and resulting SIC can be seen in voltage traces of postsynaptic neurons. (D) Top: Neuronal synchrony is evaluated as a difference in onsets of successive burst events. Bottom: Synchrony evaluated as pairwise correlation between binned spike trains.

Next, we focus on the experiment from [48] in which extracellular glutamate was increased through pre-exposure to glutamate while neuronal spiking was pharmacologically blocked by TTX. We construct a population of *N*_A_ = 100 astrocytes, *N*_E_ = 400 excitatory, and *N*_I_ = 100 inhibitory neurons. Astrocytes have only Poisson input mimicking the processing of extracellular glutamate, while neurons receive weak Gaussian noise and cannot spike without SIC input from astrocytes to the postsynaptic neuron. In response to a SIC input, neurons develop a short burst containing normally 2–4 (but up to 14) spikes and lasting for about 1 s. In this model system, this burst is interpreted as a correlate of a slow calcium response measured in [48], rather than a sequence of spikes (which cannot occur in experiments with slices exposed to TTX).

Neurons are randomly connected with probability *p*_primary_ = 0.2, however the synaptic weights are set to zero to account for the blocking of spiking via TTX. The neuron-astrocyte interaction is structured by block pools of size one, meaning that a neuron can receive SIC from only one astrocyte. This choice of parameters results in 100 non-overlapping neuronal groups, each consisting of 4 excitatory and 1 inhibitory neuron that can receive SIC from the same astrocyte. Fig 9C illustrates calcium activity of a single astrocyte (middle sub-panel) induced by Poisson input. Activity of presynaptic neurons is shown as a rasterplot (top sub-panel). However due to synaptic weights set to 0, this activity does not affect astrocytes. Synchronization induced by SIC input between an astrocyte and five neurons interacting with this astrocyte is illustrated in the bottom of Fig 9C. Fig 9D demonstrates increased synchrony in groups of neurons connected to the same astrocyte compared to the overall level of synchronization between neurons in the model. Synchrony is evaluated in two ways, either as a distance between onset of successive single-neuron bursts in pairs of neurons (upper panel) or as pairwise correlation (lower panel), for details see Methods and S3 Appendix. Bursts are well aligned in neurons that interact with the same astrocyte (distribution is collapsed to values very close to zero). Correlation coefficients are very small in general due to sparse activity in the model. Synchronization evaluated between neurons that receive inputs from the same astrocyte is significantly higher compared to synchronization evaluated across all pairs of neurons in the model (Kolmogorov-Smirnov test, *p <* 0.0001 for comparison of burst onset times, *p <* 0.0001 for comparison of pairwise correlations, notice a number of correlation coefficients distributed between 0.2–0.8); values after Bonferroni correction done for all tests in this section and all related tests in S3 Appendix).

The same result is reproduced in a model with spiking neurons and nonzero synaptic weights (Fig 10). The network has low-frequency asynchronous activity, where excitatory neurons spike at a frequency of about 0.1 spikes/s while inhibitory neurons spike at a frequency of about 2 spikes/s. Single-cell bursts resulting from SIC inputs are visible in the rasterplots of (sparser) excitatory activity in Fig 10B. Same as before, Fig 10C illustrates synchronization between an astrocyte and its postsynaptic neurons at each astrocytic calcium transient. The presynaptic activity, shown by the top sub-panel, contributes to astrocyte inputs in this case. Fig 10D shows increased synchronization (both measured as distance between burst onsets, and as pairwise correlation) between neurons connected to the same astrocyte (Kolmogorov-Smirnov test, *p <* 0.0001 for burst onset times, and *p <* 0.0001 for pairwise correlation; values after Bonferroni correction). Reproducibility of this result is confirmed by running the same simulation for different seed of the random number generator and obtaining similar results (see Fig 4 in S3 Appendix).

**Fig 10.**
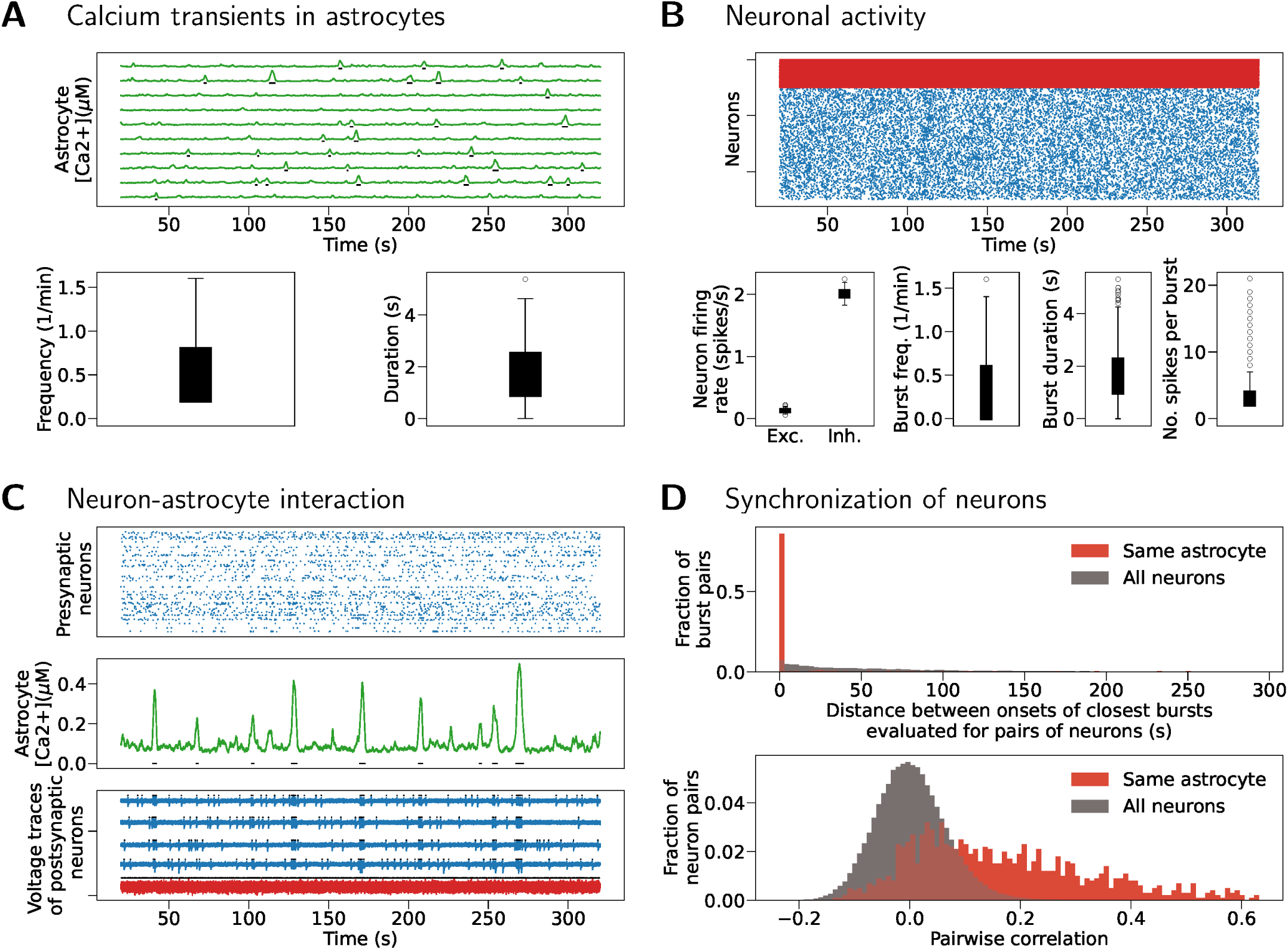
Synchronization in neuron-astrocyte networks with block pools and asynchronous activity regime. (A) Top: Spontaneous calcium transients in ten randomly selected astrocytes. Bottom: Frequency and duration of calcium transients across all astrocytes in the model. (B) Top: Rasterplot showing activity of all excitatory (blue) and inhibitory (red) neurons in the model. Bottom left: Spiking frequency for excitatory (Exc.) and inhibitory (Inh.) neurons. Bottom right: Single neuron burst frequency, duration and number of spikes per burst. (C) Single astrocyte (middle panel) with its five postsynaptic neurons (bottom panel) and all presynaptic neurons (rasterplot, top panel). (D) Top: Neuronal synchrony is evaluated as a difference in onsets of successive burst events. Bottom: Synchrony evaluated as pairwise correlation between binned spike trains.

Increasing the overall input to an astrocyte by increasing the weight *w*_pre to astro_ from 0.2 to 0.31 results in emergence of network-wide bursts lasting on average 5 s, as shown in Fig 11. Although the global synchronization increases in this activity regime, synchronization between neurons interacting with the same astrocyte still exceeds synchronization computed for all pairs of neurons in the model (Kolmogorov-Smirnov test, *p <* 0.0001 for burst onset times, *p <* 0.0001 for pairwise correlation; values after Bonferroni correction).

**Fig 11.**
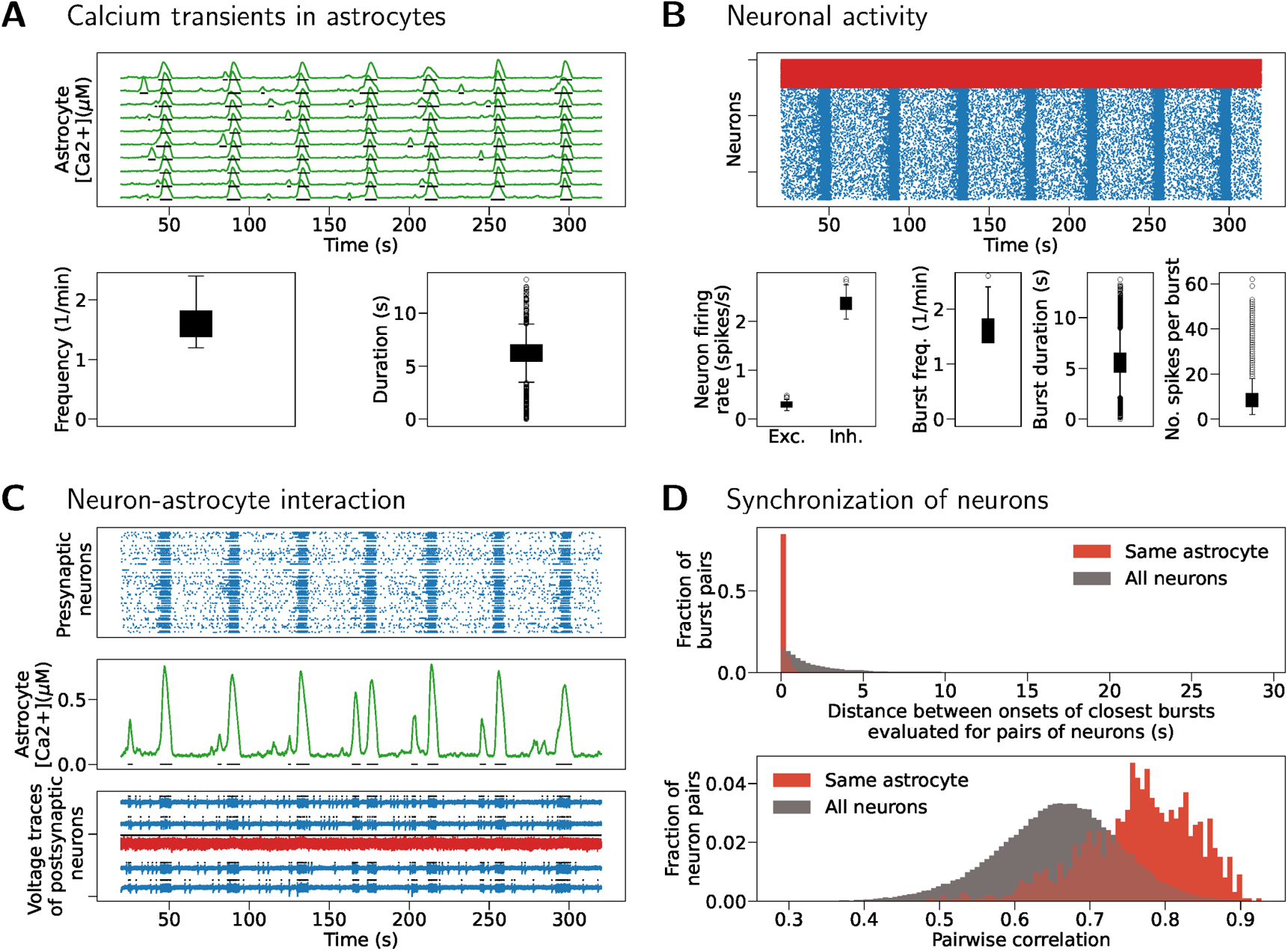
Synchronization in neuron-astrocyte networks with block pools and global network bursting activity. (A) Top: Spontaneous calcium transients in ten randomly selected astrocytes. Bottom: Frequency and duration of calcium transients for all astrocytes in the model. (B) Top: Rasterplot showing activity of all excitatory (blue) and inhibitory (red) neurons in the model. Bottom left: Spiking frequency for excitatory (Exc.) and inhibitory (Inh.) neurons. Bottom right: Single neuron burst frequency, duration and number of spikes per burst. (C) Single astrocyte (middle panel), its five postsynaptic neurons (bottom panel) and its presynaptic neurons (top panel) all follow global synchronous activity. (D) Top: Neuronal synchrony is evaluated as a difference in onsets of successive burst events. Bottom: Synchrony evaluated as pairwise correlation between binned spike trains.

So far, neurons form well separated groups, each group receiving inputs from only one astrocyte. The experimental study inspiring this model [48] shows that neurons can belong to more than one synchronized group and hypothesizes that structure of neuron-astrocyte interactions might create such groups. Using the support for the TBP connection rule implemented in NEST as part of this work, we construct a network model with tripartite connectivity using randomized astrocyte pools and evaluate the model dynamics as shown in Fig 12. The maximal number of astrocytes is determined by the model parameter *S*_pool_, which is set to 5 here. The connectivity depends on two probabilities, the probability to establish a neuron-neuron synapse, *p*_primary_ set to 0.2 here, and a probability to connect that synapse with an astrocyte, *p*_third if primary_ set to 0.03 here (but see the same result reproduced with a bigger probability value, *p*_third if primary_ = 0.05, in Fig 5 of S3 Appendix). The exact number of neuron and astrocyte inputs in this model is shown in Fig 2 of S3 Appendix. Due to overlap of astrocytic domains, synchronization within neuronal groups is established faster and it is easier to reach global bursting regime. Still, asynchronous low-frequency spiking regime is achievable also in this model, and the synchronization between neurons receiving SIC from the same astrocyte remains higher than the overall synchronization in the neuronal network (Kolmogorov-Smirnov test, *p <* 0.0001 for comparison of burst onsets, *p <* 0.0001 for comparison of pairwise correlations; values after Bonferroni correction). Increased synchronization between neurons interacting with the same astrocyte holds in the model with random pools which exhibits network bursting regime, as shown in Fig 6 of S3 Appendix. Similarly as before, network bursting regime is achieved by increasing the amount of glutamatergic inputs to astrocytes, i.e. by increasing the weight coefficient *w*_pre to astro_ from 1 to 1.5.

In order to support the here presented results, we provide several additional tests in S3 Appendix. Synaptic delay impacts the speed of neuron-neuron and neuron-astrocyte exchange and can affect the global activity regime. The simulations shown in Fig 10 (block pools, asynchronous network activity) and in Fig 6 of S3 Appendix (random pools, network bursting) are repeated for a synaptic delay ten times smaller (0.1 ms vs 1 ms) in Figs 7 and 8 of S3 Appendix, respectively. The simulations confirm robustness of our results against change in synaptic delay. We also verify that the results presented here indeed show the steady state of the system. The simulations in Fig 11 and in Fig 6 of S3 Appendix are repeated for much longer *T*_model_ in Figs 9 and 10 of S3 Appendix, respectively. The models are first simulated for 15 min, and then for additional 5 min (as opposed to 20 s and 5 min used in other figures). Analysis is done for the last 5 min of simulations and the results confirm previous findings. Finally, we test the impact of numerical integration method in three models (block pools and asynchronous, block pools and bursting, random pools and asynchronous) by changing the maximal integration step from the default 0.1 ms to 0.01 ms and 0.2 ms, and repeating the simulations. We confirm robustness of results with respect to perturbation of this parameter, which is illustrated by three examples in Figs 11-13 of S3 Appendix.

In summary, we show the emergence of synchronization in small groups of neurons as a result of SIC inputs arriving from the shared astrocyte partner. Our *in silico* results support the findings and test hypotheses set in [48, 53]. Furthermore, the results suggest an important role of astrocytes in shaping the global neural activity states through local neuron-astrocyte interactions.

## Discussion

In this study, we introduce conceptual advances in modeling and simulating neuron-astrocyte circuits. Our novel connectivity framework allows flexible integration of astrocytic influence while ensuring scalability for very large-scale simulations. We validate its efficiency through benchmarking with models containing up to one million cells and provide a formalized description of neuron-astrocyte interactions. A key contribution is our exploration of how astrocytes promote self-organization of brain circuits by contributing to the emergence of local synchrony in neuronal groups, moving beyond previous studies focused on global network effects [35, 52, 86, 87]. By incorporating structured connectivity and experimentally grounded hypotheses, our study offers new insights into the role of astrocytes in network dynamics.

The novel theoretical connectivity concepts and technology described here provide the foundation for a comprehensive and efficient modeling and simulation platform for neural systems composed of both neuronal and non-neuronal cells. We present a general concept for specifying tripartite connectivity in terms of primary and third-factor connection rules and provide a reference implementation suitable for large-scale distributed simulation. Following this concept, interactions with third factors such as astrocytes can now be attached to neuronal networks based on a wide range of connectivity rules [4].

### Advances in theoretical frameworks and simulation technology

We introduce a schema for the declarative specification of tripartite connectivity and provide a parallelized reference implementation to instantiate this connectivity in NEST. Our approach is based on the tripartite synapse hypothesis [13, 88] and the experimental observation that astrocytes form spatial domains with little overlap [5–10]. Following the tripartite synapse hypothesis, our tripartite connectivity rule requires that third-factor cells such as astrocytes are attached to pairs of pre- and post-synaptic neurons by co-occurring presynaptic-astrocyte and astrocyte-postsynaptic connections. Spatial domains of limited overlap are represented as astrocyte pools in our rule (see Concepts for tripartite connectivity). We support random and tiled (blockwise) pools of flexible size. We thus enable biologically grounded studies of tripartite interactions with a parameterized variability in connectivity as illustrated by our use case (see Use case for the technology). Future studies can set more plausible constraints as more detailed experimental data from specific brain regions becomes available.

For astrocytes, we describe a well-known Li-Rinzel model of calcium dynamics [44], based on an earlier model for IP_3_ kinetics and calcium exchange between ER and cytosol [43]. In our implementation, the astrocyte can be represented either as a single-compartment cell or by several disconnected compartments. This description of astrocytes is compatible with point-neuron models typically used in large-scale simulations of brain circuits. A point-neuron model that we consider is the standard AdEx spiking neuron [67, 68, 84] extended to receive currents induced by astrocyte activity. It should be noted that astrocytes *in vivo* have complex, highly compartmentalized morphology with non-trivial interaction between compartments. Multi-compartmental astrocytes can be integrated with multi-compartmental neurons into detailed mechanistic models of brain circuits; this relevant approach is, however, outside the scope of this study and the NEST simulator. Like other studies, we implement the astrocyte model with several constraints such as assumptions about the fixed total calcium in ER and cytosol, and the fixed ER to cytosol volume ratio.

Additionally, we allow random calcium flux across membranes by adding a noise input to the calcium equation in Eq (1). Relaxing these model constraints and implementing additional astrocyte models can be considered in future extensions of this work. A model that represents more realistic aspects of astrocyte biology can potentially account for some of the variability typically seen in calcium recordings.

Astrocyte–to–neuron interaction in our implementation follows a model first proposed by Nadkarni and Jung in 2003 [45] and later frequently used in a number of computational studies [33, 35]. This interaction assumes that astrocytes induce slow inward currents (SICs) [45, 48] to proximal postsynaptic neurons in response to the presynaptic release of glutamate. The presynaptic release of glutamate evokes a neuron–to–astrocyte signal, which is modeled as a constant instantaneous increment in IP_3_ concentration. This approximation is computationally efficient and consistent with large-scale spiking neural models even though it does not capture the complexity of involved biological mechanisms. More detailed models of IP_3_ concentration or accounting for volume transmission similar to Magloire et al. [52] can be integrated into this framework at a later stage.

The technology for simulating continuously coupled equations in a distributed computing setting as required for large-scale neuronal systems is available [2]. This enables, for example, the inclusion of gap junctions in a network model. However, due to our phenomenological description of the SIC mechanism, the full waveform relaxation framework is not required. The assumption of a finite delay in the interaction enables the use of the same optimization as for the spiking interaction: communication is only required in intervals of the minimal delay in the system [74]. An approach based on the molecular mechanisms of systems biology could unify the interaction types including spikes, gap junctions, and SIC currents, but for the price of higher computational costs.

The astrocyte and neuron models of the present study reflect common choices across the literature [32, 35], and also correspond to the models that are available in common simulation tools [40, 42, 86, 89]. As such, our choice addresses the needs of the community and facilitates model development across existing tools. We further support this goal by providing a systematic comparison of variable and parameter names used in the literature, in this article, and in the user-level documentation of our reference implementation (see S2 Appendix). Reproducibility and model development are further enhanced by a systematic description of model dynamics and model parameters in a tabular format in S1 Appendix [37]. We adapt the format originally developed for neuronal networks to include astrocytes and neuron-astrocyte interactions. We believe that this formalism should become standard for systematic and rigorous description of neuron-astrocyte models in the literature.

While here we opt for well-known models from the literature for demonstration, the framework for astrocyte modeling can be extended to other experimentally confirmed mechanisms including more complex models for astrocytic calcium dynamics, neuron-astrocyte interaction through other neurotransmitters (e.g. GABA), the release of other molecules including D-serine and ATP, and further mechanisms. Furthermore, the impact of non-stationary extracellular neurotransmitter concentrations can be accounted for in future versions of the framework as proposed in [52]. In this study, we consider only the basic AdEx neuron model as, for example, available in the NEST simulation code, but other point neuron models can be similarly generalized to receive astrocyte inputs. Connecting cells according to their explicitly specified spatial location is another potential future development that would benefit from more comprehensive morphometric data and future neuroanatomical studies.

### Benchmarking the technology

The benchmark results demonstrate the efficiency of our reference implementation when using parallel computing resources. The strong-scaling benchmark examines the simulation performance with fixed model size and up-scaled computing resources.

Parallelization decreases network connection time and state propagation time. The weak-scaling benchmark examines the performance with coupled up-scaling of model size and computing resources. In the weak scaling the time spent for phases of the simulation that require communications between parallel processes (i.e., network connection, “Spike CCD”, “SIC GD”) increases with the scale. Nevertheless, both the strong and weak scaling benchmarks show a good performance in network construction time and state propagation time. With one compute node, the real-time factor for state propagation is close to two in all tested benchmark models. The real-time factor is further reduced by strong scaling, and in weak scaling it remains below three for most of the tested benchmark models (Figs 5, 6) and below five in benchmarks with large astrocyte pool size (Fig 7).

In the first set of benchmarks, three models are used to evaluate how overall model activity and astrocytic dynamics affect simulation performance (Fig 5). Parameters of the “Sparse” and “Synchronous” models are chosen to yield sparse and synchronous neuronal activity, respectively. Even though the difference in neuronal activity is large, the overall simulation times of the “Sparse” and “Synchronous” models are similar, suggesting that the performance is only slightly affected by the level of activity in the tested models. In the “Surrogate” model, the astrocytes are replaced by surrogate cells, which do not have astrocytic dynamics but send predefined SIC currents to their target neurons. The comparison between the “Sparse” and “Surrogate” models exposes the additional amount of time spent on astrocytic dynamics. The result suggests that, in a network model where the number of astrocytes equals the number of neurons, the astrocytes are equally expensive as the neurons (Fig 5).

The second set of benchmarks shows that models with different primary (neuron-to-neuron) connectivity rules have very similar performance in general, except for the non-parallelizable fixed out-degree rule (Fig 6). The third set of benchmarks shows the effects of astrocyte pool size per neuron, i.e., the number of astrocytes that each neuron can receive inputs from. The benchmark data suggest that large astrocyte pool size increases the workload in communications for spikes and SIC currents (Fig 7). The fourth set of benchmarks tests very large model sizes and shows that the performance is preserved with sizes up to one million cells (Fig 8).

Overall, the benchmark results show good scaling performance and limited effects of model activity or network-level parameters on the performance. This suggests that our technology is capable of simulating diverse large-scale neuron-astrocyte network models.

### Use case for the technology

As a use case, we reproduce and extend the selected experimental findings [48] by implementing, fitting to the available data from the literature, and simulating a set of computational models of neuron-astrocyte networks. The study by Pirttimaki et al. [48] focuses on the biophysical origin of SIC current and its impact on neuronal activity.

The phenomenon is observed in spontaneously active circuits and in circuits where spontaneous activity is enhanced by extracellular glutamate. The study demonstrates how astrocytes support synchronization in neuronal groups, possibly in neurons interacting with the same astrocyte, in cortical, hippocampal, and thalamic slices [53, 58]. We construct a computational model that closely approximates the experimental setup of [48], conduct equivalent *in silico* experiments, and demonstrate that astrocytes promote synchronization in neuronal groups across various connectivity schemes and under different global activity regimes.

To reproduce realistic spontaneous activity in astrocytes, we conduct an extensive survey of experimental evidence from the literature, summarized in S3 Appendix, and fit a model for a single astrocyte. Spontaneous calcium activity has been observed (using confocal microscopes) in astrocytes in hippocampal slices, in soma, processes, and microdomain areas in wild-type mice [59]. Typical spontaneous calcium event statistics calculated from somatic areas of astrocytes reveal a frequency of about 1.2 transients per minute per region of interest (ROI), an amplitude of about 3 dF/F, and a duration of about 3–10 s although shorter and longer spontaneous calcium events can occur as well [59]. Similar values have been reported in [61] when using organotypic slices prepared from mice hippocampus and imaged by confocal microscopy. 0.17 transients per minute, with an average duration of about 10 s, have been shown in astrocytes using hippocampal slices [90]. Another study reveals that the overall pattern of calcium transients within astrocytes are similar between hippocampal astrocytes in brain slices and cortical astrocytes *in vivo* for wild type and IP_3_R2^-/-^ mice, implying that the measured calcium transients are not the consequence of the method employed to study them [59]. The exact statistics of calcium transients may vary depending on the brain area and developmental age [91], but not all studies report differences with age [90]. *In vivo*, spontaneous transients have had a frequency of 0.4 transients per minute per ROI in the soma and 0.64 transients per minute per ROI in the processes [92]. The amplitude of the transients varies from 0.1 dF/F in soma to about 0.35 dF/F in processes, and the average duration of the transients is about 10 s in both soma and processes [92]. Finally, Pirttimaki et al. [48] do not measure calcium dynamics directly, but the reported frequency of SIC currents is about 1.26 SIC/min and their duration is about 900 ms on average.

In our simulations, calcium transients appear with a frequency of about 0.1–1.5 transients per minute, with an average of 0.5 transients per minute in the asynchronous regime and 1.5 transients per minute in the bursting regime. These transients last for about 1–10 s and increase from the baseline level below the concentration of 0.1 µM to at most 0.7 µM. All these values correspond well to the ranges seen across the experimental literature. However, the experimental values vary greatly across studies, experimental preparations, types of calcium transients, and astrocyte domains (soma vs processes). Systematic quantification of this diversity is lacking, and its functional role is unclear. Data-driven computational modeling of astrocytes would greatly benefit from systematic recordings of calcium transients and collecting sufficiently big and well-documented benchmark data sets. Better categorization of calcium transients, their dynamic properties, and functional roles is necessary. Finally, understanding calcium dynamics across astrocyte compartments is critical not only for advancing computational models of single astrocytes and neuron-astrocyte networks, but also for understanding the role of astrocytes in brain systems in general.

We conduct an extensive parameter search to establish an astrocyte model that reproduces experimentally observed spontaneous calcium dynamics. This way we (1) ensure the correct dynamics of a single astrocyte, (2) decouple the parameter choice for the astrocyte and neuronal part of the model, and (3) constrain the parameter space of the entire network model. With this approach, we ensure that astrocytes do not synchronize in the absence of neuronal activity; this assumption allows to focus on the selected mechanism of neuron-astrocyte interaction. Therefore, all subsequent conclusions about astrocytes’ role in neuronal synchronization depend solely on the properties of calcium dynamics. Thus, any astrocyte model or model parameterization that reproduces the same calcium dynamics would support the same conclusions about the astrocytes’ role in neuronal synchrony.

The obtained optimal astrocyte model does not exhibit oscillatory dynamics, as this regime tends to produce too frequent and unrealistically regular calcium transients.

Instead, in the best models, calcium transients are evoked by a low-frequency Poisson noise and converge to the steady state without transients in the absence of noise inputs. Reproducing frequency, duration, and reasonable peak amplitudes of spontaneous calcium transients requires a low frequency of the input noise (about 4 spikes/s), relatively small time constant for IP_3_ (*τ*_IP3_ is about 1 s) and somewhat large IP_3_ increment at each synaptic event (Δ_IP3_ is about 0.05 µM). Simulations with active synaptic transmission require a reduction of the frequency of the input noise to about 70% of the fitted value to maintain similar levels of astrocyte inputs in the presence of synaptic glutamate processing. These fitted values likely reflect a rather simplified model of the IP_3_ receptor that increases instantaneously at each synaptic event and decreases exponentially between events. A model with richer dynamics for both astrocytic IP_3_ and calcium might be able to reproduce realistic spontaneous dynamics for a wider range of physiologically realistic model parameters.

We select the well-known depolarizing slow inward currents (SICs) as the primary focus for implementing neuron-astrocyte interactions in large-scale neuronal network simulations. SIC currents occur in many different brain regions, including the hippocampus [16, 18–21, 23, 30, 93], the thalamus [17], various areas of the cerebral cortex [28, 29], the olfactory bulb [23], the nucleus accumbens [24], and the spinal cord [25, 26]. SIC currents are observed in both physiological and pathological conditions, using various preparations such as cell cultures and brain slices *in vitro*. The activation of SIC currents occurs primarily through the calcium-dependent release of glutamate from astrocytes, the mechanism, which we select to model and further study. The calcium-dependent release of glutamate from astrocytes has long been controversial (see e.g., [94, 95]). In 2023, however, it was convincingly shown that specialized astrocytes indeed mediate gliotransmission in the central nervous system and have molecular machinery to release glutamate in a calcium-dependent manner [12].

Alternative mechanisms such as glutamate release via ion channels may also exist to activate SIC currents [96]. The molecular mechanisms underlying SIC current activation and their functional roles *in vivo*remain an important area for future research. Possible activation pathways may include diverse astrocytic calcium signaling processes or interactions with specific receptor subtypes, all of which require further investigation. Interestingly, SIC currents have been observed in both mice and humans, but their significance appears to diminish with aging in humans (no or relatively little SIC currents were found in over 70-year-old humans) [56]. Overall, the functional roles of SIC currents in the brain are not fully understood, highlighting the importance of developing computational brain models to better comprehend the role and function of SIC currents in the brain.

The computational model developed and analyzed in this study, a network model of spontaneous neuronal and astrocyte activity, focuses on a specific SIC-related function reported in the experimental literature [48, 53]. The use case model supports the emergence of synchronized spiking patterns in groups of neurons that interact with the same astrocyte. Our study confirms the result using two metrics, the first based on the timing of successive SIC-induced bursts and the second based on standard pairwise correlation, and reproduces it in networks with two different interaction schemes, and under two global spiking activity regimes. The first confirmation comes from models with sparse, asynchronous network activity regimes, where each neuron receives inputs from only one astrocyte. However, the result also holds in models where neurons receive inputs from several astrocytes, and in both asynchronous regimes (see Fig 12) and under network bursting (see Fig 6 in S3 Appendix). Switching from the asynchronous regime to network bursting is achieved by increasing the efficiency of interaction from neurons to astrocytes, which produces network bursts at the same frequency as astrocytic calcium transients, around 0.1-1.5 bursts/min. Global network bursting increases synchronization between all neuronal pairs in the model (e.g. see Fig 10D and Fig 11D) which can mask the local SIC-induced synchrony in groups of neurons. However, under the given frequency of network bursting, the effect of SIC-induced synchrony is still evident. Further increase of the network bursting frequency increases global neuronal synchronization and ultimately masks the effect of SIC current. Additionally, in this study, we consider a relatively small model, a network of 500 neurons and 100 astrocytes. Studying the role of this and similar mechanisms in realistic-size networks requires an efficient computational platform to tune the model to the desired global activity regime and simulate it for a long enough time to observe relatively slow (when compared to neuronal activity) changes in astrocyte dynamics. Our technology enables the future exploration of this and other similar mechanisms in realistic-size models.

**Fig 12.**
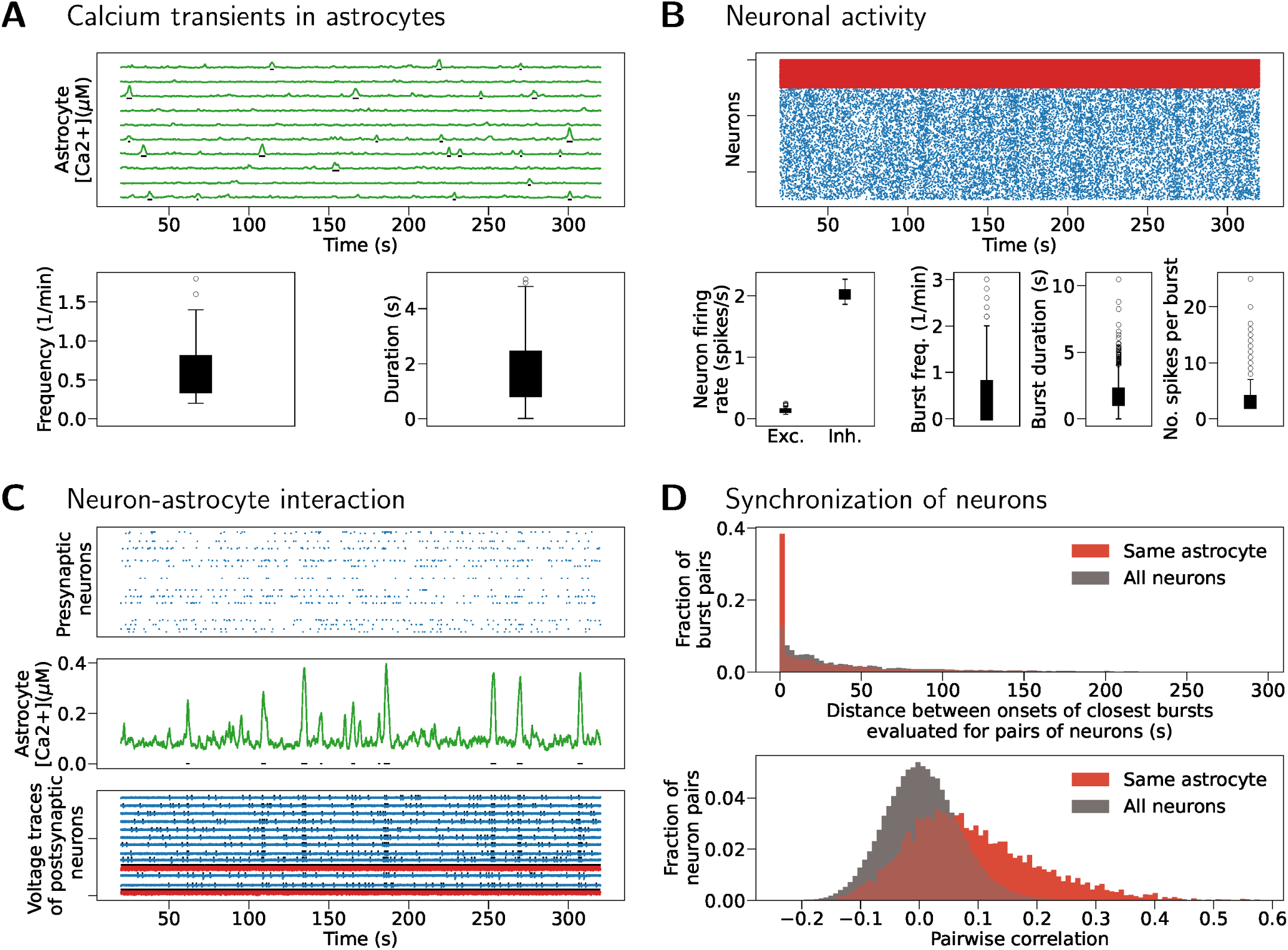
Synchronization in neuron-astrocyte networks with random pools and global low-frequency asynchronous activity. (A) Top: Astrocytic calcium transients in ten randomly selected astrocytes. Bottom: Frequency and duration of astrocytic calcium transients for all astrocytes in the model. (B) Top: Rasterplot showing activity of all excitatory (blue) and inhibitory (red) neurons in the model. Bottom left: Spiking frequency for excitatory (Exc.) and inhibitory (Inh.) neurons in the model. Bottom right: Single neuron burst frequency, duration and number of spikes per burst. (C) Single astrocyte (middle panel) with its presynaptic neuronal inputs (rasterplot, top panel) and its postsynaptic neurons (voltage traces, bottom panel). This connectivity type allows more output neurons per astrocyte and bigger variability in number and type (excitatory vs inhibitory) of neurons per astrocyte compared to the block-type connectivity. (D) Top: Neuronal synchrony is evaluated as a difference in onsets of successive burst events. Bottom: Synchrony evaluated as pairwise correlation between binned spike trains.

The functional role of SIC-mediated synchronicity and astrocytic calcium signaling remains unclear, but it might be relevant in all functions that require coherent activation of neuronal groups, such as plasticity or memory retrieval. We here focus on one specific mechanism, but synchronization in brain circuits might arise also through other astrocyte-dependent mechanisms and through the release of both glutamate and GABA, as shown *in vivo* in a sequence of studies [49, 50, 52]. Astrocytic release of neurotransmitters dynamically changes the extracellular composition of molecular species which modulates brain circuits and can induce changes in activity regimes [49], contribute to pathological states [50, 52] and pathological behavior as shown in [50]. The diversity of mechanisms through which astrocytes and their calcium signaling contribute to the activity of brain circuits and understanding their interplay in *in vitro* and *in vivo* further highlights the need for efficient simulation tools that can integrate diverse mechanisms into models of brain circuits.

While astrocyte-dependent synchronization is central to our simulation use case, astrocyte physiology is a rapidly evolving field, uncovering the complex and context-dependent roles of astrocytes and their calcium signaling in neural function and behavior [97–100]. Intracellular calcium signaling in astrocytes can lead to diverse physiological outcomes, such as brain state changes [49], sensory-evoked neuronal network activity [51], and modulation of synaptic transmission and plasticity [101].

Astrocytes can be activated by various substances through membrane mechanisms, triggering the calcium-dependent release of gliotransmitters such as glutamate, ATP, and D-serine [12, 13]. This release can enhance or suppress synaptic activity, transmission, and plasticity, depending on the context, brain area, biological sex, and receptor involvement [101–104]. Some effects likely occur through distinct intracellular signaling pathways following calcium activation, though a major challenge in experimental research has been the lack of astrocyte-specific pharmacological tools.

Nevertheless, it is now clear that astrocyte signal integration involves complex, yet identifiable intracellular pathways and receptor systems that regulate their role in neural circuit function [99, 105]. The novel theoretical methodologies for modeling tripartite connectivity and the scalable computational simulation tools, both developed in our study, will help address this challenging goal.

### Reproducibility and future considerations

The traceability of model implementations is crucial, necessitating open accessibility, such as including codes in model databases and clarifying the evolutionary path from previously published models. To maintain the principles of FAIR (Findable, Accessible, Interoperable, and Reusable) data, the use of standardized simulation tools and data-analysis and sensitivity-analysis methodologies are necessary for all computational fields [106]. Advancing model development workflows and expanding neuroinformatics tools for astrocytes are crucial in enhancing reproducibility, standardization, and sharing of astrocyte models. To facilitate comprehensive specification, we recommend employing a predefined format for describing model components and interaction schemes. In this regard, formats proposed for specifying neuronal networks [37] and connectivity schemes between neurons [4] can be expanded to accommodate neuron-astrocyte networks. All aspects of models, including network structure, cell count, interaction dynamics, and all the equations, initial and parameter values, must be explicitly disclosed as suggested recently [35]. The description of interaction schemes should seamlessly intertwine with model equations, ensuring that the reconstruction of these schemes is achievable solely through the equations [35].

Properly documenting model parameters is equally important. Expanding the level of biological details in a model, adding new components such as astrocyte-related mechanisms, or increasing the model size inevitably leads to a larger and more complex parameter space. Exhaustive evaluation of the entire parameter space becomes challenging already for relatively small models. On the other hand, fitting the parameters to the experimental data often suffers from the lack of data as well as from the presence of strong nonlinearities in models which result in complex, non-convex, multi-objective optimization tasks. Consequently, new studies often reuse the existing models and previously reported parameters, and they depend on the accurate and systematic description of models and parameters.

All these advancements are crucial for expediting model development, adding greater biological knowledge into data-driven models, and integrating astrocytic mechanisms into large-scale, realistic models of brain systems. Our reference implementation for neuron-astrocyte networks helps to catalyze the development of neuron-astrocyte network models by providing fast simulation times and a reliable model implementation via tripartite connectivity generation, not present in other similar simulation tools.

Computational models and tools in brain research have become crucial in advancing our understanding of brain functions, dysfunctions, and diseases [107]. These allow researchers to simulate and analyze complex neural processes, providing valuable insights into the functioning of the brain and helping to solve the mysteries of neurological phenomena. They enable the exploration of various scenarios and aid in deciphering the underlying mechanisms of both normal brain function and aberrations leading to disorders. In addition, they facilitate the development of hypotheses and the testing of theories, offering possibilities for breakthroughs in neuroscience research.

## Supporting information

**S1 Appendix.** Description of the neuron-astrocyte network models

**S2 Appendix.** Variables and parameters added to the NEST support for astrocytes

**S3 Appendix.** Details of construction of use case model including parameter selection. Additional data supporting the findings about astrocytes’ role in neuronal synchrony.

**S4 Appendix.** Implementation of tripartite connectivity in the NEST Simulator

## Author Contributions

**Conceptualization:** H-JJ, JA, TM, MD, SJvA, HEP, M-LL

**Data curation:** H-JJ, JA

**Formal analysis:** H-JJ, JA

**Funding acquisition:** JA, TM, MD, SJvA, HEP, M-LL

**Investigation:** H-JJ, JA, TM

**Methodology:** H-JJ, JA, TM, JS, ML, MD, SJvA, HEP, M-LL

**Project administration:** H-JJ, JA

**Resources:** MD, SJvA, HEP

**Software:** H-JJ, JA, TM, IA, JS, ML, HEP

**Supervision:** JA, MD, SJvA, HEP, M-LL

**Validation:** H-JJ, JA, TM, IA, JS, MD, SJvA, HEP, M-LL

**Visualization:** H-JJ, JA, TM, M-LL

**Writing – original draft Preparation:** H-JJ, JA, TM, JS, MD, HEP, M-LL

**Writing – review & editing:** H-JJ, JA, TM, IA, MD, SJvA, HEP, M-LL

## Acknowledgments

We thank Dr. Tom Tetzlaff and Melissa Lober for the communication that allowed us to implement a surrogate model for the astrocyte, which is adapted from their implementation of the ignore_and_fire neuron model in NEST (personal communication). We thank Dr. David Dahmen for the consultation on the continuous interaction framework for sic_connection. We thank Dr. Noora Pihlajarinne for her work on panels in Fig 1 for the illustration of neuron-astrocyte interactions and Fig 1 in S2 Appendix for the illustration of software components required for the representation of the model and their embedding in the NEST code. We further thank Agnes Korcsak-Gorzo and Jose Villamar for help on the installation and workflow implementation with beNNch as well as Pooja Babu for her help in disseminating preliminary results of this work. Dr. Jugoslava Ácimovíc is currently employed as senior data scientist at AstraZeneca, however the research presented here was conducted during her previous employment at Tampere University and is unrelated to her current employment. Thus, her affiliation with Tampere University is used in this article.

## References

1. Einevoll GT, Destexhe A, Diesmann M, Grün S, Jirsa V, de Kamps M, et al. The scientific case for brain simulations. Neuron. 2019;102(4):735–744. doi:10.1016/j.neuron.2019.03.027.

2. Hahne J, Helias M, Kunkel S, Igarashi J, Bolten M, Frommer A, et al. A unified framework for spiking and gap-junction interactions in distributed neuronal network simulations. Frontiers in Neuroinformatics. 2015;9:22. doi:10.3389/fninf.2015.00022.

3. Sherman SM, Guillery RW. The Role of the Thalamus in the Flow of Information to the Cortex. Philosophical transactions of the Royal Society of London B: Biological Sciences. 2002;357(1428):1695–1708. doi:10.1098/rstb.2002.1161.

4. Senk J, Kriener B, Djurfeldt M, Voges N, Jiang HJ, Schüttler L, et al. Connectivity concepts in neuronal network modeling. PLoS Computational Biology. 2022;18(9):e1010086. doi:10.1371/journal.pcbi.1010086.

5. Bushong EA, Martone ME, Jones YZ, Ellisman MH. Protoplasmic astrocytes in CA1 stratum radiatum occupy separate anatomical domains. Journal of Neuroscience. 2002;22(1):183–192. doi:10.1523/JNEUROSCI.22-01-00183.2002.

6. Oberheim NA, Takano T, Han X, He W, Lin JHC, Wang F, et al. Uniquely hominid features of adult human astrocytes. Journal of Neuroscience. 2009;29(10):3276–3287. doi:10.1523/JNEUROSCI.4707-08.2009.

7. Vasile F, Dossi E, Rouach N. Human astrocytes: structure and functions in the healty brain. Brain Structure and Function. 2017;222:2017–2029. doi:10.1007/s00429-017-1383-5.

8. Calì C, Agus M, Kare K, Boges DJ, Lehväslaiho H, Hadwiger M, et al. 3D cellular reconstruction of cortical glia and parenchymal morphometric analysis from Serial Block-Face Electron Microscopy of juvenile rat. Progress in Neurobiology. 2019;183:101696. doi:10.1016/j.pneurobio.2019.101696.

9. Kikuchi T, Gonzalez-Soriano J, Kastanauskaite A, Benavides-Piccione R, Merchan-Perez A, DeFelipe J, et al. Volume electron microscopy study of the relationship between synapses and astrocytes in the developing rat somatosensory cortex. Cerebral Cortex. 2020;30(6):3800–3819. doi:10.1093/cercor/bhz343.

10. Zisis E, Keller D, Kanari L, Arnaudon A, Gevaert M, Delemontex T, et al. Digital reconstruction of the neuro-glia-vascular architecture. Cerebral Cortex. 2021;31(12):5686–5703. doi:10.1093/cercor/bhab254.

11. Aten S, Kiyoshi CM, Arzola EP, Patterson JA, Taylor AT, Du Y, et al. Ultrastructural view of astrocyte arborization, astrocyte-astrocyte and astrocyte-synapse contacts, intracellular vesicle-like structures, and mitochondrial network. Progress in Neurobiology. 2022;213:102264. doi:10.1016/j.pneurobio.2022.102264.

12. de Ceglia R, Ledonne A, Litvin DG, Lind BL, Carriero G, Latagliata EC, et al. Specialized astrocytes mediate glutamatergic gliotransmission in the CNS. Nature. 2023;622(7981):120–129. doi:10.1038/s41586-023-06502-w.

13. Araque A, Parpura V, Sanzgiri RP, Haydon PG. Tripartite synapses: glia, the unacknowledged partner. Trends in Neurosciences. 1999;22(5):208–215. doi:10.1016/S0166-2236(98)01349-6.

14. Parpura V, Basarsky TA, Liu F, Jeftinija K, Jeftinija S, Haydon PG. Glutamate-mediated astrocyte–neuron signalling. Nature. 1994;369:744–747. doi:10.1038/369744a0.

15. Navarrete M, Perea G, de Sevilla DF, Gómez-Gonzalo M, Núñez A, Martín ED, et al. Astrocytes mediate in vivo cholinergic-induced synaptic plasticity. PLoS Biology. 2012;10(2):e1001259. doi:10.1371/journal.pbio.1001259.

16. Araque A, Parpura V, Sanzgiri RP, Haydon PG. Glutamate-dependent astrocyte modulation of synaptic transmission between cultured hippocampal neurons. European Journal of Neuroscience. 1998;10:2129–2142. doi:10.1046/j.1460-9568.1998.00221.x.

17. Parri HR, Gould TM, Crunelli V. Spontaneous astrocytic Ca^2+^ oscillations in situ drive NMDAR-mediated neuronal excitation. Nature Neuroscience. 2001;4(8):803–812. doi:10.1038/90507.

18. Angulo MC, Kozlov AS, Charpak S, Audinat E. Glutamate released from glial cells synchronizes neuronal activity in the hippocampus. Journal of Neuroscience. 2004;24(31):6920–6927. doi:10.1523/JNEUROSCI.0473-04.2004.

19. Fellin T, Pascual O, Gobbo S, Pozzan T, Haydon PG, Carmignoto G. Neuronal synchrony mediated by astrocytic glutamate through activation of extrasynaptic NMDA receptors. Neuron. 2004;43(5):729–743. doi:10.1016/j.neuron.2004.08.011.

20. Perea G, Araque A. Properties of synaptically evoked astrocyte calcium signal reveal synaptic information processing by astrocytes. Journal of Neuroscience. 2005;25(9):2192–2203. doi:10.1523/JNEUROSCI.3965-04.2005.

21. Carmignoto G, Fellin T. Glutamate release from astrocytes as a non-synaptic mechanism for neuronal synchronization in the hippocampus. Journal of Physiology (Paris). 2006;99(2-3):98–102. doi:10.1016/j.jphysparis.2005.12.008.

22. Gao WJ, Goldman-Rakic PS. NMDA receptor-mediated epileptiform persistent activity requires calcium release from intracellular stores in prefrontal neurons. Experimental Neurology. 2006;197(2):495–504. doi:10.1016/j.expneurol.2005.05.018.

23. Kozlov AS, Angulo MC, Audinat E, Charpak S. Target cell-specific modulation of neuronal activity by astrocytes. Proceedings of the National Academy of Sciences of the United States of America. 2006;103(26):10058–10063. doi:10.1073/pnas.0603741103.

24. D’Ascenzo M, Fellin T, Terunuma M, Revilla-Sanchez R, Meaney DF, Auberson YP, et al. mGluR5 stimulates gliotransmission in the nucleus accumbens. Proceedings of the National Academy of Sciences of the United States of America. 2007;104(6):1995–2000. doi:10.1073pnas.0609408104.

25. Bardoni R, Ghirri A, Zonta M, Betelli C, Vitale G, Ruggieri V, et al. Glutamate-mediated astrocyte-to-neuron signalling in the rat dorsal horn. Journal of Physiology. 2010;588(5):831–846. doi:10.1113/jphysiol.2009.180570.

26. Nie H, Zhang H, Weng HR. Bidirectional neuron–glia interactions triggered by deficiency of glutamate uptake at spinal sensory synapses. Journal of Neurophysiology. 2010;104(2):713–725. doi:10.1152/jn.00282.2010.

27. Reyes-Haro D, Müller J, Boresch M, Pivneva T, Benedetti B, Scheller A, et al. Neuron–astrocyte interactions in the medial nucleus of the trapezoid body. Journal of General Physiology. 2010;135(6):583–594. doi:10.1085/jgp.200910354.

28. Chen N, Sugihara H, Sharma J, Perea G, Petravicz J, Le C, et al. Nucleus basalis-enabled stimulus-specific plasticity in the visual cortex is mediated by astrocytes. Proceedings of the National Academy of Sciences of the United States of America. 2012;109(41):E2832–E2841. doi:10.1073/pnas.1206557109.

29. Perea G, Yang A, Boyden ES, Sur M. Optogenetic astrocyte activation modulates response selectivity of visual cortex neurons *in vivo*. Nature Communications. 2014;5:3262. doi:10.1038/ncomms4262.

30. Lauderdale K, Murphy T, Tung T, Davila D, Binder DK, Fiacco TA. Osmotic edema rapidly increases neuronal excitability through activation of NMDA receptor-dependent slow inward currents in juvenile and adult hippocampus. ASN Neuro. 2015;7(5):1–12. doi:10.1177/1759091415605115.

31. Navarrete M, Perea G, Maglio L, Pastor J, de Sola RG, Araque A. Astrocyte calcium signal and gliotransmission in human brain tissue. Cerebral Cortex. 2013;23:1240–1246. doi:10.1093/cercor/bhs122.

32. Manninen T, Ácimovíc J, Havela R, Teppola H, Linne ML. Challenges in reproducibility, replicability, and comparability of computational models and tools for neuronal and glial networks, cells, and subcellular structures. Frontiers in Neuroinformatics. 2018;12:20. doi:10.3389/fninf.2018.00020.

33. Manninen T, Havela R, Linne ML. Computational models for calcium-mediated astrocyte functions. Frontiers in Computational Neuroscience. 2018;12:14. doi:10.3389/fncom.2018.00014.

34. Manninen T, Havela R, Linne ML. Computational models of astrocytes and astrocyte-neuron interactions: characterization, reproducibility, and future perspectives. In: De Pittà M, Berry H, editors. Computational Glioscience. Cham, Switzerland: Springer; 2019. p. 423–454.

35. Manninen T, Ácimovíc J, Linne ML. Analysis of network models with neuron-astrocyte interactions. Neuroinformatics. 2023;21:375–406. doi:10.1007/s12021-023-09622-w.

36. Linne ML, Ácimovíc J, Saudargiene A, Manninen T. Neuron–glia interactions and brain circuits. In: Giugliano M, Negrello M, Linaro D, editors. Computational Modelling of the Brain: Modelling Approaches to Cells, Circuits and Networks. Cham, Switzerland: Springer; 2022. p. 87–103.

37. Nordlie E, Gewaltig MO, Plesser HE. Towards reproducible descriptions of neuronal network models. PLoS Computational Biology. 2009;5(8):e1000456. doi:10.1371/journal.pcbi.1000456.

38. Graber S, Mitchell J, Kurth AC, Terhorst D, Skaar JEW, Schöfmann CM, et al.. NEST 3.8; 2024. Available from: 10.5281/zenodo.12624784.

39. Gewaltig MO, Diesmann M. NEST (neural simulation tool). Scholarpedia. 2007;2(4):1430.

40. Aleksin SG, Zheng K, Rusakov DA, Savtchenko LP. ARACHNE: a neural-neuroglial network builder with remotely controlled parallel computing. PLoS Computational Biology. 2017;13(3):e1005467. doi:10.1371/journal.pcbi.1005467.

41. Stimberg M, Brette R, Goodman DFM. Brian 2, an intuitive and efficient neural simulator. eLife. 2019;8:e47314. doi:10.7554/eLife.47314.

42. Stimberg M, Goodman DFM, Brette R, De Pittà M. Modeling neuron–glia interactions with the Brian 2 simulator. In: De Pittà M, Berry H, editors. Computational Glioscience. Cham, Switzerland: Springer; 2019. p. 471–505.

43. De Young GW, Keizer J. A single-pool inositol 1,4,5-trisphosphate-receptor-based model for agonist-stimulated oscillations in Ca2+ concentration. Proceedings of the National Academy of Sciences of the United States of America. 1992;89(20):9895–9899. doi:10.1073/pnas.89.20.9895.

44. Li YX, Rinzel J. Equations for InsP_3_ receptor-mediated [Ca^2+^]_i_ oscillations derived from a detailed kinetic model: a Hodgkin-Huxley like formalism. Journal of Theoretical Biology. 1994;166(4):461–473. doi:10.1006/jtbi.1994.1041.

45. Nadkarni S, Jung P. Spontaneous oscillations of dressed neurons: a new mechanism for epilepsy? Physical Review Letters. 2003;91(26):268101. doi:10.1103/PhysRevLett.91.268101.

46. Allegrini P, Fronzoni L, Pirino D. The influence of the astrocyte field on neuronal dynamics and synchronization. Journal of Biological Physics. 2009;35(4):413–423. doi:10.1007/s10867-009-9166-8.

47. Liu Y, Li C. Firing rate propagation through neuronal–astrocytic network. IEEE Transactions on Neural Networks and Learning Systems. 2013;24(5):789–799. doi:10.1109/TNNLS.2013.2245678.

48. Pirttimaki TM, Sims RE, Saunders G, Antonio SA, Codadu NK, Parri HR. Astrocyte-mediated neuronal synchronization properties revealed by false gliotransmitter release. Journal of Neuroscience. 2017;37(41):9859–9870. doi:10.1523/JNEUROSCI.2761-16.2017.

49. Poskanzer KE, Yuste R. Astrocytes regulate cortical state switching in vivo. Proceedings of the National Academy of Sciences of the United States of America. 2016;113(19):E2675–E2684. doi:10.1073/pnas.1520759113.

50. Yu X, Taylor AMW, Nagai J, Golshani P, Evans CJ, Coppola G, et al. Reducing astrocyte calcium signaling in vivo alters striatal microcircuits and causes repetitive behavior. Neuron. 2018;99(6):1170–1187. doi:10.1016/j.neuron.2018.08.015.

51. Lines J, Martin ED, Kofuji P, Aguilar J, Araque A. Astrocytes modulate sensory-evoked neuronal network activity. Nature Communications. 2020;11:3689. doi:10.1038/s41467-020-17536-3.

52. Magloire V, Savtchenko LP, Jensen TP, Sylantyev S, Kopach O, Cole N, et al. Volume-transmitted GABA waves pace epileptiform rhythms in the hippocampal network. Current Biology. 2023;33(7):1249–1264. doi:10.1016/j.cub.2023.02.051.

53. Pirttimaki TM, Hall SD, Parri HR. Sustained neuronal activity generated by glial plasticity. Journal of Neuroscience. 2011;31(21):7637–7647. doi:10.1523/JNEUROSCI.5783-10.2011.

54. Wetherington J, Serrano G, Dingledine R. Astrocytes in the epileptic brain. Neuron. 2008;58(2):168–178. doi:10.1016/j.neuron.2008.04.002.

55. Dong QP, He JQ, Chai Z. Astrocytic Ca^2+^ waves mediate activation of extrasynaptic NMDA receptors in hippocampal neurons to aggravate brain damage during ischemia. Neurobiology of Disease. 2013;58:68–75. doi:10.1016/j.nbd.2013.05.005.

56. Csemer A, Kovács A, Maamrah B, Pocsai K, Korpás K, Klekner Á, et al. Astrocyte- and NMDA receptor-dependent slow inward currents differently contribute to synaptic plasticity in an age-dependent manner in mouse and human neocortex. Aging Cell. 2023;22(9):e13939. doi:10.1111/acel.13939.

57. Kovács A, Pál B. Astrocyte-dependent slow inward currents (SICs) participate in neuromodulatory mechanisms in the pedunculopontine nucleus (PPN). Frontiers in Cellular Neuroscience. 2017;11:16. doi:10.3389/fncel.2017.00016.

58. Pirttimaki TM, Parri HR. Glutamatergic input–output properties of thalamic astrocytes. Neuroscience. 2012;205:18–28. doi:10.1016/j.neuroscience.2011.12.049.

59. Srinivasan R, Huang BS, Venugopal S, Johnston AD, Chai H, Zeng H, et al. Ca^2+^ signaling in astrocytes from Ip3r2^-/-^ mice in brain slices and during startle responses *in vivo*. Nature Neuroscience. 2015;18(5):708–717. doi:10.1038/nn.4001.

60. Stobart JL, Ferrari KD, Barrett MJP, Glück C, Stobart MJ, Zuend M, et al. Cortical circuit activity evokes rapid astrocyte calcium signals on a similar timescale to neurons. Neuron. 2018;98(4):726–735. doi:10.1016/j.neuron.2018.03.050.

61. Arizono M, Inavalli VVG, Panatier A, Pfeiffer T, Angibaud J, Levet F, et al. Structural basis of astrocytic Ca^2+^ signals at tripartite synapses. Nature Communications. 2020;11(1):1906. doi:10.1038/s41467-020-15648-4.

62. Ácimovíc J, Jiang HJ, Manninen T, Stapmanns J, Lehtimäki M, Linne ML, et al. The NEST module for computational modeling of neuron-astrocyte networks. In: Proceedings of Annual Computational Neuroscience Meeting. Leipzig, Germany; 2023.

63. Jiang HJ, Ácimovíc J, Manninen T, Stapmanns J, Lehtimäki M, Linne ML, et al. Computational modelling of neuron-astrocyte interactions in the NEST simulator. In: Proceedings of Göttingen Meeting of the German Neuroscience Society. Göttingen, Germany; 2023.

64. Villamar J, Vogelsang J, Linssen C, Kunkel S, Kurth A, Schöfmann CM, et al.. NEST 3.6; 2023. Available from: 10.5281/zenodo.8344932.

65. Espinoza Valverde JA, Müller E, Haug N, Schöfmann CM, Linssen C, Senk J, et al.. NEST 3.7; 2024. Available from: 10.5281/zenodo.10834751.

66. Plesser HE. Astrocyte Surrogate Module; 2024. Available from: https://github.com/heplesser/astrocyte-surrogate-module.

67. Brette R, Gerstner W. Adaptive exponential integrate-and-fire model as an effective description of neuronal activity. Journal of Neurophysiology. 2005;94:3637–3642. doi:10.1152/jn.00686.2005.

68. Naud R, Marcille M, Clopath C, Gerstner W. Firing patterns in the adaptive exponential integrate-and-fire model. Biological Cybernetics. 2008;99:335–347. doi:10.1007/s00422-008-0264-7.

69. Milo R, Shen-Orr S, Itzkovitz S, Kashtan N, Chklovskii D, Alon U. Network motifs: simple building blocks of complex networks. Science. 2002;298(5594):824–827. 10.1126/science.298.5594.824.

70. Pernice V, Staude B, Cardanobile S, Rotter S. How structure determines correlations in neuronal networks. PLoS Computational Biology. 2011;7(5):e1002059. 10.1371/journal.pcbi.1002059.

71. Ácimovíc J, Mäki-Marttunen T, Linne ML. The effects of neuron morphology on graph theoretic measures of network connectivity: The analysis of a two-level statistical model. Frontiers in Neuroanatomy. 2015;9. doi:10.3389/fnana.2015.00076.

72. Jovanovíc S, Rotter S. Interplay between Graph Topology and Correlations of Third Order in Spiking Neuronal Networks. PLoS Computational Biology. 2016;12(6):1–28. doi:10.1371/journal.pcbi.1004963.

73. Halassa MM, Fellin T, Takano H, Dong JH, Haydon PG. Synaptic islands defined by the territory of a single astrocyte. Journal of Neuroscience. 2007;27(24):6473–6477. doi:10.1523/JNEUROSCI.1419-07.2007.

74. Morrison A, Mehring C, Geisel T, Aertsen A, Diesmann M. Advancing the boundaries of high connectivity network simulation with distributed computing. Neural Computation. 2005;17:1776–1801. doi:10.1162/0899766054026648.

75. Jordan J, Ippen T, Helias M, Kitayama I, Sato M, Igarashi J, et al. Extremely scalable spiking neuronal network simulation code: From laptops to exascale computers. Frontiers in Neuroinformatics. 2018;12:2. doi:10.3389/fninf.2018.00002.

76. Plesser HE, Eppler JM, Morrison A, Diesmann M, Gewaltig MO. Efficient Parallel Simulation of Large-Scale Neuronal Networks on Clusters of Multiprocessor Computers. In: Kermarrec AM, Bouǵe L, Priol T, editors. Euro-Par 2007: Parallel Processing. vol. 4641 of Lecture Notes in Computer Science. Berlin: Springer-Verlag; 2007. p. 672–681.

77. Albers J, Pronold J, Kurth AC, Vennemo SB, Mood KH, Patronis A, et al. A modular workflow for performance benchmarking of neuronal network simulations. Frontiers in Neuroinformatics. 2022;16:837549. doi:10.3389/fninf.2022.837549.

78. Brunel N. Dynamics of sparsely connected networks of excitatory and inhibitory spiking neurons. Journal of Computational Neuroscience. 2000;8:183–208. doi:10.1023/A:1008925309027.

79. Van Albada SJ, Rowley AG, Senk J, Hopkins M, Schmidt M, Stokes AB, et al. Performance Comparison of the Digital Neuromorphic Hardware SpiNNaker and the Neural Network Simulation Software NEST for a Full-Scale Cortical Microcircuit Model. Frontiers in Neuroscience. 2018;12:291. doi:10.3389/fnins.2018.00291.

80. Nedergaard M, Ransom B, Goldman SA. New roles for astrocytes: redefining the functional architecture of the brain. Trends in Neurosciences. 2003;26(10):523–530. doi:10.1016/j.tins.2003.08.008.

81. Van Albada SJ, Helias M, Diesmann M. Scalability of asynchronous networks is limited by one-to-one mapping between effective connectivity and correlations. PLoS Computational Biology. 2015;11(9):e1004490. doi:10.1371/journal.pcbi.1004490.

82. Neske GT, Patrick SL, Connors BW. Contributions of diverse excitatory and inhibitory neurons to recurrent network activity in cerebral cortex. Journal of Neuroscience. 2015;35(3):1089–1105. doi:10.1523/JNEUROSCI.2279-14.2015.

83. Sun QQ, Huguenard JR, Prince DA. Barrel cortex microcircuits: thalamocortical feedforward inhibition in spiny stellate cells is mediated by a small number of fast-spiking interneurons. Journal of Neuroscience. 2006;26(4):1219–1230. doi:10.1523/JNEUROSCI.4727-04.2006.

84. Touboul J, Brette R. Dynamics and bifurcations of the adaptive exponential integrate-and-fire model. Biological Cybernetics. 2008;99:319–334. doi:10.1007/s00422-008-0267-4.

85. Thörnig P. JURECA: Data centric and booster modules implementing the modular supercomputing architecture at Jülich supercomputing centre. Journal of large-scale research facilities JLSRF. 2021;7:A182–A182.

86. De Pittà M, Brunel N. Multiple forms of working memory emerge from synapse–astrocyte interactions in a neuron–glia network model. Proceedings of the National Academy of Sciences of the United States of America. 2022;119(43):e2207912119. doi:10.1073/pnas.2207912119.

87. Moyse LB, Berry H. Modelling the modulation of cortical Up-Down state switching by astrocytes. PLoS Computational Biology. 2022;18(7):e1010296. doi:10.1371/journal.pcbi.1010296.

88. Halassa MM, Fellin T, Haydon PG. The tripartite synapse: roles for gliotransmission in health and disease. Trends in Molecular Medicine. 2007;13(2):54–63. doi:10.1016/j.molmed.2006.12.005.

89. Savtchenko LP, Bard L, Jensen TP, Reynolds JP, Kraev I, Medvedev N, et al. Disentangling astroglial physiology with a realistic cell model in silico. Nature Communications. 2018;9(1):3554. doi:10.1038/s41467-018-05896-w.

90. Sasaki T, Kuga N, Namiki S, Matsuki N, Ikegaya Y. Locally synchronized astrocytes. Cerebral Cortex. 2011;21(8):1889–1900. doi:10.1093/cercor/bhq256.

91. Nakayama R, Sasaki T, Tanaka KF, Ikegaya Y. Subcellular calcium dynamics during juvenile development in mouse hippocampal astrocytes. European Journal of Neuroscience. 2016;43(7):923–932. doi:10.1111/ejn.13188.

92. Stobart JL, Ferrari KD, Barrett MJP, Stobart MJ, Looser ZJ, Saab AS, et al. Long-term in vivo calcium imaging of astrocytes reveals distinct cellular compartment responses to sensory stimulation. Cerebral Cortex. 2018;28(1):184–198. doi:10.1093/cercor/bhw366.

93. Araque A, Sanzgiri RP, Parpura V, Haydon PG. Calcium elevation in astrocytes causes an NMDA receptor-dependent increase in the frequency of miniature synaptic currents in cultured hippocampal neurons. Journal of Neuroscience. 1998;18(17):6822–6829. doi:10.1523/JNEUROSCI.18-17-06822.1998.

94. Fiacco TA, McCarthy KD. Multiple lines of evidence indicate that gliotransmission does not occur under physiological conditions. Journal of Neuroscience. 2018;38(1):3–13. doi:10.1523/JNEUROSCI.0016-17.2017.

95. Savtchouk I, Volterra A. Gliotransmission: beyond black-and-white. Journal of Neuroscience. 2018;38(1):14–25. doi:10.1523/JNEUROSCI.0017-17.2017.

96. Gómez-Gonzalo M, Zehnder T, Requie LM, Bezzi P, Carmignoto G. Insights into the release mechanism of astrocytic glutamate evoking in neurons NMDA receptor-mediated slow depolarizing inward currents. Glia. 2018;66(10):2188–2199. doi:10.1002/glia.23473.

97. Khakh BS, Sofroniew MV. Diversity of astrocyte functions and phenotypes in neural circuits. Nature Neuroscience. 2015;18(7):942–952. doi:10.1038/nn.4043.

98. Santello M, Toni N, Volterra A. Astrocyte function from information processing to cognition and cognitive impairment. Nature Neuroscience. 2019;22(2):154–166. doi:10.1038/s41593-018-0325-8.

99. Yu X, Nagai J, Marti-Solano M, Soto JS, Coppola G, Babu MM, et al. Context-specific striatal astrocyte molecular responses are phenotypically exploitable. Neuron. 2020;108(6):1146–1162. doi:10.1016/j.neuron.2020.09.021.

100. Hirrlinger J, Nimmerjahn A. A perspective on astrocyte regulation of neural circuit function and animal behavior. Glia. 2022;70(8):1554–1580. doi:10.1002/glia.24168.

101. Perea G, Araque A. Astrocytes potentiate transmitter release at single hippocampal synapses. Science. 2007;317(5841):1083–1086. doi:10.1126/science.1144640.

102. Min R, Nevian T. Astrocyte signaling controls spike timing–dependent depression at neocortical synapses. Nature Neuroscience. 2012;15(5):746–753. doi:10.1038/nn.3075.

103. Falćon-Moya R, Pérez-Rodŕıguez M, Prius-Mengual J, Andrade-Talavera Y, Arroyo-Garćıa LE, Pérez-Artés R, et al. Astrocyte-mediated switch in spike timing-dependent plasticity during hippocampal development. Nature Communications. 2020;11(1):4388. doi:10.1038/s41467-020-18024-4.

104. Noh K, Cho WH, Lee BH, Kim DW, Kim YS, Park K, et al. Cortical astrocytes modulate dominance behavior in male mice by regulating synaptic excitatory and inhibitory balance. Nature Neuroscience. 2023;26(9):1541–1554. doi:10.1038/s41593-023-01406-4.

105. Verkhratsky A, Nedergaard M. Physiology of astroglia. Physiological Reviews. 2018;98(1):239–389. doi:10.1152/physrev.00042.2016.

106. Eriksson O, Bhalla US, Blackwell KT, Crook SM, Keller D, Kramer A, et al. Combining hypothesis- and data-driven neuroscience modeling in FAIR workflows. eLife. 2022;11:e69013. doi:10.7554/eLife.69013.

107. Amunts K, Axer M, Banerjee S, Bitsch L, Bjaalie JG, Brauner P, et al. The coming decade of digital brain research: A vision for neuroscience at the intersection of technology and computing. Imaging Neuroscience. 2024;2:1–35. doi:10.1162/imaga00137.

