## Supplementary material for "Modeling neuron-astrocyte interactions in neural networks using distributed simulation": S1 Appendix

### S1 Appendix: Description of the neuron-astrocyte network models

This appendix contains details of the neuron-astrocyte network models considered in this article, following the convention of previous studies (Nordlie et al., 2009; Senk et al., 2022). Table A contains a general and systematic description of the models in a standardized tabular format summarizing model components, mathematical equations, and inputs and outputs. Tables B and C list specific parameter values of the models. Table B lists parameter values used in the benchmarks of this study. Table C lists parameter values used to construct the *in silico* model based on the experimental study by Pirttimaki et al. (2017). This model reproduces and extends the previously presented analysis of astrocytic role in neuronal synchronization (Pirttimaki et al., 2017). The parameters are carefully fitted to the experimental findings from the literature as described in Methods and Results of the article and S3 Appendix. Notations in Table B and C follow the style of mathematical descriptions in this article, or the style of NEST parameters if there is no corresponding notation in the mathematical descriptions (see S2 Appendix).

Table A: Neuron-astrocyte network model presented in the standardized tabular format.

| **Model summary** | |
|---|---|
| **Populations** | Excitatory neurons (E), Inhibitory neurons (I), Astrocytes (A) |
| **Topology** | Random connectivity, no spatial organization |
| **Connectivity** | |
| Neuron-to-neuron | Each pair of neurons is connected with probability $p_{\text{primary}}$. |
| Astrocyte-to-neuron, Neuron-to-astrocyte | Each generated excitatory synapse interacts with one selected astrocyte with probability $p_{\text{third\_if\_primary}}$. In case interaction is established, connections from the presynaptic neuron to astrocyte, and from the astrocyte to the postsynaptic neuron are generated. |
| **Neuron model** | Adaptive Exponential Integrate-and-fire (AdEx) (Brette and Gerstner, 2005; Naud et al., 2008) |
| **Astrocyte model** | $[Ca^{2+}]$ dynamics according to De Young-Keizer (De Young and Keizer, 1992) and Li-Rinzel (Li and Rinzel, 1994) models; Slow Inward Current (SIC) according to Nadkarni-Jung (NJ) model (Nadkarni and Jung, 2003) |
| **Synapse model** | |
| Neuron-to-neuron, neuron-to-astrocyte | Short-term facilitation and depression according to Tsodyks-Markram model (Tsodyks et al., 1998, 2000). The kinetic is exponentially decaying for neuron-to-neuron interactions, and delta function for neuron-to-astrocyte interactions. |
| Astrocyte-to-neuron | Delivers continuous SIC to the target (postsynaptic) neuron |
| **External inputs** | |
| Neuron | Poisson noise that induces postsynaptic current |
| Astrocyte | 1) Poisson noise that affects $IP_3$ concentration<br>2) Gaussian noise that affects calcium concentration |

| **Populations** | | | |
|---|---|---|---|
| Name | Elements | Size (no. neurons) | Indices in connectivity matrices $\mathbf{\Gamma}_{\text{NN}}$, $\mathbf{\Gamma}_{\text{NA}}$,$\mathbf{\Gamma}_{\text{AN}}$ |
| Excitatory neurons | AdEx | $N_{\text{E}}$ | $i = 1...N_{\text{E}}$ |
| Inhibitory neurons | AdEx | $N_{\text{I}} = N - N_{\text{E}}$ | $i = (N_{\text{E}} + 1)...N$ |
| Astrocytes | NJ | $N_{\text{A}}$ | $k = 1...N_{\text{A}}$ |

| **Connectivity** | | |
|---|---|---|
| Source | Target | Pattern |
| {E,I} | {E,I} | Connectivity matrix $\mathbf{\Gamma}_{\mathrm{NN}} = \{\gamma_{ji}^{\mathrm{NN}}\}_{i,j=1\ldots N}$. $\gamma_{ji}^{\mathrm{NN}} = 1$ if there is a synapse from neuron $j$ to neuron $i$, $\gamma_{ji}^{\mathrm{NN}} = 0$ otherwise. Each pair of neurons (excitatory and inhibitory) is connected with probability $p_{\mathrm{primary}}$. This results in binomial distribution of inputs and outputs: $\mathbf{B}(N^2, p_{\mathrm{primary}})$. |
| E | A | Each astrocyte is associated with a set of synapses and receives inputs from the presynaptic neurons of these synapses according to $\mathbf{\Gamma}_{\mathrm{NA}} = \left\{\gamma_{ji,k}^{\mathrm{NA}}\right\}_{k=1\ldots N_{\mathrm{A}},i=1\ldots N,j=1\ldots N_{\mathrm{E}}}$. $\gamma_{ji,k}^{\mathrm{NA}} = 1$ if an astrocyte receives inputs from a synapse $ji$, $\gamma_{ji,k}^{\mathrm{NA}} = 0$ otherwise. Each generated synapse can be associated with only one selected astrocyte, and the probability of this association is $p_{\mathrm{third\_if\_primary}}$. |
| A | {E,I} | Astrocyte outputs to all postsynaptic neurons of its associated synapses according to $\mathbf{\Gamma}_{\mathrm{AN}} = \{\gamma_{k,ji}^{\mathrm{AN}}\}_{k=1\ldots N_{\mathrm{A}},i=1\ldots N,j=1\ldots N_{\mathrm{E}}}$. $\gamma_{k,ji}^{\mathrm{AN}} = 1$ if the postsynaptic neuron $i$ receives input from the astrocyte $k$, $\gamma_{k,ji}^{\mathrm{AN}} = 0$ otherwise. For each postsynaptic neuron, the astrocyte pool from which to select the associated astrocyte(s) is confined by two parameters $S_{\mathrm{pool}}$ and *pool_type*, which define the pool size and the way to determine the pool, respectively. |

| Connectivity rules | |
|---|---|
| 1) Primary connectivity rule | The primary connectivity rule is the rule to establish primary (neuron-to-neuron) connections with. Possible rules include `pairwise_bernoulli`, `fixed_indegree`, `fixed_outdegree`, and `fixed_total_number`. For specifications of these rules, see Senk et al. (2022) and NEST documentation for connectivity concepts. |
| 2) Third-factor connectivity rule | The third-factor connectivity rule is the rule to establish third-in (neuron-to-astrocyte) and third-out (astrocyte-to-neuron) connections with. `third_factor_bernoulli_with_pool` is the only rule available. For specifications of this rule, see NEST documentation for `TripartiteConnect()` and `third_factor_bernoulli_with_pool`. |

| **Neuron model** | |
|---|---|
| | For each neuron $i = 1 \ldots N$ in the network |
| 1) Subthreshold dynamics | $C_{\mathrm{m}} \cdot \frac{dV_i(t)}{dt} = f(V_i(t)) - W_i(t) + I_{\mathrm{syn},i}(t) + I_{\mathrm{SIC},i}(t) + I_{\mathrm{noise},i}(t)$<br>$f(V_i(t)) = g_{\mathrm{L}} \cdot (E_{\mathrm{L}} - V_i(t)) + g_{\mathrm{L}} \cdot \Delta_{\mathrm{T}} \cdot \exp\left(\frac{V_i(t) - V_{\mathrm{th}}}{\Delta_{\mathrm{T}}}\right)$<br>with membrane capacitance $C_{\mathrm{m}}$, membrane potential $V_i$, adaptation variable $W_i$, summed synaptic current $I_{\mathrm{syn},i}$ from all connected neurons and external inputs (see **Synaptic input for neuron**), summed SIC input $I_{\mathrm{SIC},i}$ from all connected astrocytes, external noise current $I_{\mathrm{noise},i}$ (see **External inputs**), leak conductance $g_{\mathrm{L}}$, leak reversal potential $E_{\mathrm{L}}$, slope factor $\Delta_{\mathrm{T}}$, and spike initiation threshold $V_{\mathrm{th}}$. |
| 2) Adaptation | $\tau_W \cdot \frac{dW_i(t)}{dt} = a \cdot (V_i(t) - E_{\mathrm{L}}) - W_i(t)$<br>with subthreshold adaptation $a$ and adaptation time constant $\tau_W$. |
| 3) Spiking and reset | Spiking condition: $V_i(t_{i,sp}^-) \leq V_{\mathrm{peak}}$ and $V_i(t_{i,sp}) > V_{\mathrm{peak}}$ with spiking threshold $V_{\mathrm{peak}}$ and spike time $t_{i,sp}$ of the $sp$th spike of neuron $i$.<br>If the spiking condition is met then:<br>1. Save the new spike time: $t_{i,sp} \rightarrow \mathcal{S}_i$ ($\mathcal{S}_i$: spikes recorded from neuron $i$)<br>2. Reset: $V_i(t \in [t_{i,sp}, t_{i,sp} + t_{\mathrm{ref}}]) \leftarrow V_{\mathrm{reset}}, W_i(t_{i,sp}^+) \leftarrow W_i(t_{i,sp}^-) + b$<br>with spike-triggered adaptation $b$, refractory period $t_{\mathrm{ref}}$, and voltage reset $V_{\mathrm{reset}}$. |

| **Astrocyte model** | |
|---|---|
| | For each astrocyte $k = 1 ... N_{\mathrm{A}}$ in the network |
| 1) $\mathrm{IP}_3$ | $\frac{d[\mathrm{IP}_3]_k(t)}{dt} = \frac{[\mathrm{IP}_3]_0 - [\mathrm{IP}_3]_k(t)}{\tau_{\mathrm{IP}_3}} + \Delta_{\mathrm{IP}_3} \cdot J_{\mathrm{syn},k}(t)$ <br> with baseline value $[\mathrm{IP}_3]_0$ of $\mathrm{IP}_3$ concentration, time constant $\tau_{\mathrm{IP}_3}$ of the decay of $\mathrm{IP}_3$ concentration, and parameter $\Delta_{\mathrm{IP}_3}$ determining the increase in $\mathrm{IP}_3$ concentration induced by synaptic input $J_{\mathrm{syn},k}$ (see **Synaptic input for astrocyte**). |
| 2) Calcium | $\frac{d[\mathrm{Ca}^{2+}]_k(t)}{dt} = J_{\mathrm{channel},k}(t) - J_{\mathrm{pump},k}(t) + J_{\mathrm{leak},k}(t) + J_{\mathrm{noise},k}(t)$ <br> with $\mathrm{IP}_3\mathrm{R}$-mediated release $J_{\mathrm{channel},k}$ of calcium from ER to cytosol, ATP-dependent pumping $J_{\mathrm{pump},k}$ of calcium from cytosol to ER, leak $J_{\mathrm{leak},k}$ of calcium between ER and cytosol, and random fluctuations $J_{\mathrm{noise},k}$ in calcium dynamics (see **External inputs**). |
| 3) Gating variable for $\mathrm{IP}_3\mathrm{R}$ inactivation | $\frac{dh_{\mathrm{IP}_3\mathrm{R},k}(t)}{dt} = \alpha_{h,k}(t) \cdot (1 - h_{\mathrm{IP}_3\mathrm{R},k}(t)) - \beta_{h,k}(t) \cdot h_{\mathrm{IP}_3\mathrm{R},k}(t)$ <br> $\alpha_{h,k}(t) = k_{\mathrm{IP}_3\mathrm{R}} \cdot K_{\mathrm{d,inh}} \cdot \frac{[\mathrm{IP}_3]_k(t) + K_{\mathrm{d,IP}_{3,1}}}{[\mathrm{IP}_3]_k(t) + K_{\mathrm{d,IP}_{3,2}}}$ <br> $\beta_{h,k}(t) = k_{\mathrm{IP}_3\mathrm{R}} \cdot [\mathrm{Ca}^{2+}]_k(t)$ <br> with fraction $h_{\mathrm{IP}_3\mathrm{R}}$ of $\mathrm{IP}_3\mathrm{Rs}$ that are not yet inactivated by calcium, two rates $\alpha_{h,k}$ and $\beta_{h,k}$ of the exchange between activated and inactivated $\mathrm{IP}_3\mathrm{Rs}$, astrocytic $\mathrm{IP}_3\mathrm{R}$ binding constant $k_{\mathrm{IP}_3\mathrm{R}}$ for calcium inhibition, astrocytic $\mathrm{IP}_3\mathrm{R}$ dissociation constant $K_{\mathrm{d,inh}}$ of calcium (inhibition), and the first and second astrocytic $\mathrm{IP}_3\mathrm{R}$ dissociation constants $K_{\mathrm{d,IP}_{3,1}}$ and $K_{\mathrm{d,IP}_{3,2}}$ of $\mathrm{IP}_3$. |
| 4) Channel, pump and leak currents | $J_{\mathrm{channel},k}(t) = r_{\mathrm{ER,cyt}} \cdot v_{\mathrm{IP}_3\mathrm{R}} \cdot m^3_{\infty,k}(t) \cdot n^3_{\infty,k}(t) \cdot h^3_{\mathrm{IP}_3\mathrm{R},k}(t) \cdot \left([\mathrm{Ca}^{2+}]_{\mathrm{ER},k}(t) - [\mathrm{Ca}^{2+}]_k(t)\right)$ <br> $J_{\mathrm{pump},k}(t) = \frac{v_{\mathrm{SERCA}} \cdot [\mathrm{Ca}^{2+}]^2_k(t)}{K^2_{\mathrm{m,SERCA}} + [\mathrm{Ca}^{2+}]^2_k(t)}$ <br> $J_{\mathrm{leak},k}(t) = r_{\mathrm{ER,cyt}} \cdot v_{\mathrm{L}} \cdot \left([\mathrm{Ca}^{2+}]_{\mathrm{ER},k}(t) - [\mathrm{Ca}^{2+}]_k(t)\right)$ <br> $m_{\infty,k}(t) = \frac{[\mathrm{IP}_3]_k(t)}{[\mathrm{IP}_3]_k(t) + K_{\mathrm{d,IP}_{3,1}}}$ <br> $n_{\infty,k}(t) = \frac{[\mathrm{Ca}^{2+}]_k(t)}{[\mathrm{Ca}^{2+}]_k(t) + K_{\mathrm{d,act}}}$ <br> with ratio $r_{\mathrm{ER,cyt}}$ between astrocytic ER and cytosol volumes, maximum rate $v_{\mathrm{IP}_3\mathrm{R}}$ of calcium release via astrocytic $\mathrm{IP}_3\mathrm{Rs}$, astrocytic $\mathrm{IP}_3\mathrm{R}$ dissociation constant $K_{\mathrm{d,act}}$ of calcium (activation), maximum rate $v_{\mathrm{SERCA}}$ of calcium uptake by astrocytic SERCA pumps, half-activation constant $K_{\mathrm{m,SERCA}}$ of astrocytic SERCA pumps, rate constant $v_{\mathrm{L}}$ of calcium leak from astrocytic ER to cytosol, and steady-state values $m_\infty$ and $n_\infty$ for two gating variables of $\mathrm{IP}_3\mathrm{Rs}$. |
| 5) Calcium concentration in ER | $[\mathrm{Ca}^{2+}]_{\mathrm{ER},k}(t) = \frac{[\mathrm{Ca}^{2+}]_{\mathrm{tot}} - [\mathrm{Ca}^{2+}]_k(t)}{r_{\mathrm{ER,cyt}}}$ <br> with parameter $[\mathrm{Ca}^{2+}]_{\mathrm{tot}}$ determining the maximal cytosolic calcium concentration. |

| **Synaptic input for neuron** | |
|---|---|
| Synaptic current | For each neuron $i = 1...N$ in the network, the summed synaptic current is<br><br>$I_{\text{syn},i}(t) = \sum_j \gamma_{ji}^{\text{NN}} \cdot I_{\text{syn},ji}(t) + I_{\text{Poiss},i}(t)$ |
| 1) from neurons in the network | $I_{\text{syn},ji}(t) = -g_{\text{exc},ji}(t) \cdot (V_i(t) - E_{\text{exc}}) - g_{\text{inh},ji}(t) \cdot (V_i(t) - E_{\text{inh}})$<br><br>with excitatory and inhibitory synaptic conductances $g_{\text{exc},ji}$ and $g_{\text{inh},ji}$, excitatory and inhibitory reversal potentials $E_{\text{exc}}$ and $E_{\text{inh}}$.<br><br>The synaptic conductance is alpha-shaped and modeled with short-term plasticity:<br>$g_{X,ji}(t) = \sum_{sp} w_{X,ji} \cdot u_{ji}(t_{j,sp}) \cdot x_{ji}(t_{j,sp}) \cdot (\frac{t-t_{j,sp}-d_{X,ji}}{\tau_{X,\text{syn}}}) \cdot \exp(1 - \frac{t-t_{j,sp}-d_{X,ji}}{\tau_{X,\text{syn}}})$<br>$\frac{du_{ji}(t)}{dt} = -\frac{u_{ji}(t)}{\tau_{\text{fac}}} + \sum_{sp} U \cdot (1 - u_{ji}(t_{j,sp})) \cdot \delta(t - t_{j,sp} - d_{X,ji})$<br>$\frac{dx_{ji}(t)}{dt} = \frac{z_{ji}(t)}{\tau_{\text{rec}}} - \sum_{sp} u_{ji}(t_{j,sp}) \cdot x_{ji}(t_{j,sp}) \cdot \delta(t - t_{j,sp} - d_{X,ji})$<br>$\frac{dy_{ji}(t)}{dt} = -\frac{y_{ji}(t)}{\tau_{\text{psc}}} + \sum_{sp} u_{ji}(t_{j,sp}) \cdot x_{ji}(t_{j,sp}) \cdot \delta(t - t_{j,sp} - d_{X,ji})$<br>$\frac{dz_{ji}(t)}{dt} = \frac{y_{ji}(t)}{\tau_{\text{psc}}} - \frac{z_{ji}(t)}{\tau_{\text{rec}}}$<br><br>with time constant $\tau_{X,\text{syn}}$ of postsynaptic conductance in the neuron, time constant $\tau_{\text{psc}}$ of postsynaptic current in the short-term plasticity model, weight $w_{X,ji}$ of neuron-to-neuron connection, release probability $u_{ji}$, readily releasable fraction $x_{ji}$ of neurotransmitter, active fraction $y_{ji}$ of neurotransmitter, inactive fraction $z_{ji}$ of neurotransmitter, facilitation time constant $\tau_{\text{fac}}$, depression time constant $\tau_{\text{rec}}$, parameter $U$ determining the increase in $u_{ji}$ with each spike, spike time $t_{j,sp}$ of the $sp$th spike of neuron $j$, delay $d_{X,ji}$ of neuron-to-neuron connection. $X = \text{exc}$ if $j \leq N_{\text{E}}$ and $X = \text{inh}$ if $j > N_{\text{E}}$.<br><br>Synaptic weights $w_{X,ji}$ and delays $d_{X,ji}$ are supplied to the TBP rule via the `primary` synapse specification. |
| 2) from external input | $I_{\text{Poiss},i}(t)$ (see **External inputs**) |
| **Synaptic input for astrocyte** | |
| Synaptic input | For each astrocyte $k = 1...N_{\text{A}}$ in the network, the summed synaptic input is<br><br>$J_{\text{syn},k}(t) = \sum_{ji} \gamma_{ji,k}^{\text{NA}} \cdot J_{\text{syn},ji,k}(t) + J_{\text{Poiss},k}(t)$ |
| 1) from neurons in the network | $J_{\text{syn},ji,k}(t) = w_{\text{pre\_to\_astro},ji,k} \cdot \sum_{sp} u_{ji}(t_{j,sp}) \cdot x_{ji}(t_{j,sp}) \cdot \delta(t - t_{j,sp} - d_{\text{pre\_to\_astro},ji,k})$<br><br>where $t_{j,sp}$ is the time of the $sp$th spike of neuron $j$. The weights $w_{\text{pre\_to\_astro},ji,k}$ and delays $d_{\text{pre\_to\_astro},ji,k}$ of the neuron-to-astrocyte connections are supplied to the TBP rule via the `third_in` synapse specification. |
| 2) from external input | $J_{\text{Poiss},k}(t)$ (see **External inputs**) |

| **Astrocyte-to-neuron current** | |
|---|---|
| Slow inward current (SIC) | For each neuron $i = 1...N$ in the network, the summed SIC input from all connected astrocytes is<br>$I_{\mathrm{SIC},i}(t) = \sum_k \sum_j \gamma^{\mathrm{AN}}_{k,ji} \cdot w_{\mathrm{astro\_to\_post},k,ji} \cdot F_{\mathrm{SIC}}\left([\mathrm{Ca}^{2+}]_k(t - d_{\mathrm{astro\_to\_post},k,ji})\right)$<br>$F_{\mathrm{SIC}}\left([\mathrm{Ca}^{2+}]_k(t)\right) = a_{\mathrm{SIC}} \cdot \Theta\left(\log(c_{\mathrm{scaled},k}(t))\right) \cdot \log\left(c_{\mathrm{scaled},k}(t)\right)$<br>$c_{\mathrm{scaled},k}(t) = \left([\mathrm{Ca}^{2+}]_k(t) - \theta_{\mathrm{SIC}}\right)/\mathrm{nM}$<br>where $a_{\mathrm{SIC}}$ is the scale of SIC output, $\theta_{\mathrm{SIC}}$ the calcium threshold for SIC generation, and $\Theta(x)$ the Heaviside step function. The weights $w_{\mathrm{astro\_to\_post},k,ji}$ and delays $d_{\mathrm{astro\_to\_post},k,ji}$ of the astrocyte-to-neuron connections are supplied to the TBP rule via the `third_out` synapse specification. |

| **External inputs** | |
|---|---|
| Poisson noise input for neurons | For each neuron $i = 1...N$ in the network, a Poisson noise input is applied:<br>$I_{\mathrm{Poiss},i}(t) = -g_{\mathrm{Poiss},i}(t) \cdot (V_i(t) - E_{\mathrm{exc}})$<br>$g_{\mathrm{Poiss},i}(t) = \sum_{sp} w_{\mathrm{Poiss},i} \cdot (\frac{t - t_{\mathrm{Poiss},sp} - d_{\mathrm{Poiss}}}{\tau_{\mathrm{exc,syn}}}) \cdot \exp(1 - \frac{t - t_{\mathrm{Poiss},sp} - d_{\mathrm{Poiss}}}{\tau_{\mathrm{exc,syn}}})$<br>with Poisson input-induced conductance $g_{\mathrm{Poiss},i}$, weight $w_{\mathrm{Poiss},i}$ of input, spike time $t_{\mathrm{Poiss},sp}$ of the $sp$th spike of input, and delay $d_{\mathrm{Poiss}}$ of input. The average firing rate ($\lambda_{\mathrm{Poiss,N},i}$) of the input for each neuron is fixed. |
| Poisson noise input for astrocytes | For each astrocyte $k = 1...N_{\mathrm{A}}$ in the network, a Poisson noise input is applied:<br>$J_{\mathrm{Poiss},k}(t) = w_{\mathrm{Poiss}} \cdot \sum_{sp} \delta(t - t_{\mathrm{Poiss},sp} - d_{\mathrm{Poiss}}),$<br>with weight $w_{\mathrm{Poiss}}$ of input, spike time $t_{\mathrm{Poiss},sp}$ of the $sp$th spike of input, and delay $d_{\mathrm{Poiss}}$ of input. The average firing rate ($\lambda_{\mathrm{Poiss,A}}$) of the input for each astrocyte is fixed. The Poisson noise input affects the $\mathrm{IP}_3$ dynamics in the astrocyte (see **Astrocyte model** and **Synaptic input for astrocyte**). |
| Gaussian noise input for neurons | For each neuron $i = 1...N$ in the network, a Gaussian noise current $I_{\mathrm{noise},i}$ is applied. $I_{\mathrm{noise},i}$ has a fixed mean of zero and variance $\sigma_{\mathrm{Gauss,N}}$. |
| Gaussian noise input for astrocytes | For each astrocyte $k = 1...N_{\mathrm{A}}$ in the network, a Gaussian noise input $J_{\mathrm{noise},k}(t)$ is applied. $J_{\mathrm{noise},k}(t)$ has a fixed mean of zero and variance $\sigma_{\mathrm{Gauss,A}}$. The Gaussian noise input affects the calcium dynamics in the astrocyte (see **Astrocyte model**). |

Table B: Parameters of the neuron-astrocyte network models for the benchmarks in this study. All weights and delays of synaptic interactions are listed. For the other parameters, only those that differ from NEST defaults are listed. For a few parameters, values for different models are listed separately. Otherwise, the value applies to all models. The parameters required for the models but not listed here follow defaults of `astrocyte_lr_1994`, `aeif_cond_alpha_astro`, and `tsodyks_synapse` in NEST.

| Notation | Value | Description |
|---|---|---|
| **Populations** | | |
| $N_{\mathrm{E}}$ | $8,000 \times scale$ | Number of excitatory neurons |
| $N_{\mathrm{I}}$ | $2,000 \times scale$ | Number of inhibitory neurons |
| $N_{\mathrm{A}}$ | $10,000 \times scale$ | Number of astrocytes or surrogates |
| *scale* | 1 (strong-scaling benchmarks)<br>1, 2, 3, 4 (weak-scaling benchmarks)<br>1.25, 2.5, 5, 10, 20, 30, 40, 50 (weak-scaling benchmarks for very large models) | Scale of model |
| **Connectivity** | | |
| `rule` under `conn_spec` | `pairwise_bernoulli` (benchmarks with neuron-astrocyte network models)<br>`pairwise_bernoulli`, `fixed_indegree`, `fixed_outdegree`, `fixed_total_number` (benchmarks with different primary connectivity rules)<br>`pairwise_bernoulli` (weak-scaling benchmarks with different astrocyte pool sizes)<br>`fixed_indegree` (weak-scaling benchmarks for very large models) | The primary connectivity rule |
| `rule` under `third_factor_conn_spec` | `third_factor_bernoulli_with_pool` | The third-factor connectivity rule |
| $p_{\mathrm{primary}}$ | 0.1 / *scale* | Connection probability between neurons |
| $p_{\mathrm{third\_if\_primary}}$ | 0.5 | Probability of each created neuron-to-neuron connection to be paired with one astrocyte |
| $S_{\mathrm{pool}}$ | 10 (benchmarks with neuron-astrocyte network models, benchmarks with different primary connectivity rules)<br>10, 100, 1000, 10000 (weak-scaling benchmarks with different astrocyte pool sizes)<br>10, 1000 (weak-scaling benchmarks for very large models) | The size of astrocyte pool for each target (postsynaptic) neuron |
| *pool_type* | `"random"` | The way to determine the astrocyte pool for each target neuron |
| **Astrocyte parameters** | | |
| $[\mathrm{IP}_3][0]$ | 0.4 µM | Initial $\mathrm{IP}_3$ concentration |
| **`astrocyte_surrogate` parameters (for the "surrogate" model only)** | | |
| SIC output | 3.5 pA | Predefined SIC output |
| **Synaptic interaction parameters** | | |
| $w_{\mathrm{exc}}$ | 1 nS | Weight of excitatory neuron-to-neuron connections |
| $w_{\mathrm{inh}}$ | 4 nS | Weight of inhibitory neuron-to-neuron connections |
| $w_{\mathrm{Poiss}}$ | 1 nS | Weight of excitatory noise-to-neuron and noise-to-astrocyte connections |
| $w_{\mathrm{pre\_to\_astro}}$ | 1 | Weight of excitatory neuron-to-astrocyte connections (the unit is included in the astrocyte parameter $\Delta_{\mathrm{IP}_3}$; see Table A and S2 Appendix) |
| $w_{\mathrm{astro\_to\_post}}$ | 0.05 pA | Weight of astrocyte-to-neuron connections |

| | | |
|---|---|---|
| $d_{\text{exc}}, d_{\text{pre\_to\_astro}}$ | 2 ms | Delay of excitatory neuron-to-neuron and neuron-to-astrocyte connections |
| $d_{\text{astro\_to\_post}}$ | 1 ms | Delay of astrocyte-to-neuron connections |
| $d_{\text{inh}}$ | 1 ms (all except the "Synchronous" model)<br>2 ms (the "Synchronous" model) | Delay of inhibitory neuron-to-neuron connections |
| $d_{\text{Poiss}}$ | 1 ms | Delay of excitatory noise-to-neuron and noise-to-astrocyte connections |
| $\tau_{\text{exc,syn}}$ | 2 ms | Time constant for excitatory connections |
| $\tau_{\text{inh,syn}}$ | 4 ms (all except the "Synchronous" model)<br>2 ms (the "Synchronous" model) | Time constant for inhibitory connections |
| **External input** | | |
| $\lambda_{\text{Poiss,N}}$ | 2000 spikes/s | Rate of the Poisson noise for each neuron |

Table C: Parameters for the models developed to study the impact of neuron-astrocyte interactions on neuronal synchronization. The table lists parameters that were fitted or selected when creating the model, that might differ from the NEST default values. The other parameters required for the models but not listed here follow defaults of `astrocyte_lr_1994`, `aeif_cond_alpha_astro`, and `tsodyks_synapse` in NEST. The same parameters are also listed in the JSON file `neuron_astrocyte_model_parameters.json` in the model implementation code.

| Notation | Value | Description |
|---|---|---|
| **Populations** | | |
| $N_{\mathrm{E}}$ | 400 | Number of excitatory neurons |
| $N_{\mathrm{I}}$ | 100 | Number of inhibitory neurons |
| $N_{\mathrm{A}}$ | 100 | Number of astrocytes |
| **Connectivity** | | |
| $p_{\mathrm{primary}}$ | 0.2 | Connection probability between neurons |
| $p_{\mathrm{third\_if\_primary}}$ | <br>0.2<br>0.03 | Probability of each created neuron-to-neuron connection to be paired with one astrocyte<br>`pool_type` = `"block"`<br>`pool_type` = `"random"` |
| $S_{\mathrm{pool}}$ | <br>1<br>5 | The size of astrocyte pool for each target neuron<br>`pool_type` = `"block"` (1 astrocyte input per neuron)<br>`pool_type` = `"random"` (max. 5 astrocyte inputs per neuron) |
| *pool_type* | `"block"`, `"random"` | The way to determine the astrocyte pool for each target neuron |
| **Neuron parameters** | | |
| $V_{\mathrm{th}}$ | E: $-50$ mV, I: $-52$ mV | Spiking initiation threshold |
| $C_{\mathrm{m}}$ | E: 130 pF, I: 104 pF | Cell membrane capacitance |
| $g_{\mathrm{L}}$ | E: 18 nS, I: 4.3 nS | Leak conductance |
| $E_{\mathrm{L}}$ | E: $-58$ mV, I: $-65$ mV | Resting potential |
| $\Delta_{\mathrm{T}}$ | E, I: 2 mV | Threshold slope factor |
| $\tau_W$ | E: 450 ms, I: 264 ms | Time constant of adaptation current |
| $a$ | E: 4 nS, I: $-0.8$ nS | Adaptation parameter |
| $V_{\mathrm{reset}}$ | Gaussian distribution:<br>E, mean: $\bar{V}_{\mathrm{reset}} = -50$ mV, var: $0.05\bar{V}_{\mathrm{reset}}$<br>allowed range of values: $[0.9\bar{V}_{\mathrm{reset}}, 1.1\bar{V}_{\mathrm{reset}}]$<br>I, mean: $\bar{V}_{\mathrm{reset}} = -60$ mV, var: $0.05\bar{V}_{\mathrm{reset}}$<br>allowed range of values: $[0.9\bar{V}_{\mathrm{reset}}, 1.1\bar{V}_{\mathrm{reset}}]$ | Voltage reset |
| $b$ | Gaussian distribution:<br>E, mean: $\bar{b} = 300$ pA, var: $0.05\bar{b}$<br>allowed range of values: $[0.9\bar{b}, 1.1\bar{b}]$<br>I, mean: $\bar{b} = 130$ pA, var: $0.05\bar{b}$<br>allowed range of values: $[0.9\bar{b}, 1.1\bar{b}]$ | Spike triggered adaptation |

| **Astrocyte parameters** | | |
|---|---|---|
| $[\mathrm{Ca}^{2+}]_{\mathrm{tot}}$ | Gaussian distribution, mean value fitted to the experimental data:<br>mean: $[\bar{\mathrm{Ca}}^{2+}]_{\mathrm{tot}} = 1.8958264782990153$ µM, var: $0.1[\bar{\mathrm{Ca}}^{2+}]_{\mathrm{tot}}$<br>allowed range of values: $[0.9[\bar{\mathrm{Ca}}^{2+}]_{\mathrm{tot}}, 1.05[\bar{\mathrm{Ca}}^{2+}]_{\mathrm{tot}}]$ | Total free calcium concentration in terms of cytosolic volume |
| $[\mathrm{IP}_3]_0$ | Gaussian distribution, mean value fitted to the data:<br>mean: $[\bar{\mathrm{IP}}_3]_0 = 0.010925583871830699$ µM, var: $0.1[\bar{\mathrm{IP}}_3]_0$<br>allowed range of values: $[0.9[\bar{\mathrm{IP}}_3]_0, 1.05[\bar{\mathrm{IP}}_3]_0]$ | Baseline value of $\mathrm{IP}_3$ concentration |
| $[\mathrm{IP}_3][0]$ | 0.010925583871830699 µM | $\mathrm{IP}_3$ concentration is set to be the same as $[\mathrm{IP}_3]_0$ at the start of each simulation |
| $\tau_{\mathrm{IP}_3}$ | Gaussian distribution, mean fitted to the data:<br>mean: $\bar{\tau}_{\mathrm{IP}_3} = 1058.7589568400845$ ms, var: $0.1\bar{\tau}_{\mathrm{IP}_3}$<br>allowed range of values: $[0.9\bar{\tau}_{\mathrm{IP}_3}, 1.05\bar{\tau}_{\mathrm{IP}_3}]$ | Time constant of the exponential decay of astrocytic $\mathrm{IP}_3$ |
| $\Delta_{\mathrm{IP}_3}$ | Gaussian distribution, mean fitted to the data:<br>mean: $\bar{\Delta}_{\mathrm{IP}_3} = 0.04911592657233997$ µM, var: $0.1\bar{\Delta}_{\mathrm{IP}_3}$<br>allowed range of values: $[0.9\bar{\Delta}_{\mathrm{IP}_3}, 1.05\bar{\Delta}_{\mathrm{IP}_3}]$ | Parameter determining the increase in astrocytic $\mathrm{IP}_3$ concentration induced by synaptic input |
| $a_{\mathrm{SIC}}$ | 1 | Scaling coefficient, scales all SICs coming from the same astrocyte |
| **Synaptic interaction parameters (weights)** | | |
| $w_{\mathrm{exc}}$ | 5 nS | Weight of excitatory neuron-to-neuron connections |
| $w_{\mathrm{inh}}$ | 5 nS | Weight of inhibitory neuron-to-neuron connections |
| $w_{\mathrm{Poiss}}$ | 1 nS | Weight of excitatory noise-to-neuron and noise-to-astrocyte connections |
| $d_{\mathrm{exc}}$ | 1 ms | Delay of excitatory neuron-to-neuron connections |
| $d_{\mathrm{inh}}$ | 1 ms | Delay of inhibitory neuron-to-neuron connections |
| $d_{\mathrm{pre\_to\_astro}}$ | 1 ms | Delay of excitatory neuron-to-astrocyte connections |
| $d_{\mathrm{astro\_to\_post}}$ | 1 ms | Delay of astrocyte-to-neuron connections |
| $\tau_{\mathrm{exc,syn}}$ | 2 ms | Time constant for excitatory connections |
| $\tau_{\mathrm{inh,syn}}$ | 2 ms | Time constant for inhibitory connections |
| **Synaptic interaction parameters for neuron-astrocyte interaction** | | |
| $w_{\mathrm{pre\_to\_astro}}$ | Kept equal for all synapses<br>0.2<br>0.31<br>1<br>1.5 | Weight of excitatory neuron-to-astrocyte interaction<br>`pool_type` = `"block"`, asynchronous<br>`pool_type` = `"block"`, network bursting<br>`pool_type` = `"random"`, asynchronous<br>`pool_type` = `"random"`, network bursting (in S3 Appendix) |
| $w_{\mathrm{astro\_to\_post}}$ | Kept equal for all synapses<br>1 pA<br>1 pA<br>7 pA<br>7 pA | Weight of astrocyte-to-neuron connections<br>`pool_type` = `"block"`, asynchronous<br>`pool_type` = `"block"`, network bursting<br>`pool_type` = `"random"`, asynchronous<br>`pool_type` = `"random"`, network bursting (in S3 Appendix) |

| External inputs, noise | | |
|---|---|---|
| $\sigma_{\text{Gauss,N}}$ | 100 pA | Variance of Gaussian noise for neurons |
| $\sigma_{\text{Gauss,A}}$ | 0.0003 $\frac{\mu\text{M}}{\text{ms}}$ | Variance of Gaussian noise for astrocytes. Note that parameter is given in $\frac{\mu\text{M}}{\text{ms}}$ even though the NEST implementation of Gaussian noise usually has the physical unit of pA when used in other models. |
| $\lambda_{\text{Poiss,E}}, \lambda_{\text{Poiss,I}}$ | E: 2700 spikes/s, I: 2500 spikes/s | Rate of the Poisson noise for each neuron |
| $\lambda_{\text{Poiss,A}}$ | 4.3281650042083655 Hz<br><br>3.0297155029458556 Hz | Rate of Poisson noise for each astrocyte (fitted to experimental data)<br><br>Rate of Poisson noise, used in all simulations except the one shown in Fig 9 in the main article (reproducing TTX blocking) |
| **Simulation parameters** | | |
| `pre_sim_time` | 20 s | Preliminary simulation that allows model settle in the steady state regime |
| `sim_time` | 5 min | Simulated model time |
| **Simulation analysis parameters** | | |
| `max_allowed _distance_astro` | 2000 ms | Used when detecting calcium transients |
| `max_allowed _distance_exc`, `max_after_sic_exc` | 2000 ms | Used when detecting single-neuron bursts in excitatory neurons |
| `max_allowed _distance_inh`, `max_after_sic_inh` | 400 ms | Used when detecting single-neuron bursts in inhibitory neurons |
| `hist_window` | 2000 ms | Length of a sliding window for binning spike trains when computing correlation coefficients |
| `hist_shift` | 400 ms | Shift of the sliding window |

### References


R. Brette and W. Gerstner. Adaptive exponential integrate-and-fire model as an effective description of neuronal activity. *Journal of Neurophysiology*, 94:3637–3642, 2005. doi: 10.1152/jn.00686.2005.

G. W. De Young and J. Keizer. A single-pool inositol 1,4,5-trisphosphate-receptor-based model for agonist-stimulated oscillations in $Ca^{2+}$ concentration. *Proceedings of the National Academy of Sciences of the United States of America*, 89(20):9895–9899, 1992. doi: 10.1073/pnas.89.20.9895.

Y.-X. Li and J. Rinzel. Equations for $InsP_3$ receptor-mediated $[Ca^{2+}]_i$ oscillations derived from a detailed kinetic model: a Hodgkin-Huxley like formalism. *Journal of Theoretical Biology*, 166(4):461–473, 1994. doi: 10.1006/jtbi.1994.1041.

S. Nadkarni and P. Jung. Spontaneous oscillations of dressed neurons: a new mechanism for epilepsy? *Physical Review Letters*, 91(26):268101, 2003. doi: 10.1103/PhysRevLett.91.268101.

R. Naud, M. Marcille, C. Clopath, and W. Gerstner. Firing patterns in the adaptive exponential integrate-and-fire model. *Biological Cybernetics*, 99:335–347, 2008. doi: 10.1007/s00422-008-0264-7.

E. Nordlie, M.-O. Gewaltig, and H. E. Plesser. Towards reproducible descriptions of neuronal network models. *PLoS Computational Biology*, 5(8):e1000456, 2009. doi: 10.1371/journal.pcbi.1000456.

T. M. Pirttimaki, R. E. Sims, G. Saunders, S. A. Antonio, N. K. Codadu, and H. R. Parri. Astrocyte-mediated neuronal synchronization properties revealed by false gliotransmitter release. *Journal of Neuroscience*, 37(41):9859–9870, 2017. doi: 10.1523/JNEUROSCI.2761-16.2017.

J. Senk, B. Kriener, M. Djurfeldt, N. Voges, H.-J. Jiang, L. Schüttler, G. Gramelsberger, M. Diesmann, H. E. Plesser, and S. J. van Albada. Connectivity concepts in neuronal network modeling. *PLoS Computational Biology*, 18(9): e1010086, 2022. doi: 10.1371/journal.pcbi.1010086.

M. Tsodyks, K. Pawelzik, and H. Markram. Neural networks with dynamic synapses. *Neural Computation*, 10(4): 821–835, 1998. doi: 10.1162/089976698300017502.

M. Tsodyks, A. Uziel, and H. Markram. Synchrony Generation in Recurrent Networks with Frequency-Dependent Synapses. *Journal of Neuroscience*, 20(1):RC50, 2000. doi: 10.1523/JNEUROSCI.20-01-j0003.2000.
