## Supplementary material for "Modeling neuron-astrocyte interactions in neural networks using distributed simulation": S2 Appendix

### S2 Appendix: Variables and parameters added to the NEST support for astrocytes

Here, we provide additional details of new developed NEST support for astrocytes to supplement the description given in Methods of the article. The new development implements a model for astrocytic calcium dynamics derived by Li and Rinzel (1994) as a reduction of De Young and Keizer's model (De Young and Keizer, 1992) for calcium in non-excitable cells. The model for astrocyte-neuron interaction through slow inward current (SIC) was published by Nadkarni and Jung (2003). These models have been frequently used in later publications of neuron-astrocyte network models (see Manninen et al., 2018, 2023). The main software components supporting the respective model are illustrated in Fig 1. These components are mainly explained in the article and S1 Appendix, and only partly in this appendix.

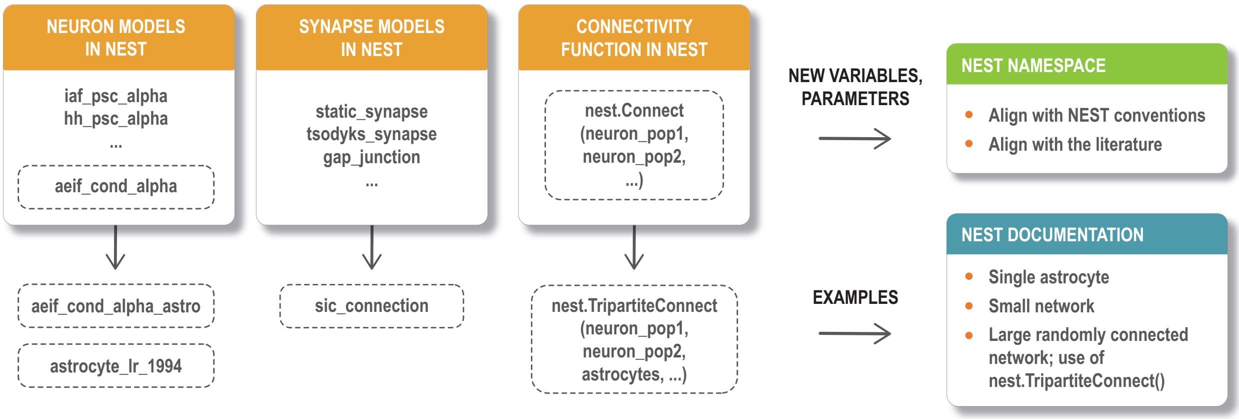


Figure 1: Schematic representation of the software components required for the representation of the models and their embedding in the NEST code. The orange blocks illustrate the added astrocyte-specific models derived from the previously existing NEST models; the old models are listed in boxes, the added models are below the boxes. The green box to the right shows efforts in aligning naming conventions. The blue box below shows examples added to the NEST documentation.

The rest of this appendix describes alignment of naming conventions illustrated in Fig 1 (green box). Tables A-C compare the parameter and variable names in NEST, the mathematical expressions used in this article, and the mathematical expressions used in publications (Li and Rinzel, 1994; Nadkarni and Jung, 2003) and frequently in later network modeling studies (Manninen et al., 2018, 2023). In addition, the NEST default values (for parameters), the NEST default initial conditions (for variables), the physical units, and description of each variable and parameter are given. **Table A** lists parameters for the new connectivity function `TripartiteConnect()`. **Tables B** and **C** list variables and parameters for the astrocyte model `astrocyte_lr_1994`, respectively. These variables and parameters are harmonized with NEST naming conventions, and thus they differ from the notation used by Li and Rinzel (1994) and Nadkarni and Jung (2003).

#### References

G. W. De Young and J. Keizer. A single-pool inositol 1,4,5-trisphosphate-receptor-based model for agonist-stimulated oscillations in $Ca^{2+}$ concentration. *Proceedings of the National Academy of Sciences of the United States of America*, 89(20):9895–9899, 1992. doi: 10.1073/pnas.89.20.9895.

Y.-X. Li and J. Rinzel. Equations for $InsP_3$ receptor-mediated $[Ca^{2+}]_i$ oscillations derived from a detailed kinetic model: a Hodgkin-Huxley like formalism. *Journal of Theoretical Biology*, 166(4):461–473, 1994. doi: 10.1006/jtbi.1994.1041.

T. Manninen, R. Havela, and M.-L. Linne. Computational models for calcium-mediated astrocyte functions. *Frontiers in Computational Neuroscience*, 12:14, 2018. doi: 10.3389/fncom.2018.00014.

T. Manninen, J. Aćimović, and M.-L. Linne. Analysis of network models with neuron-astrocyte interactions. *Neuroinformatics*, 21:375–406, 2023. doi: 10.1007/s12021-023-09622-w.

S. Nadkarni and P. Jung. Spontaneous oscillations of dressed neurons: a new mechanism for epilepsy? *Physical Review Letters*, 91(26):268101, 2003. doi: 10.1103/PhysRevLett.91.268101.

Table A: Parameters of the connectivity function `TripartiteConnect()`.

| Parameter in NEST | For mathematical descriptions in this article | Default value in NEST | Description |
|---|---|---|---|
| `conn_spec` | - | - | Parameters that specify the primary connectivity, i.e., connections between nodes in the two primary populations. See `conn_spec` in the NEST documentation for `Connect()`. |
| `third_factor_conn_spec` | - | - | Parameters that specify the third-factor connectivity, i.e., connections between primary nodes and nodes in the third population. |
| `syn_specs` | - | - | Parameters that specify the synaptic interactions in the primary and the third-factor connectivity. |
| `third_factor_conn_spec` parameters | | | |
| `rule` | - | - | The third-factor connectivity rule. At present, `third_factor_bernoulli_with_pool` is the only rule available. |
| `p` | $p_{\text{third\_if\_primary}}$ | - | Probability of each created primary connection to be paired with one third-factor node. |
| `pool_size` | $S_{\text{pool}}$ | Total number of third-factor nodes (if `pool_type` = `"random"`)<br>- (if `pool_type` = `"random"`) | The size of third-factor node pool for each target (postsynaptic) primary node. Each target primary node only receives inputs from the third-factor nodes in this pool. |
| `pool_type` | *pool_type* | `"random"` | The way to determine the third-factor node pool for each target primary node. If `"random"`, a number (`pool_size`) of third-factor nodes are randomly chosen from all third-factor nodes (without replacement) and assigned as the pool. If `"block"`, the third-factor nodes are evenly distributed to the primary nodes in blocks without overlapping, and the specified `pool_size` has to be compatible with this arrangement. |
| `syn_specs` parameters | | | |
| | | | Specifications of the synaptic interaction (model, weight, delay, etc.): |
| `primary` | - | - | – between primary nodes |
| `third_in` | - | - | – from primary nodes to third-factor nodes |
| `third_out` | - | - | – from third-factor nodes to primary nodes |
| | | | Each follows the style of `syn_specs` in NEST. See `syn_specs` in the NEST documentation for `Connect()`. |

Table B: Variables of the astrocyte model `astrocyte_lr_1994`.

| Variable in NEST | This article | Original publication | Default initial condition in NEST | Physical unit | Description |
|---|---|---|---|---|---|
| `IP3` | $[\mathrm{IP}_3]$ | $[\mathrm{IP}_3]$ | 0.16 | µM | $\mathrm{IP}_3$ concentration in the astrocytic cytosol |
| `Ca_astro` | $[\mathrm{Ca}^{2+}]$ | $[\mathrm{Ca}^{2+}]$ | 0.073 | µM | Calcium concentration in the astrocytic cytosol |
| `h_IP3R` | $h_{\mathrm{IP}_3\mathrm{R}}$ | $q$ | 0.793 | - | Fraction of $\mathrm{IP}_3$ receptors on the astrocytic ER that are not yet inactivated by calcium |

Table C: Parameters of the astrocyte model astrocyte_lr_1994.

| Parameter in NEST | This article | Original publication | Default value in NEST | Physical unit | Description |
|---|---|---|---|---|---|
| Ca_tot | $[\mathrm{Ca}^{2+}]_{\mathrm{tot}}$ | $c_0$ | 2 | µM | Total free calcium concentration in terms of cytosolic volume |
| IP3_0 | $[\mathrm{IP}_3]_0$ | $[\mathrm{IP}_3]^*$ | 0.16 | µM | Baseline value of astrocytic $\mathrm{IP}_3$ concentration |
| Kd_act | $K_{\mathrm{d,act}}$ | $d_5$ | 0.08234 | µM | Astrocytic $\mathrm{IP}_3\mathrm{R}$ dissociation constant of calcium (activation) |
| Kd_inh | $K_{\mathrm{d,inh}}$ | $d_2$ | 1.049 | µM | Astrocytic $\mathrm{IP}_3\mathrm{R}$ dissociation constant of calcium (inhibition) |
| Kd_IP3_1 | $K_{\mathrm{d,IP_3,1}}$ | $d_1$ | 0.13 | µM | First astrocytic $\mathrm{IP}_3\mathrm{R}$ dissociation constant of $\mathrm{IP}_3$ |
| Kd_IP3_2 | $K_{\mathrm{d,IP_3,2}}$ | $d_3$ | 0.9434 | µM | Second astrocytic $\mathrm{IP}_3\mathrm{R}$ dissociation constant of $\mathrm{IP}_3$ |
| Km_SERCA | $K_{\mathrm{m,SERCA}}$ | $k_3$ | 0.1 | µM | Half-activation constant of astrocytic SERCA pump |
| ratio_ER_cyt | $r_{\mathrm{ER,cyt}}$ | $c_1$ | 0.185 | - | Ratio between astrocytic ER and cytosol volumes |
| delta_IP3 | $\Delta_{\mathrm{IP_3}}$ | $r_{\mathrm{IP_3}}$ | 0.0002 | µM | Parameter determining the increase in astrocytic $\mathrm{IP}_3$ concentration induced by synaptic input |
| k_IP3R | $k_{\mathrm{IP_3R}}$ | $a_2$ | 0.0002 | $\frac{1}{\mu\mathrm{M\ ms}}$ | Astrocytic $\mathrm{IP}_3\mathrm{R}$ binding constant for calcium inhibition |
| rate_L | $v_{\mathrm{L}}$ | $v_2$ | 0.00011 | $\frac{1}{\mathrm{ms}}$ | Rate constant of calcium leak from astrocytic ER to cytosol |
| rate_IP3R | $v_{\mathrm{IP_3R}}$ | $v_1$ | 0.006 | $\frac{1}{\mathrm{ms}}$ | Maximum rate of calcium release via astrocytic $\mathrm{IP}_3\mathrm{R}$ |
| rate_SERCA | $v_{\mathrm{SERCA}}$ | $v_3$ | 0.0009 | $\frac{\mu\mathrm{M}}{\mathrm{ms}}$ | Maximum rate of calcium uptake by astrocytic SERCA pump |
| tau_IP3 | $\tau_{\mathrm{IP_3}}$ | $\tau_{\mathrm{IP_3}}$ | 7142 | ms | Time constant of the exponential decay of astrocytic $\mathrm{IP}_3$ |
| SIC_th | $\theta_{\mathrm{SIC}}$ | 0.19669 | 0.19669 | µM | Threshold that determines the minimal level of astrocytic cytosolic calcium sufficient to induce SIC |
| SIC_scale | $a_{\mathrm{SIC}}$ | 2.11 | 1 | - | Parameter determining the scale of astrocytic SIC output |
