## Supplementary material for "Modeling neuron-astrocyte interactions in neural networks using distributed simulation": S3 Appendix

### S3 Appendix: Details of construction of use case model including parameter selection. Additional data supporting the findings about astrocytes' role in neuronal synchrony.

**Han-Jia Jiang, Jugoslava Aćimović, Tiina Manninen, Iiro Ahokainen, Jonas Stapmanns, Mikko Lehtimäki, Markus Diesmann, Sacha J. van Albada, Hans Ekkehard Plesser, and Marja-Leena Linne**

Here, we describe in detail the process of constructing and fitting computational model of neuron-astrocyte population motivated by publication by Pirttimaki et al. (2017). The model is used to examine the influence of neuron-astrocyte interaction on neuronal synchronization and to demonstrate capabilities of the new NEST support for astrocyte modeling. The selection of model parameters is documented in detail; a more compact description is also given in Methods of the article. The selected parameters are systematically listed in S1 Appendix.

Tables A and B summarize experimental data, collected from publications of others, used to fine tune the model parameters. The tables contain a) name of the measured quantity, b) data shown in a publication, c) physical unit, d) experimental setup (model animal, type of preparation, brain region etc.), and e) publication where the data is taken from. Astrocyte and neuron models are fitted according to this data, as described in what follows.

Fig 1 illustrates responses of the selected neuron models to a SIC-mimicking current. Figs 2 and 3 illustrate connectivity in neuron-astrocyte networks with `pool_type` = `"random"` and $p_{\mathrm{third\_if\_primary}} = 0.03$ and 0.05, respectively. Results for networks with this connectivity are shown in Results of the article (for $p_{\mathrm{third\_if\_primary}} = 0.03$), and in the supplementary Fig 6 given in this appendix.

Finally, this appendix contains additional simulations that validate and extend the results presented in Results of the article. Fig 4 shows the model with `pool_type` = `"block"` and asynchronous network activity for a different seed of the random number generator, confirming that results do not depend on the noise properties. Fig 5 shows that results persist when probability of interaction between neurons and astrocytes, i.e. the parameter $p_{\mathrm{third\_if\_primary}}$, is increased from 0.03 to 0.05 in model with `pool_type` = `"random"` and asynchronous activity. Fig 6 illustrates network synchrony/bursting in a model with random connectivity, model with the same connectivity scheme and asynchronous network activity is shown in Results of the article. Figs 7 and 8 confirm that simulation results remain stable when synaptic delay is reduced from (default) 1 ms to 0.1 ms. Figs 9 and 10 confirm long-term stability of results; the models simulated for 20 min show consistent results with the models simulated for 5 minutes. Figs 11, 12, and 13 confirm that the choice of the maximal step size in integration scheme (Runge-Kutta-Fehlberg-45) does not affect the conclusions. Figs 11 and 12 repeat some of the results for the maximal step size set to 0.01 ms and Fig 13 to 0.2 ms; in all other simulations it is fixed to 0.1 ms, the default value in NEST.

#### Fitting the astrocyte model to reproduce dynamics of spontaneous calcium transients from the literature

**Step 1 (Experimental data):** In order to reproduce realistic dynamics of astrocytic calcium transients, we first collect their quantitative characterizations from the literature and summarize them in Table A. Calcium transients are typically described by frequency and duration of transients as well as the maximal change of calcium level during transients. We collect these values from several studies that measured calcium levels in astrocytes, and also from the study by Pirttimaki et al. (2017) who recorded slow inward currents (SICs) in neurons induced by calcium transients in proximal astrocytes. Based on the values in Table A, the realistic ranges of values are assumed to be the following:

- **Frequency:** 0.15–1.5 transients/minute.
- **Duration:** 1–20 s.
  Duration of calcium transients might vary a lot in the literature. If both single and multi-peak calcium transients

are included, then variability of transients can be substantial. We focus on modeling single-peak transients, and thus, the time course, frequency, and duration of the simulated transients correspond to the single-peak case.

- **Maximal calcium level:** Calcium transients are established by measuring relative increase in the calcium level compared to the baseline value. The values vary as much as from 0.1 to 10-fold increase. For modeling purposes, we need to select the feasible range of *absolute values* for calcium concentration. A transient is detected whenever calcium level exceeded the default threshold in the astrocyte model by Nadkarni and Jung (2003), also set to be the default threshold in NEST, $\theta_{\mathrm{SIC}} = 0.19669$ µM + 1 nM $\approx \theta_{\mathrm{SIC}}$. We consider that the peak calcium level within transients should be within $[\theta_{\mathrm{SIC}}, 1]$ µM.

**Step 2 (Fitting):** In order to select a model that reproduces the desired astrocyte dynamics, $10^5$ randomly selected models are simulated. For each model, five parameters are randomly selected from the predefined intervals, while other parameters are kept at NEST default values. These parameters and their predefined intervals are the following.

- $[\mathrm{Ca}^{2+}]_{\mathrm{tot}} \in [0.5, 4]$ µM
- $[\mathrm{IP}_3]_0 \in [0.01, 1]$ µM
- $\tau_{\mathrm{IP}_3} \in [100, 2000]$ ms
- $\Delta_{\mathrm{IP}_3} \in [0.001, 0.1]$ µM
- $\lambda_{\mathrm{Poiss,A}} \in [1, 1000]$ spikes/s

An isolated astrocyte receives Poisson input that mimics spontaneous noisy inputs through $\mathrm{IP}_3$ receptors. The frequency of these inputs $\lambda_{\mathrm{Poiss,A}}$ is one of the fitted parameters.

Each model is simulated for 10 min of model time. In order to increase reliability of results, each simulation is repeated five times, for five different seeds of the random number generator. All models that show at least three calcium transients in each of five simulations are saved for latter. This step only removes models where calcium levels do not depart from the baseline value, or quickly increase to a persistent high level.

**Step 3 (Selection):** Next, all models selected in the previous step are examined semi-manually in three steps:

- Plot model dynamics for visual inspection.
- Simulate model for five random seeds. Count how many times frequency, duration, and peak amplitude of each simulation fall within the range of values from Step 1. Ideally, all three values should fall within the desired range in all five simulations.
- Test model sensitivity. Change one of the five fitted parameters for ±1%, ±5% or ±10% of its original value. Check if the simulated calcium transients still follow the expected dynamics. Do this for each of the five parameters. Sensitivity analysis provided the intervals for parameter values within which the astrocyte model retains the desired dynamics. In the final model for neuron-astrocyte population, these five fitted parameters are randomly drawn from a Gaussian distribution on the intervals selected through sensitivity analysis. However, it should be noted that changing parameters for ±5% or more often lead to the lost of desired dynamics in some of the five repetitions. Thus, we opt to keep model parameters within a narrow range of 90% to 105% of the fitted parameter value.

The final selected set of astrocyte parameters is listed in S1 Appendix and also in the JSON file supplied with Python implementation of the model.

Table A: Quantitative measures from the literature used to fit astrocyte model parameters ($\lambda_{\mathrm{Poiss,A}}$, $\Delta_{\mathrm{IP_3}}$, $[\mathrm{Ca}^{2+}]_{\mathrm{tot}}$, $[\mathrm{IP_3}]_0$, $\tau_{\mathrm{IP_3}}$). Measures represent spontaneous astrocyte activity recorded under various experimental preparations. They are used to tune astrocyte calcium dynamics to model *in silico* spontaneous calcium transients.

| Experimentally measured quantity | Value | Physical unit | Experimental preparation | Original publication describing the measurement |
|---|---|---|---|---|
| SIC frequency | $1.26 \pm 0.2$ | SIC/min | Rat (P10–P16, P19–P21); thalamocortical, hippocampal, somatosensory slices | Pirttimaki et al. (2017): Fig 1A |
| Frequency of $[\mathrm{Ca}^{2+}]$ transients | 0.9–1.5 (soma), 0.4–0.5 (process) | events/min/ROI | Mouse (P46–P67); hippocampal slices | Srinivasan et al. (2015): Fig 1D |
| Frequency of $[\mathrm{Ca}^{2+}]$ transients | 0.1–0.2 | transients/min | Mouse (P9–P12); hippocampal slices | Sasaki et al. (2011): Fig 1F |
| Frequency of $[\mathrm{Ca}^{2+}]$ transients | 0.4 (soma), 0.64 (process) | events/min/ROI | Mouse (8–10 weeks old); somatosensory cortex, layer 1 and 2/3, *in vivo* | Stobart et al. (2018): Fig 1D |
| Frequency of $[\mathrm{Ca}^{2+}]$ transients | $\sim$ 1–20 | events/min | Mouse (P5–P7); organotypic hippocampal slices | Arizono et al. (2020): Fig 7H–K |
| SIC rise time | $172.4 \pm 7.8$ | ms | Rat (P10–P16, P19–P21); thalamocortical, hippocampal, somatosensory slices | Pirttimaki et al. (2017): Fig 1B |
| SIC decay time | $729.8 \pm 40.8$ | ms | Rat (P10–P16, P19–P21); thalamocortical, hippocampal, somatosensory slices | Pirttimaki et al. (2017): Fig 1B |
| SIC duration (rise + decay time) | $902.2 \pm 48.6$ | ms | Rat (P10–P16, P19–P21); thalamocortical, hippocampal, somatosensory slices | Pirttimaki et al. (2017): Fig 1B |
| Spontaneous astrocytic $[\mathrm{Ca}^{2+}]$ duration | 3–10; half-width: $\sim$ 3.3 (soma), 1.5–3 (process) | s | Mouse (P46–67); hippocampal slices | Srinivasan et al. (2015): Fig 1F |
| Spontaneous astrocytic $[\mathrm{Ca}^{2+}]$ duration | 1–20; median 4 | s | Mouse (P5–P7); organotypic hippocampal slices | Arizono et al. (2020): Fig 6G |
| Duration of $[\mathrm{Ca}^{2+}]$ transients | 9–18 (single, multi peaks) | s | Mouse (8–10 weeks old); somatosensory cortex, layer 1 and 2/3, *in vivo* | Stobart et al. (2018): Fig 1F,J |
| Duration of $[\mathrm{Ca}^{2+}]$ transients | 10–15 | s | Mouse (P9–P12); hippocampal slices | Sasaki et al. (2011): Fig 1G |

| **Experimentally measured quantity** | **Value** | **Physical unit** | **Experimental preparation** | **Original publication describing the measurement** |
|---|---|---|---|---|
| Increase of [$Ca^{2+}$] level during transient w.r.t. baseline level | 3 (soma), 2.5–5.5 (process) | $dF/F_0$ | Mouse (P46–P67); hippocampal slices | Srinivasan et al. (2015): Fig 1E |
| Increase of [$Ca^{2+}$] level | 2–6; median 3.16 | $dF/F_0$ | Mouse (P5–P7); organotypic hippocampal slices | Arizono et al. (2020): Fig 6F |
| Increase of [$Ca^{2+}$] level | 0.1 (soma), 0.35 (process) | $dF/F_0$ | Mouse (8–10 weeks old); somatosensory cortex, layer 1 and 2/3, *in vivo* | Stobart et al. (2018): Fig 1E |
| Increase of [$Ca^{2+}$] level | 5–15 | $dF/F_0$ | Mouse (P9–P12); hippocampal slices | Sasaki et al. (2011): Fig 1H |
| SIC amplitude | $243 \pm 7.3$ | pA | Rat (P10–P16, P19–P21); thalamocortical, hippocampal, somatosensory slices | Pirttimaki et al. (2017): Fig 1A |

#### Selecting neuron model parameters

Neurons are modeled as standard spiking neurons with adaptation (AdEx neurons, Naud et al., 2008; Touboul and Brette, 2008). AdEx model neuron has been extensively analyzed in the literature and is implemented in NEST. The experimental study (Pirttimaki et al., 2017), guiding here presented *in silico* work, does not offer much data to constrain the neuron model. Furthermore, in the study by Pirttimaki et al. (2017) neuronal dynamics is examined and neuronal synchrony is demonstrated in recordings of neuronal calcium activity, a signal that is not explicitly represented in AdEx model. However, Fig 5 in that article confirms simultaneous emergence of neuronal bursts and neuronal calcium transients in response to an input SIC. Relying on this result, we model neuronal response to SIC as an emergence of neuronal burst, and we study neuronal dynamics and synchronization by evaluating appropriate metrics from neuronal bursts.

In the study by Pirttimaki et al. (2017), a single neuron responds to an input SIC with a burst of about 2–20 spikes lasting for a couple of seconds. This is used as the only constraint to select parameters of the neuron model. Naud et al. (2008) presents detailed analysis of the AdEx parameter space and gives a handful of model parameters that support specific spiking patterns. We adapt the parameters from Table 1 of Naud et al. (2008) to create neurons that respond to a SIC-mimicking current with spike numbers that fall within the experimentally observed range of 2–20 spikes (Fig 1). **Excitatory neurons** are based on the initial bursters (Table 1, Fig 4c in Naud et al., 2008), i.e. when injected with a suprathreshold current they respond with a short intensive burst which is then followed by several spikes with progressively longer inter-spike intervals. **Inhibitory neurons** are based on the regular spiking type (Table 1, Fig 8c in Naud et al., 2008), which has more consistent inter-spike intervals than the initial bursters. This way, inhibitory neurons are allowed to generate more spikes per input signal, which could help to account for higher average firing rates of inhibitory neurons seen in the literature (Table B). Parameters $\tau_w$, $\bar{V}_{\mathrm{reset}}$, and $\bar{b}$ are modified with respect to the values given by Naud et al. (2008) to achieve the required number of spikes per burst. Also, $\Delta_{\mathrm{T}}$ is set to 2 ms for both neuron types as it does not affect the spiking pattern significantly in this case.

To achieve variability of spiking patterns across neurons, two parameters that substantially affect adaptation and inter-spike intervals, namely $V_{\mathrm{reset}}$ and $b$, are randomized. In the final network model these two parameters are drawn from Gaussian distribution on the interval 90% to 110% around the selected value. In addition, both neuron types receive Poisson and Gaussian noise.

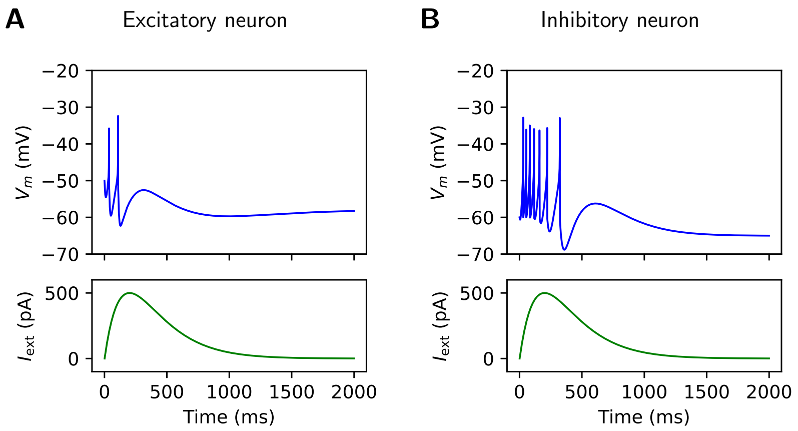


Figure 1: Responses of the selected neuron models to a SIC-mimicking current. A) Excitatory neuron. The parameters are adapted from an initial bursting neuron model (Table 1, Fig 4c in Naud et al., 2008). Here, the adaptive time constant $\tau_w = 450$ ms, $b = 300$ pA, and other parameters are the same as the original model. B) Inhibitory neuron. The parameters are adapted from a regular spiking neuron model (Table 1, Fig 8c in Naud et al., 2008). Here, the slope factor $\Delta_{\mathrm{T}} = 2$ mV, $\tau_w = 264$ ms, $V_{\mathrm{reset}} = -60$ mV, $b = 130$ pA, and other parameters are the same as the original model. $V_{\mathrm{m}}$: membrane potential of the neuron. $I_{\mathrm{ext}}$: SIC-mimicking current applied to the neuron. This current is created with an alpha-shaped function and an amplitude of 500 pA to mimic the SIC current from Fig 5D by Pirttimaki et al. (2017).

Table B: Quantitative measures from the literature used to select neuron model parameters ($a$, $b$, $C_\mathrm{m}$, $\Delta_\mathrm{T}$, $g_\mathrm{L}$, $E_\mathrm{L}$, $V_\mathrm{th}$, $V_\mathrm{reset}$, $\tau_w$ in AdEx neuron model) and synaptic weights, and to tune frequency and spiking patterns of model neurons.

| Experimentally measured quantity | Value | Physical unit | Experimental preparation | Original publication describing the measurement |
|---|---|---|---|---|
| EPSC amplitude | $392 \pm 83.68$ | pA | Rat (P10–P16, P19–P21); thalamocortical, hippocampal, somatosensory slices | Pirttimaki et al. (2017): Fig 2D |
| Spike pattern | Bursting, $\sim$ 2–20 spikes per burst | - | Rat (P10–P16, P19–P21); thalamocortical, hippocampal, somatosensory slices | Pirttimaki et al. (2017): Fig 5G,E |
| Spontaneous firing rate of excitatory neurons *in vivo* | 0.18–10 | spikes/s | Rat; somatosensory (awake), auditory (anesthetized), and visual (awake, sleep) cortex *in vivo* | Maksimov et al. (2018): Table 1 |
| Spontaneous firing rates of neurons *in vivo* | **excitatory:** $1.0\pm0.3$ (quiet), $1.0\pm0.3$ (whisking), **fast spiking:** $10.6 \pm 2.1$ (quiet), $4.2 \pm 1.4$ (whisking), **non-fast-spiking:** $2.2 \pm 0.4$ (quiet), $4.5 \pm 0.9$ (whisking) | spikes/s | Mouse; L2/3 barrel cortex *in vivo* (quiet or whisking) | Gentet et al. (2010) |
| Spontaneous firing rates of neurons *in vivo* | **excitatory:** $0.4 \pm 0.1$, **fast spiking (GABAergic):** $9.4 \pm 2.1$, **SOM (GABAergic):** $6.3 \pm 0.6$, **non-fast-spiking (GABAergic):** $3.7 \pm 0.7$ | spikes/s | Mouse; L2/3 barrel cortex *in vivo* (quiet) | Gentet et al. (2012) |

#### Selecting connectivity weights

The final part of the model construction is the selection of synaptic weight parameters $w_{\mathrm{exc},ji}$ (excitatory synapses), $w_{\mathrm{inh},ji}$ (inhibitory synapses), $w_{\mathrm{pre\_to\_astro},ji,k}$ (glutamatergic input from excitatory neurons to astrocytes), as well as $w_{\mathrm{astro\_to\_post},k,ji}$ (SIC current from astrocytes to neurons). In this model, all weights of the same type are kept equal, i.e. $\forall i,j \in \{1..N\}$ and $\forall k \in \{1..N_{\mathrm{A}}\}$ the weights are $w_{\mathrm{exc},ji} = w_{\mathrm{exc}}$, $w_{\mathrm{inh},ji} = w_{\mathrm{inh}}$, $w_{\mathrm{pre\_to\_post},ji,k} = w_{\mathrm{pre\_to\_astro}}$ and $w_{\mathrm{astro\_to\_post},k,ji} = w_{\mathrm{astro\_to\_post}}$.

**Neuronal network:** We first select weights of the neuron-neuron connections together with the rate of Poisson inputs to excitatory and inhibitory neurons. As a guidance for selection of these parameters, we compile a list of experimentally measured spiking rates of excitatory and inhibitory cells from the literature. The list is given in Table B. According to this list, excitatory neurons spike at the rate of 0.18–1 spikes/s but in some cases up to 10 spikes/s. Inhibitory neurons spike at roughly 2–10 spikes/s. Assuming that SIC from astrocytes increases neuronal spiking rate, we set the parameters of the neuronal network in such a way that both neuronal types show spiking rate at the lower end of the biologically plausible interval. Furthermore, the neuron-only network shows asynchronous low-firing activity regime.

This regime is obtained for $\lambda_{\mathrm{Poiss,E}} = 2700$ spikes/s, $\lambda_{\mathrm{Poiss,I}} = 2500$ spikes/s, $w_{\mathrm{exc}} = 5$ nS, and $w_{\mathrm{inh}} = -5$ nS. Same as before, the rate of Poisson noise is equal for all neurons of the same type and the synaptic weights are identical for all synapses of the same type, thus indices are omitted.

**Neuron-astrocyte interaction** is determined by four model parameters, two parameters for the connectivity scheme and two weight coefficients. We consider two (out of three possible, see Results of the article) connectivity schemes:

- `pool_type = "block"` with $S_{\mathrm{pool}} = 1$. Here the astrocyte domains, in our model the groups of neurons receiving SIC from the same astrocyte, are non-overlapping and of fixed size, the size being determined by ratio between the number of astrocytes and the number of neurons. In our model, each astrocyte sends SIC to 4 excitatory and 1 inhibitory neuron.
- `pool_type = "random"`. Here, each postsynaptic neuron receives SIC current from at most $S_{\mathrm{pool}}$ astrocytes. However, the number and selection of neurons that interact with an astrocyte is random, defined by two connectivity parameters $p_{\mathrm{primary}}$, the probability of synaptic connection between two neurons, and $p_{\mathrm{third\_if\_primary}}$, the probability of synapse interacting with an astrocyte.

The third connectivity option available in NEST astrocyte module, `pool_type = "block"` with $S_{\mathrm{pool}} > 1$, allows SIC inputs from multiple astrocytes coming to the same postsynaptic neurons, but not multiple postsynaptic neurons receiving inputs from the same astrocyte (see illustrations of tripartite connectivity in Fig 2 in the main article). Therefore, this option does not allow to evaluate synchronization in groups of neurons interacting with the same astrocyte.

For both of these connectivity schemes, the probability of neuron-neuron connection is $p_{\mathrm{primary}} = 0.2$ and the number of astrocytes and neurons is the same, $N_{\mathrm{A}} = 100$, $N_{\mathrm{E}} = 400$, and $N_{\mathrm{I}} = 100$.

We tune parameters $w_{\mathrm{astro\_to\_post}}$ and $w_{\mathrm{pre\_to\_astro}}$ separately for each of these connectivity schemes. We first set $w_{\mathrm{pre\_to\_astro}} = 0$ and test the range of parameter $w_{\mathrm{astro\_to\_post}}$ that can induce single-neuron bursts in response to SICs. Here, SICs result from spontaneous astrocytic transients induced by Poisson noise. Next, we set $w_{\mathrm{astro\_to\_post}} = 0$ and also $\lambda_{\mathrm{Poiss,A}} = 0$ and test the values of $w_{\mathrm{pre\_to\_astro}}$ that can induce astrocytic calcium transients in response to the spiking inputs from excitatory neurons. Once these parameters are determined, we set all parameters to non-zero and test the activity regime of the final neuron-astrocyte model.

#### Connectivity for `pool_type` = "block" and `pool_type` = "random"

In models with `pool_type = "block"`, connectivity parameters are set to $S_{\mathrm{pool}} = 1$ and $p_{\mathrm{third\_if\_primary}} = p_{\mathrm{primary}} = 0.2$, while ratio between cell types is $N_{\mathrm{A}} = N_{\mathrm{E}}/4 = N_{\mathrm{I}} = 100$. This means that each astrocyte can interact with the fixed pool of 4 excitatory and 1 inhibitory neuron. For each input synapse to these five neurons, the interaction with their dedicated astrocyte is established with probability $p_{\mathrm{third\_if\_primary}}$. The selected cell numbers and connection probabilities lead to each astrocyte interacting with multiple synaptic inputs to each of the five neurons, and consequently each neuron receives multiple identical SICs from the same astrocyte. This is taken into account when tuning the weight coefficient $w_{\mathrm{astro\_to\_post}}$.

In models with `pool_type = "random"` connectivity parameters are set to $S_{\mathrm{pool}} = 5$ and $p_{\mathrm{third\_if\_primary}} = 0.03$, the results are also validated for $p_{\mathrm{third\_if\_primary}} = 0.05$. The number of neurons and astrocytes is the same as before. In this case, interaction with astrocytes and synapses is established randomly with the given probability $p_{\mathrm{third\_if\_primary}}$ but each neuron can receive SIC inputs from at most $S_{\mathrm{pool}}$ astrocytes. For this choice of connectivity type and parameters, each astrocyte sends SIC inputs to several neurons, typically more than 5 (as in the case with block connectivity). Each astrocyte receives less synapses than before but some of those are synapses to the same postsynaptic neuron, and thus each postsynaptic neuron still receives multiple SIC inputs from the same astrocyte. Clearly, $p_{\mathrm{third\_if\_primary}}$ is set to a very small value for random compared to block pool type. Setting this parameter to 0.2 leads to each astrocyte interacting with an excessive number of postsynaptic neurons which quickly synchronizes the model to a network bursting regime. In order to examine both, asynchronous and bursting regimes, it is necessary to decrease $p_{\mathrm{third\_if\_primary}}$. Larger network size might permit different choice for $p_{\mathrm{third\_if\_primary}}$. Numbers of neuron and astrocyte inputs and outputs for `pool_type = "random"` are illustrated in Fig 2 for $p_{\mathrm{third\_if\_primary}} = 0.03$, and in Fig 3 for $p_{\mathrm{third\_if\_primary}} = 0.05$.

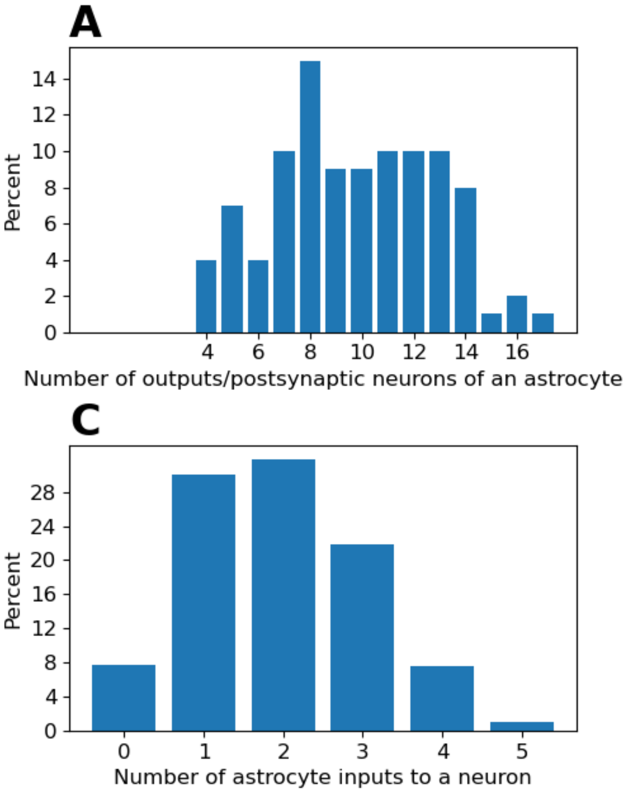


Figure 2: Connectivity in neuron-astrocyte network with `pool_type` = `"random"` and $S_{\text{pool}} = 5$. Each synapse is assigned to a random astrocyte with probability $p_{\text{third\_if\_primary}} = 0.03$. Panels show distributions of the following: A) number of postsynaptic neurons that receive SIC from the same astrocyte, B) number of presynaptic neurons that interact with the same astrocyte, C) number of astrocytes that interact through SIC with the same neuron. Pool size set to 5 means that each postsynaptic neuron receives SIC from at most 5 astrocytes. For this value of $p_{\text{third\_if\_primary}}$, neurons receive SIC inputs from between 0 and 4 astrocytes, with 2 astrocytes being the most common number of inputs.

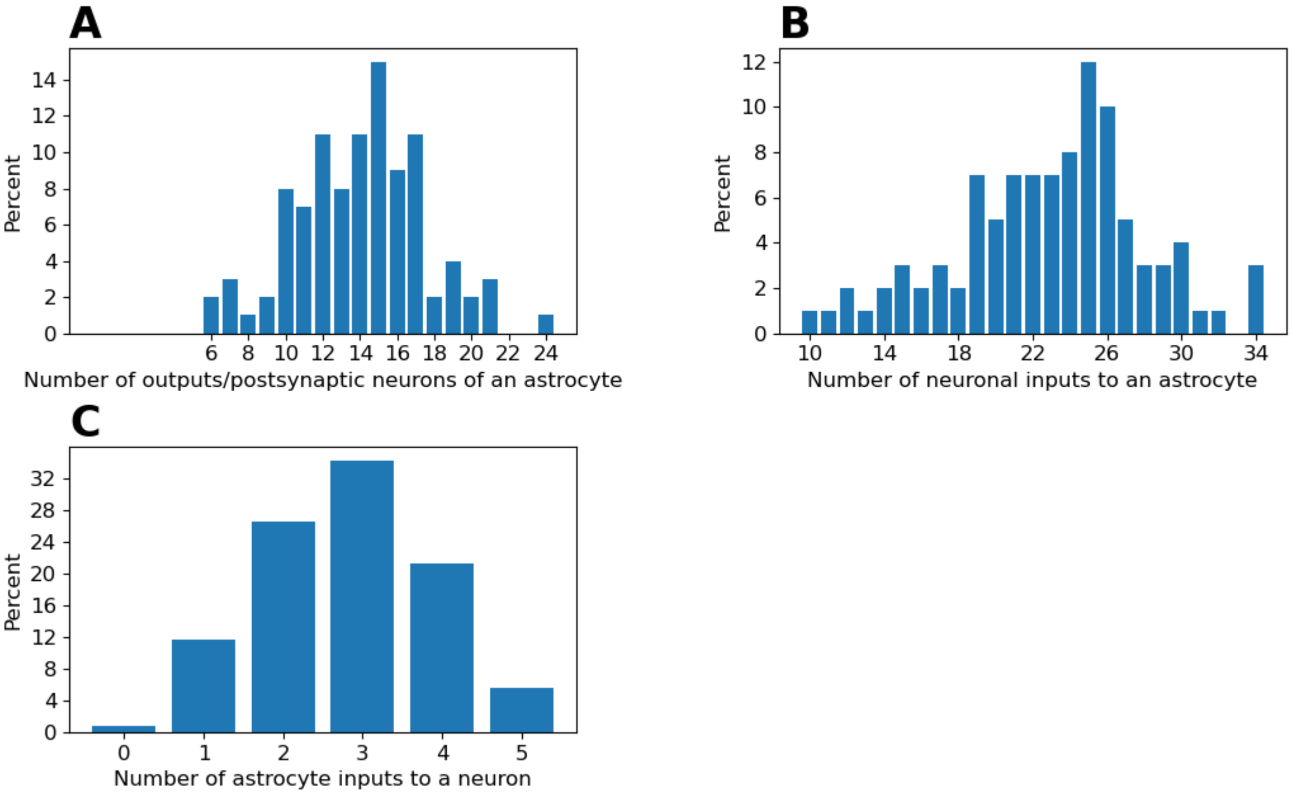


Figure 3: Connectivity in neuron-astrocyte network with `pool_type` = `"random"`, $S_{\text{pool}} = 5$, and $p_{\text{third\_if\_primary}} = 0.05$. Slightly higher probability of synapse-astrocyte interaction results in more input presynaptic neurons to an astrocyte, more postsynaptic neuron outputs from an astrocytes, and more input astrocytes to each postsynaptic neuron. The number of input astrocytes per neuron ranges from 0 to 5, with most common number being 3 astrocytes.

#### Single-neuron burst detection

To test the impact of neuron-astrocyte interaction through SIC, we evaluate single-neuron bursts arising from SIC inputs. We record burst onset, duration, frequency of bursts, and number of spikes per burst in each neuron. Burst onsets are used to evaluate SIC-induced synchronization in pairs of neurons, the results are reported in Figs 9–12 in the article, as well as in Figs 4–13 in this appendix.

For each neuron, the properties of bursts are quantified as follows:

1. Find $t_{\mathrm{sic-on}}$, $t_{\mathrm{sic-off}}$, onset and offset times for all SIC inputs to the considered neuron (regardless of the sending astrocyte).
2. Find the number of spikes within the interval $[t_{\mathrm{sic-on}}, t_{\mathrm{sic-off}} + \texttt{max\_after\_sic}]$; spiking should not start before SIC but it may continue for a short time after SIC determined by parameter `max_after_sic`. This parameter is set to 2 s for excitatory and 400 ms for inhibitory neurons.
3. The spikes have to be close enough to each other to belong to the same burst. Closeness is determined by the parameters `max_allowed_distance_exc` and `max_allowed_distance_inh` for the two neuron types. Here, these two parameters are set to 2 s and 400 ms, which is shorter than the average inter-spike-interval in these neuron types.
4. Burst onset is the time of the first spike. Burst duration is the time between the last and the first spike in the burst. Burst frequency is calculated as the total number of bursts divided by the total span of model time simulated.

Parameters used in this four-step procedure are also listed in Table C in S1 Appendix.

#### Detection of astrocyte transients

Frequency and duration of calcium transients shown in Figs 9–12 in the article, and in Figs 4-13 in this appendix are determined from calcium recordings in each astrocyte according to the following four-step algorithm. Parameters of the algorithm are also listed in Table C in S1 Appendix.

1. Threshold the calcium recording using the same threshold for triggering SIC ($\theta_{\mathrm{SIC}} + 1$ nM).
2. Noisy inputs to astrocytes cause calcium to flicker around the threshold which results in spurious threshold crossings. Multiple crossings are merged if they are too close, i.e. if they are closer than the parameter `max_allowed_distance_astro`. Fixing this parameter to 2 s gives good results for the network activity regimes considered in this study.
3. Transient onsets are ascending crossing points found above, while transient endings are descending crossing points. Duration of calcium transients is calculated as the time between ending points and onsets.
4. Frequency of astrocytic calcium transients is computed as the total number of transients recorded divided by the total span of model time simulated.

### Supplementary results for the Result section "Use case: Astrocytes promote neuronal synchronization *in silico* through neuron-astrocyte interaction"

The following figures supplement results shown in the article and have the same format as the article Figs 9–12. **Panel A)** shows calcium transients from 10 randomly selected astrocytes and distributions of frequency and duration across $N_\mathrm{A}$ astrocytes. **Panel B)** shows rasterplot of all neurons in the model, blue dots correspond to spikes recorded from $N_\mathrm{E}$ excitatory neurons and red dots correspond to spikes recorded from $N_\mathrm{I}$ inhibitory neurons. The frequency of excitatory and inhibitory neurons is shown in the lower left sub-panel, and distribution of single-neuron burst frequency, duration, and number of spikes per bursts is shown in the three lower right sub-panels. **Panel C)** illustrates calcium dynamics in one astrocyte (middle sub-panel), activity of its presynaptic neuronal inputs is shown as raster plot (top panel) and activity of its postsynaptic neurons is shown as voltage traces (bottom panel). Finally, **panel D)** confirms that neurons connected to the same astrocyte synchronize. This is confirmed using two measures - distance between single-neuron burst onsets in pairs of neurons (top panel), and pairwise cross-correlation between pairs of neurons (bottom panel). Orange bars correspond to the metric computed between neurons receiving SIC from the same astrocyte. Grey bars are computed across all pairs of neurons in the model. Two distributions (orange and grey plots) are compared using Kolmogorov-Smirnov test with Bonferroni correction done across all tests presented in the article and in this appendix.

**Choice of the random seed of the noise generator does not affect the conclusions of the study:** Fig 4 is obtained using identical model as in the article Fig 10, i.e. a network with `pool_type` = `"block"` in asynchronous activity regime, but different random seed is used when simulating this example. The obtained astrocyte and neuronal activity is the same as in Fig 10 confirming robustness of results when changing noise in the model. Neurons respond with short bursts of spikes to input SICs from astrocytes, which can be seen in rasterplots of (less active) excitatory neurons in panel B. The frequency, duration and number of neurons per bursts is quantified in the same panel. Postsynaptic neurons synchronize to astrocyte SICs and to other neurons interacting with the same astrocyte, as shown in panel C. Synchronization between neurons that receive inputs from the same astrocyte is confirmed in panel D (Kolmogorov-Smirnov test, $p < 0.0001$ for inter-burst intervals, and $p < 0.0001$ for cross-correlation coefficients after Bonferroni correction across all simulations and p-values shown in Results of the article and in this appendix).

**Conclusions obtained for "random" connectivity and asynchronous regime hold for increased probability of neuron-astrocyte interaction:** Fig 5 is obtained using the same model as the article Fig 12 (`pool_type` = `"random"`, asynchronous activity regime) except that $p_\mathrm{third\_if\_primary} = 0.05$ and synaptic weight $w_\mathrm{pre\_to\_astro}$ is decreased from 1 to 0.5 to account for the increased number of synaptic inputs. The weight $w_\mathrm{astro\_to\_post}$ is increased from 7 pA to 10 pA in this example. Although in this case neurons receive inputs from a bigger number of astrocytes, synchronization is still significantly increased between neurons that interact with (at least one) shared astrocyte (Kolmogorov-Smirnov test, $p < 0.0001$ for both measures of synchronization).

**Conclusions hold for "random" connectivity and synchronous activity regime:** Fig 6 complements the result shown in the article Fig 12. In the article Fig 12, synchronization of neuronal groups interacting with the same astrocyte is shown for in a network with `pool_type` = `"random"`, $p_\mathrm{third\_if\_primary} = 0.03$, and asynchronous network activity. In Fig 6, the synaptic coefficient $w_\mathrm{pre\_to\_astro}$ is increased from 1 to 1.5 which is sufficient to induce network-wide bursts each lasting up to 15 s. In this activity regime, overall pairwise neuronal synchronization increases, as can be seen in panel D. However, the pairs of neurons receiving SIC from the same astrocyte still synchronize significantly more than the overall network ($p < 0.0001$ for both measures of synchronization).

**Decreasing synaptic delay does not affect the conclusions of the study:** Figs 7 and 8 show that results from the main article Fig 10 and Fig 6 from this appendix do not significantly change when synaptic delay is reduced from NEST default 1 ms to a small value of 0.1 ms. Except for decreasing synaptic delay, all other parameters are the same.

**Conclusions are validated using longer simulations:** Figs 9 and 10 confirm that the article Fig 10 and Fig 6 from this appendix indeed show steady state dynamic of the model. In Figs 9 and 10 the simulations from the article Fig 10 and the supplementary Fig 6 are repeated for much longer interval of time. The models are simulated for 20 min of model time. The last 5 min are used to evaluate synchronization. As shown in the figures, the results remain the same as in much shorter simulations of 5 min used to produce all other results. All other simulations are first run for 20 s to reach the steady state dynamics, then for 5 more minutes to produce data shown in the result figures.

**Maximal step size in numerical integration schema does not change the conclusions of the study:** Figs 11 to 13 demonstrate that conclusions of this study hold regardless of parameter choice in numerical integration scheme. We simulate all models using the NEST-provided solution based on Runge-Kutta-Fehlberg-45 adaptive step size method bounded from above by the maximal simulation step size. The NEST default maximal step size of 0.1 ms is used to obtain most of the results shown in this study. In Figs 11 and 12 obtained for `pool_type` = `"block"` and asynchronous and bursting regime, respectively, we decreased the maximal step size to 0.01 ms and obtained consistent results as before.

In Fig 13, we increased the maximal step size to 0.2 ms for the model with `pool_type` = `"random"` and asynchronous activity regime and obtained the same results as before for the same model parameters. In conclusion, we simulated and reproduced all of the results shown in this study for the maximal step sizes of 0.01 ms and 0.2 ms and concluded that the choice of parameters in numerical simulation does not impact the results.

### References

M. Arizono, V. V. G. Inavalli, A. Panatier, T. Pfeiffer, J. Angibaud, F. Levet, M. J. T. Ter Veer, J. Stobart, L. Bellocchio, K. Mikoshiba, G. Marsicano, B. Weber, S. H. R. Oliet, and U. V. Nägerl. Structural basis of astrocytic $Ca^{2+}$ signals at tripartite synapses. *Nature Communications*, 11(1):1906, 2020. doi: 10.1038/s41467-020-15648-4.

L. J. Gentet, M. Avermann, F. Matyas, J. F. Staiger, and C. C. Petersen. Membrane potential dynamics of gabaergic neurons in the barrel cortex of behaving mice. *Neuron*, 65(3):422–435, 2010. doi: 10.1016/j.neuron.2010.01.006.

L. J. Gentet, Y. Kremer, H. Taniguchi, Z. J. Huang, J. F. Staiger, and C. C. Petersen. Unique functional properties of somatostatin-expressing gabaergic neurons in mouse barrel cortex. *Nature Neuroscience*, 15(4):607–612, 2012. doi: 10.1038/nn.3051.

A. Maksimov, M. Diesmann, and S. J. Van Albada. Criteria on balance, stability, and excitability in cortical networks for constraining computational models. *Frontiers in Computational Neuroscience*, 12:44, 2018. doi: 10.3389/fncom.2018.00044.

S. Nadkarni and P. Jung. Spontaneous oscillations of dressed neurons: a new mechanism for epilepsy? *Physical Review Letters*, 91(26):268101, 2003. doi: 10.1103/PhysRevLett.91.268101.

R. Naud, M. Marcille, C. Clopath, and W. Gerstner. Firing patterns in the adaptive exponential integrate-and-fire model. *Biological Cybernetics*, 99:335–347, 2008. doi: 10.1007/s00422-008-0264-7.

T. M. Pirttimaki, R. E. Sims, G. Saunders, S. A. Antonio, N. K. Codadu, and H. R. Parri. Astrocyte-mediated neuronal synchronization properties revealed by false gliotransmitter release. *Journal of Neuroscience*, 37(41):9859–9870, 2017. doi: 10.1523/JNEUROSCI.2761-16.2017.

T. Sasaki, N. Kuga, S. Namiki, N. Matsuki, and Y. Ikegaya. Locally synchronized astrocytes. *Cerebral Cortex*, 21(8): 1889–1900, 2011. doi: 10.1093/cercor/bhq256.

R. Srinivasan, B. S. Huang, S. Venugopal, A. D. Johnston, H. Chai, H. Zeng, P. Golshani, and B. S. Khakh. $Ca^{2+}$ signaling in astrocytes from $Ip3r2^{-/-}$ mice in brain slices and during startle responses *in vivo*. *Nature Neuroscience*, 18(5):708–717, 2015. doi: 10.1038/nn.4001.

J. L. Stobart, K. D. Ferrari, M. J. P. Barrett, M. J. Stobart, Z. J. Looser, A. S. Saab, and B. Weber. Long-term in vivo calcium imaging of astrocytes reveals distinct cellular compartment responses to sensory stimulation. *Cerebral Cortex*, 28(1):184–198, 2018. doi: 10.1093/cercor/bhw366.

J. Touboul and R. Brette. Dynamics and bifurcations of the adaptive exponential integrate-and-fire model. *Biological Cybernetics*, 99:319–334, 2008. doi: 10.1007/s00422-008-0267-4.

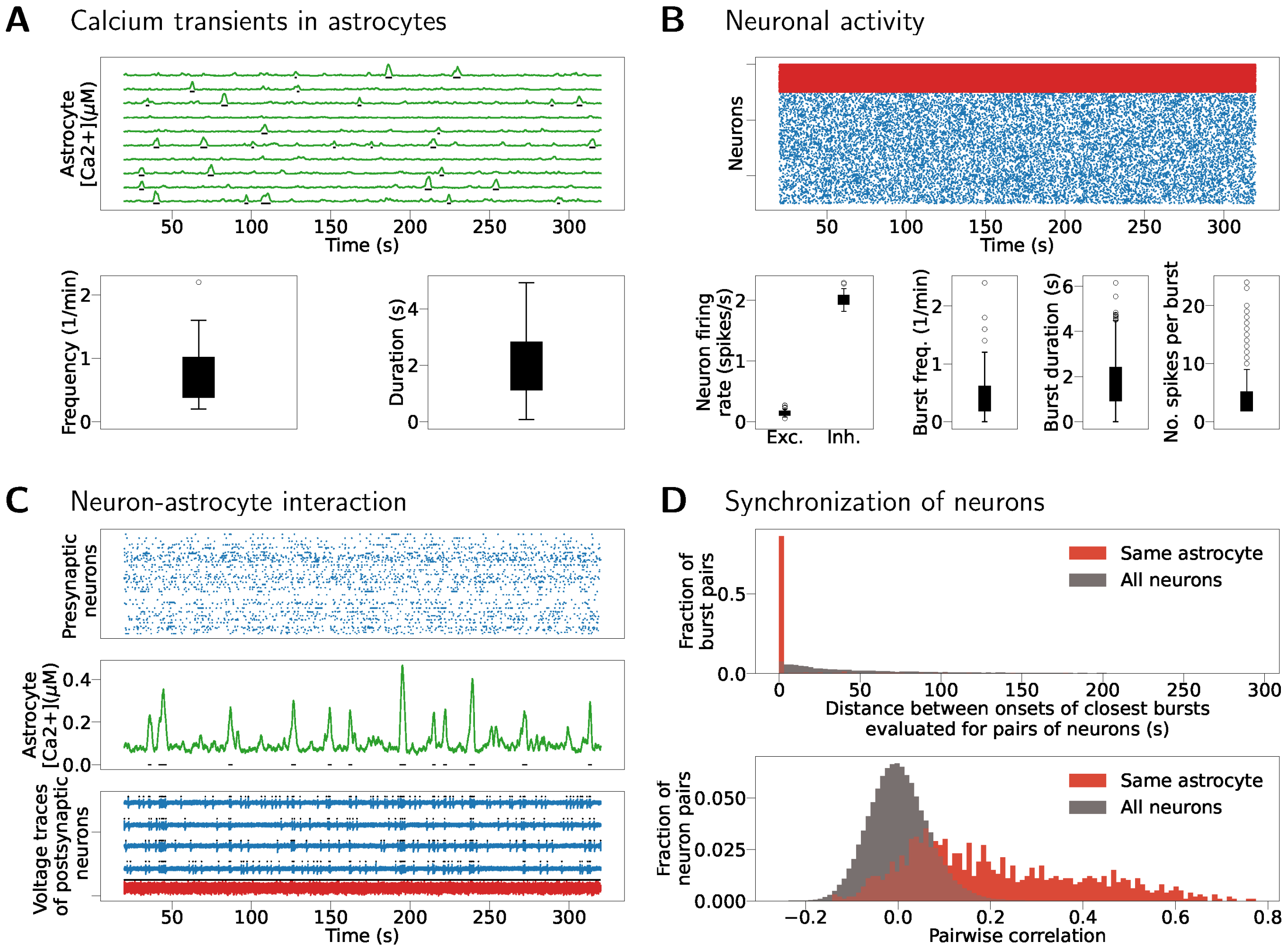


Figure 4: Fig 10 from the article (`pool_type` = `"block"`, asynchronous network activity regime) is reproduced for different seed of the random number generator to confirm robustness of this result. A) Calcium transients in 10 random astrocytes (top), frequency and duration are computed across $N_{\mathrm{A}}$ astrocytes in the model (bottom). B) Rasterplot of neuronal activity (top); blue - spike times of excitatory neurons, red - spike times of inhibitory neurons. Below, spiking frequency of excitatory and inhibitory neurons and frequency, duration, and number of spikes per neuron for single-neuron bursts induced by SIC inputs. All metrics are computed for all $N_{\mathrm{E}}$ and $N_{\mathrm{I}}$ neurons. C) Calcium dynamics in an astrocyte (middle), rasterplot of its presynaptic inputs (top), voltage traces of five excitatory (blue) and one inhibitory (red) neuron receiving SIC inputs from this astrocyte (bottom). D) Distance between onset times of single-neuron bursts evaluated for pairs of postsynaptic neurons interacting with the same astrocyte (orange) and for all pairs of neurons in the model (grey). Pairwise cross-correlation coefficients computed between pairs of postsynaptic neurons interacting with the same astrocyte (orange), and between all pairs of neurons in the model (grey).

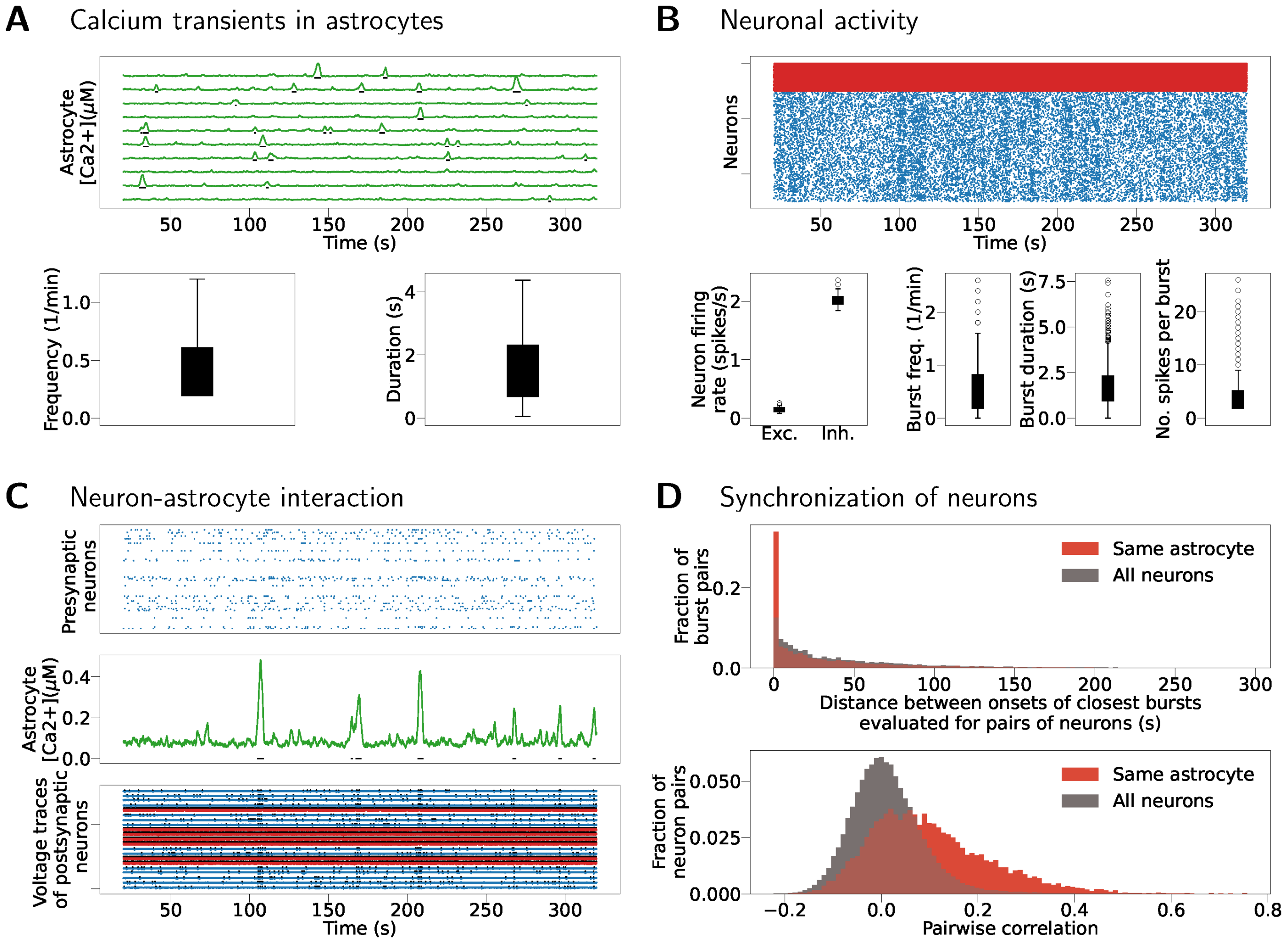


Figure 5: Fig 12 from the article (`pool_type` = `"random"`, asynchronous regime) reproduced for a higher value of $p_{\mathrm{third\_if\_primary}} = 0.05$. A) Calcium transients of 10 randomly selected astrocytes (top) and frequency and duration of calcium transients for all $N_{\mathrm{A}}$ astrocytes (bottom). B) (top) Neuronal spiking activity, blue-excitatory neurons, red-inhibitory neurons. SIC induced bursts in individual neurons and small synchronizing groups are visible in the spiking of excitatory neurons. Spiking frequency (bottom left) and the frequency, duration and the number of spikes per single neuron burst (bottom right) are computed for each neuron in the model. C) Single astrocyte (middle) with its presynaptic inputs (rasterplot, top) and postsynaptic outputs (voltage traces, bottom). D) Distance between successive neuron burst onsets in pairs of neurons and pairwise correlation between pairs of neurons.

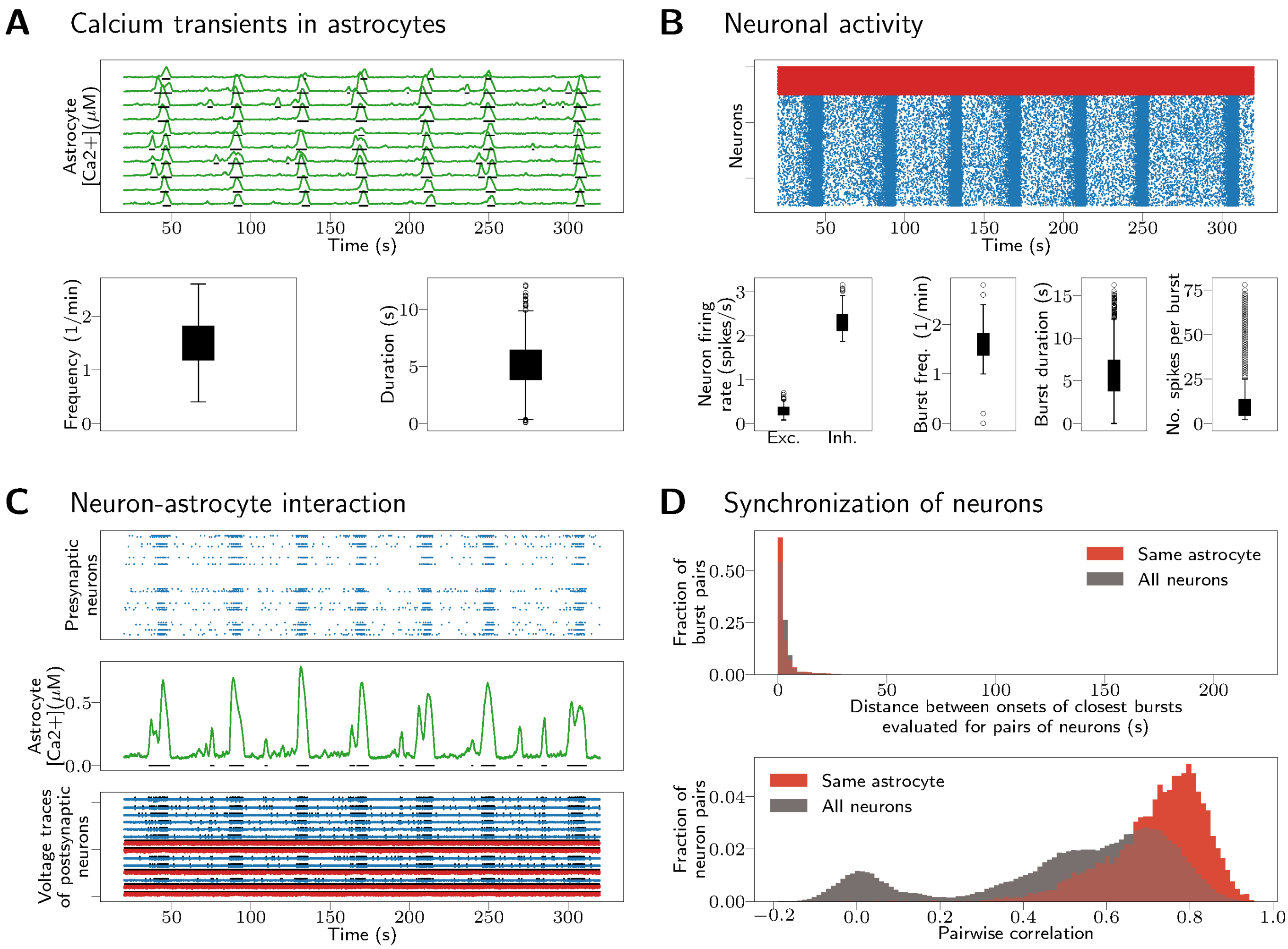


Figure 6: Model with `pool_type` = `"random"` connectivity and network bursting activity regime. The result complements Fig 12 from the article that evaluates the same connectivity but asynchronous activity regime. Switching from asynchronous activity to bursting is obtained by increasing the parameter $w_{\text{pre\_to\_astro}}$ from 1 to 1.5. A) Calcium dynamics in 10 randomly selected astrocytes (top), and frequency and duration of calcium transients in $N_A$ astrocytes (bottom). B) Neuronal spiking activity, blue-excitatory neurons, red-inhibitory neurons (top), spiking frequency (bottom left), frequency, duration and number of spikes per burst in single-neuron bursts (bottom right). C) Calcium transients in one astrocyte (middle), its presynaptic inputs (rasterplots, top) and postsynaptic outputs (voltage traces, bottom). D) Evaluation of synchronization using two metrics, the difference in onsets of single-neuron bursts (top) and the correlation coefficient (bottom).

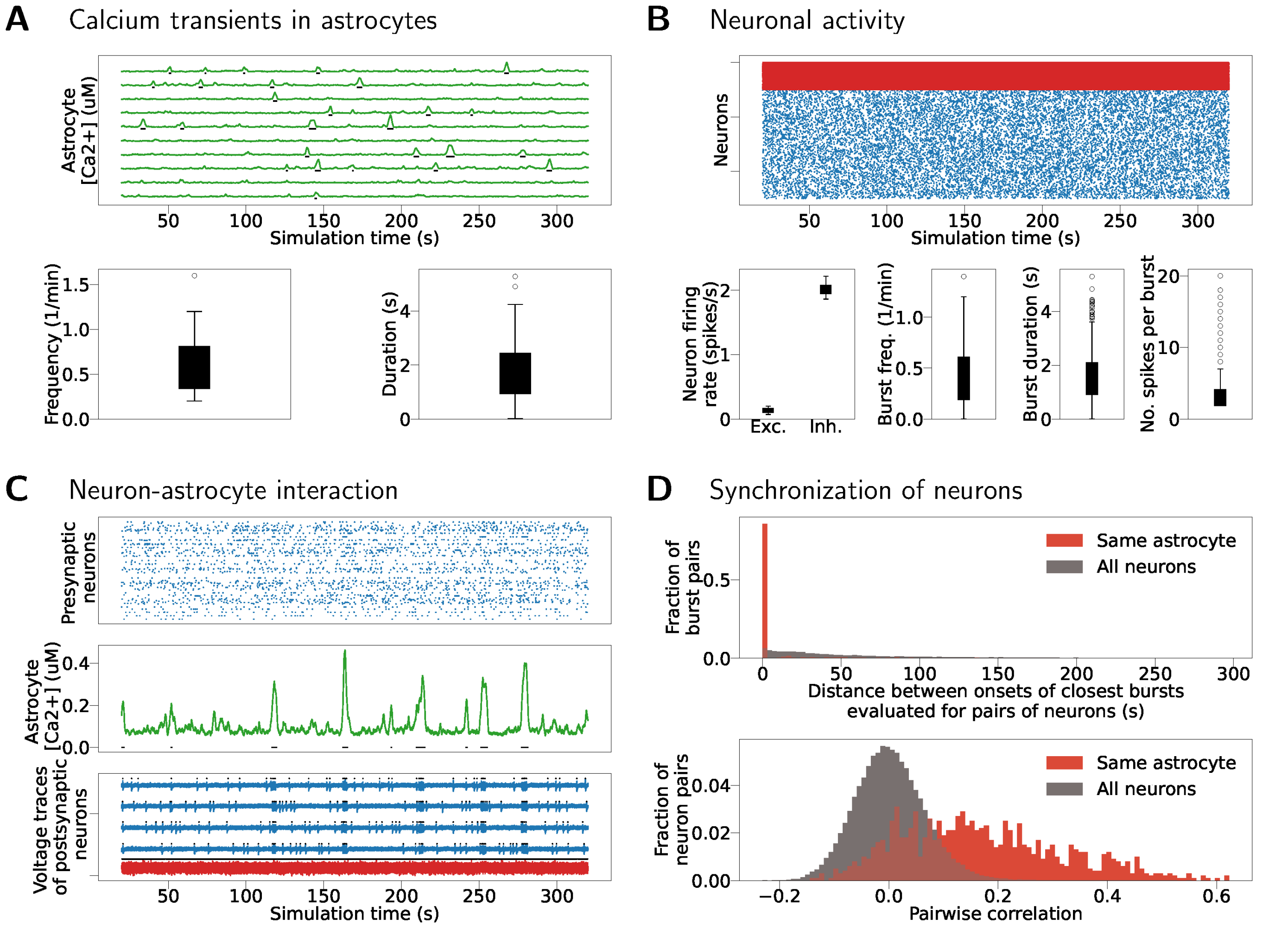


Figure 7: Simulation shown in the article Fig 10 (`pool_type` = `"block"`, asynchronous activity regime) is repeated for much smaller synaptic delay of 0.1 ms. The results minimally differ from those shown in the article Fig 10, the conclusions about astrocyte's impact on neuronal synchronization hold in this case as well. A) Calcium transients in 10 randomly selected astrocytes (top), frequency and duration of calcium transients (bottom). B) Neuronal spiking activity (top), blue-excitatory neurons, red-inhibitory neurons. Frequency of neuronal spiking (bottom left); and frequency, duration and number of spikes per single neuron bursts (bottom right). C) Calcium dynamics of one astrocyte (middle), its presynaptic inputs (up) and postsynaptic outputs (bottom). D) Evaluation of synchronization using two metrics, difference in onsets of single neuron bursts and pairwise correlation.

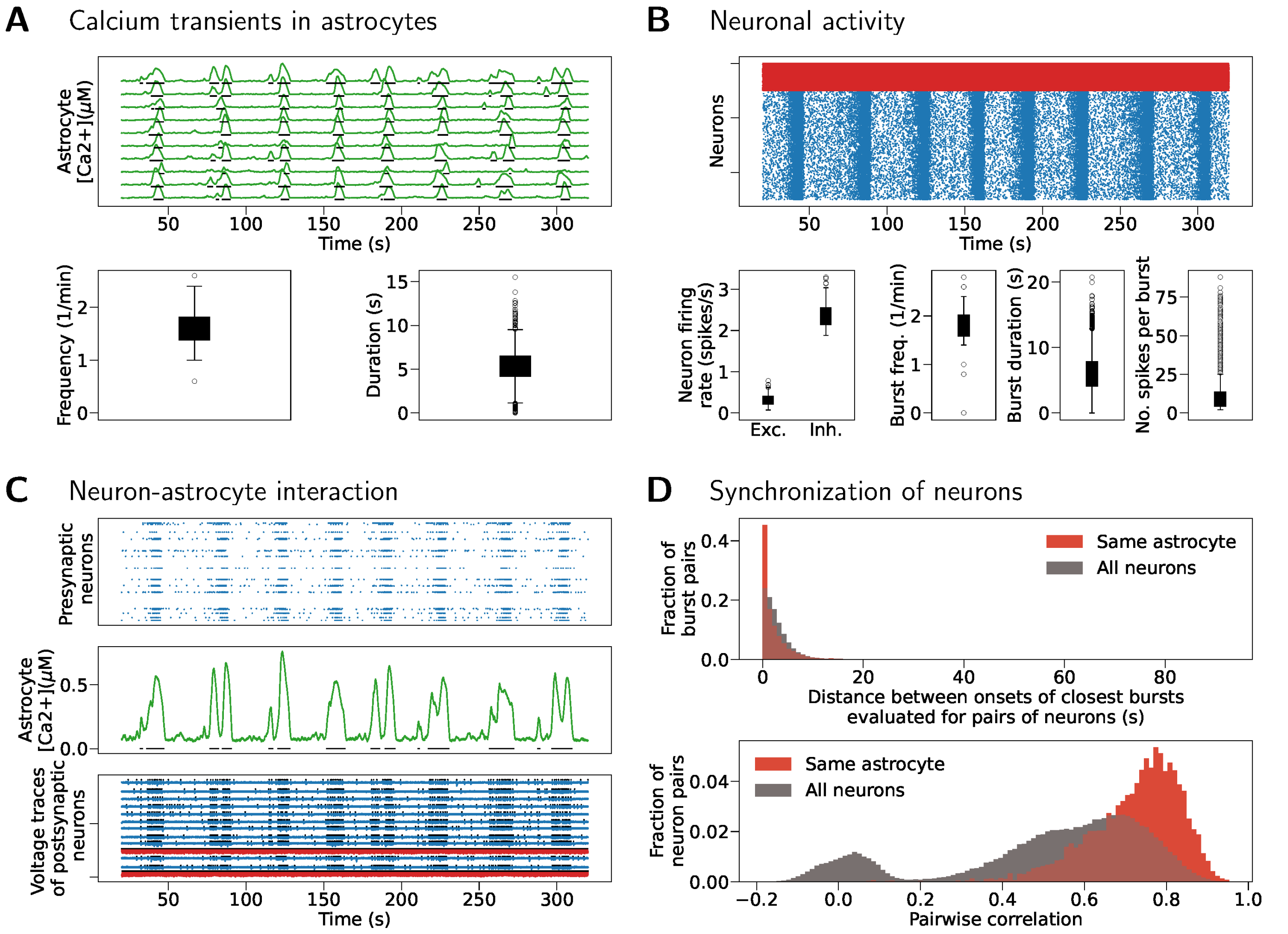


Figure 8: Fig 6 from this appendix is repeated for synaptic delay of 0.1 ms. Results remain the same in spite of faster cell-cell transfer. A) Calcium dynamics of 10 randomly selected astrocytes (top); (bottom) frequency and duration of calcium transients in all astrocytes in the network. B) Neuronal spiking activity, blue-excitatory neurons, red-inhibitory neurons (top); spiking frequency (bottom left), and neuron bursting frequency, duration and number of spikes per neuron (right) are computed for all neurons in the model. C) Calcium dynamics of one astrocyte (middle), spiking inputs from its presynaptic neurons (top) and voltage traces of its postsynaptic neurons (bottom). D) Synchronization evaluated using distance between successive burst onsets (top) and using pairwise correlation (bottom).

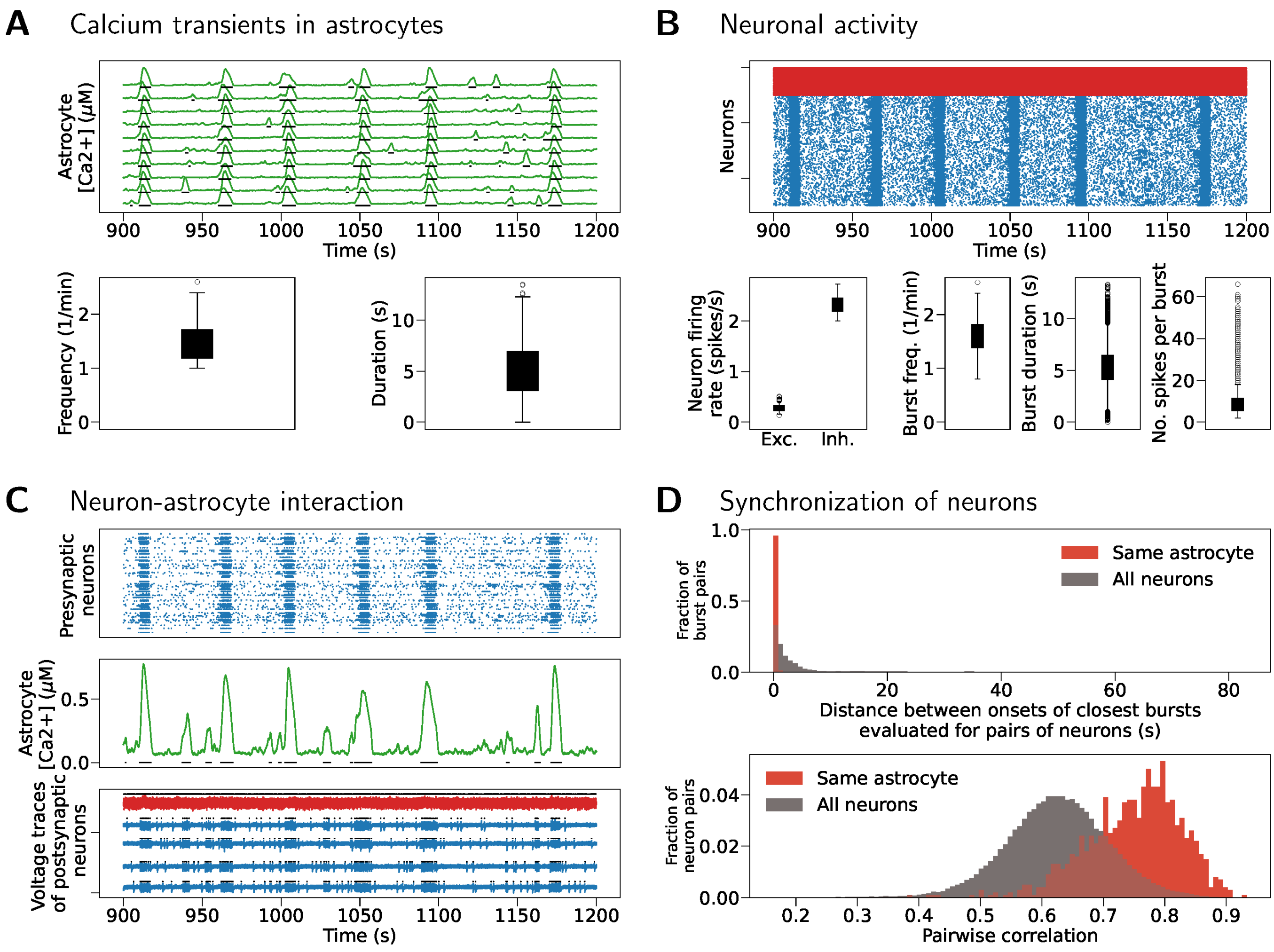


Figure 9: Network model with `pool_type` = `"block"` and bursting network activity is simulated for extended time. The model is simulated for 20 min of model time. The results do not differ significantly from the results obtained from 5 min simulations shown in the article Fig 11. A) Calcium dynamics in 10 randomly selected astrocytes (top); frequency and duration of calcium transiets (bottom) is computed for all astrocytes in the model. B) Spiking activity for excitatory (blue) and inhibitory (red) neurons. Spiking frequency (bottom left) and frequency, duration and number of spikes per single neuron bursts (bottom right) are evaluated for all neurons in the model. C) Calcium dynamics of an astrocyte (middle), its presynaptic inputs (top) and voltage traces of its postsynaptic neurons (bottom). D) Two measures of synchrony, distance between successive burst onsets (up) and pairwise correlation between neurons (bottom).

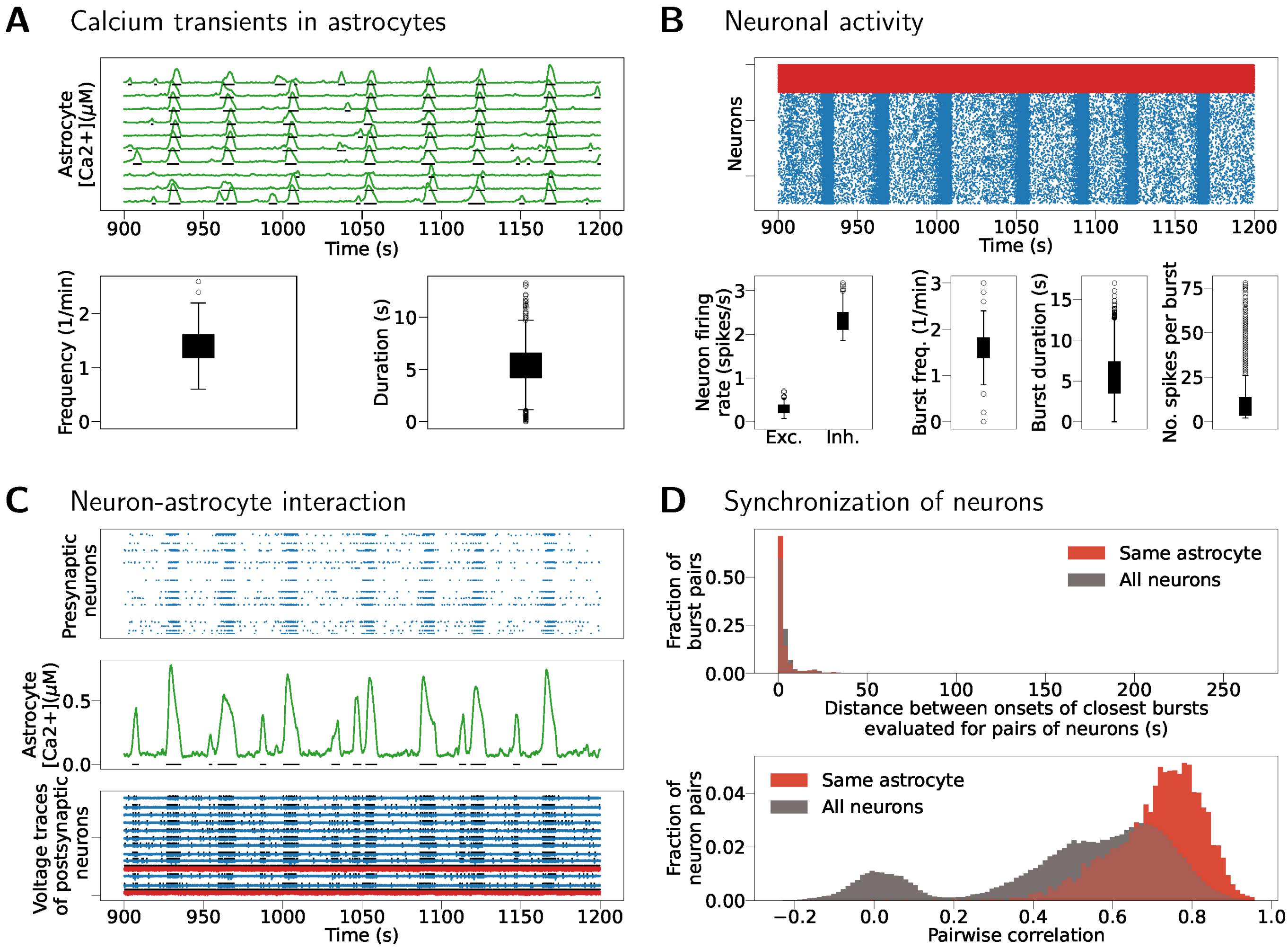


Figure 10: Network model with `pool_type = "random"` and bursting network activity is simulated for 20 min of model time. The results do not differ significantly from those obtained for 5 min simulations and shown in Fig 6 in this appendix. A) Calcium transients in 10 randomly selected astrocytes (top), frequency and duration of calcium transients in all $N_A$ astrocytes (bottom). B) Spiking activity in excitatory (blue) and inhibitory (red) neurons (top); frequency of spiking (bottom left); frequency, duration and number of spikes per single neuron burst (bottom right). C) Calcium dynamics of an astrocyte (middle), its presynaptic inputs (top) and voltage traces of its postsynaptic neurons (bottom). D) Synchronization evaluated as distance between successive burst onsets (top) and as pairwise correlation (bottom).

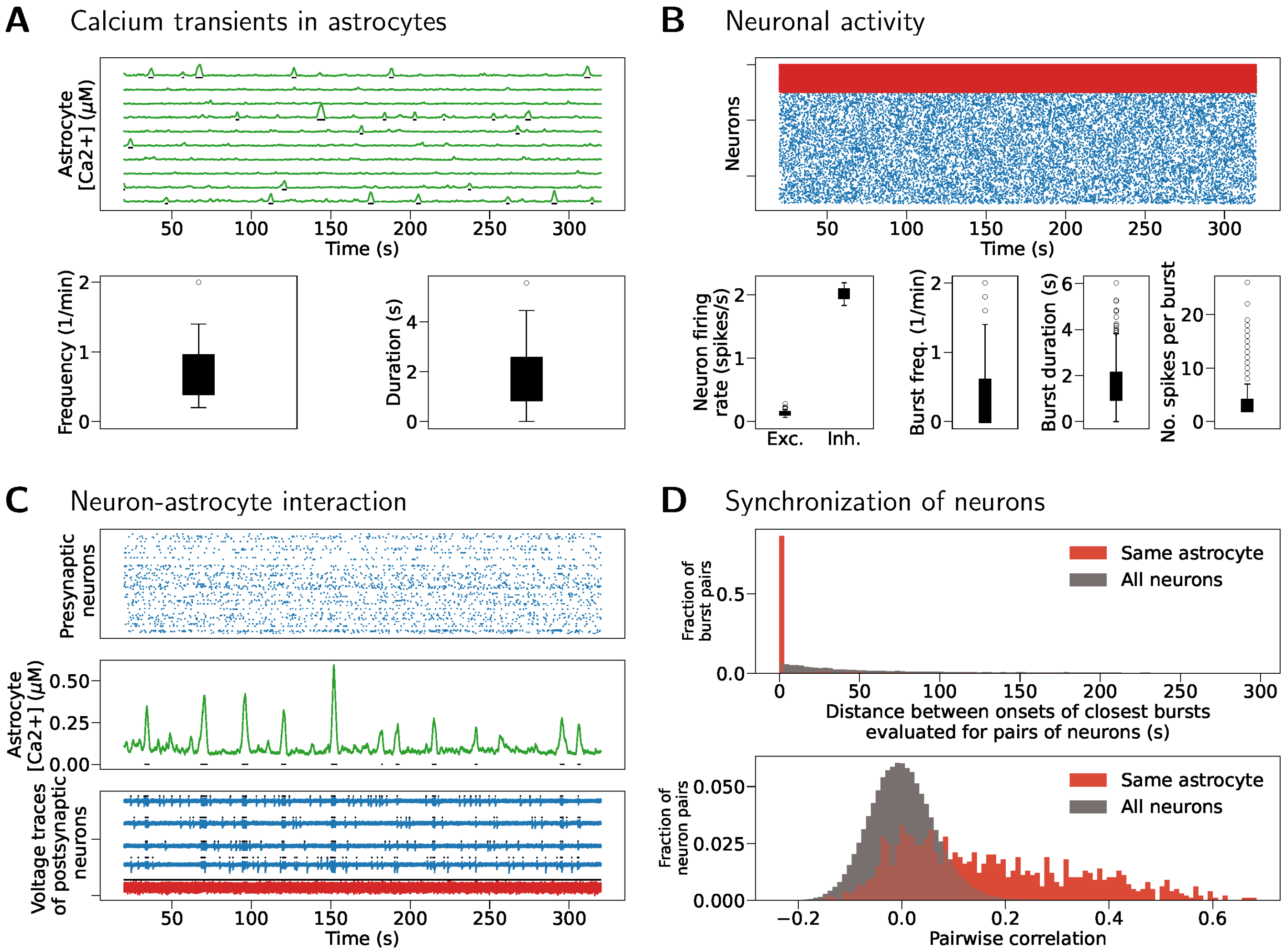


Figure 11: Article Fig 10 (`pool_type` = `"block"`, asynchronous) is reproduced for the maximal simulation step size set to 0.01 ms. A) Calcium transients in 10 randomly selected astrocytes (top), frequency and duration of calcium transients in all astrocytes (bottom). B) Spiking activity in excitatory (blue) and inhibitory (red) neurons (top); frequency of spiking (bottom left); frequency, duration and number of spikes per single neuron burst (bottom right). C) Calcium dynamics of an astrocyte (middle), its presynaptic inputs (top) and voltage traces of its postsynaptic neurons (bottom). D) Synchronization evaluated as distance between successive burst onsets (top) and as pairwise correlation (bottom).

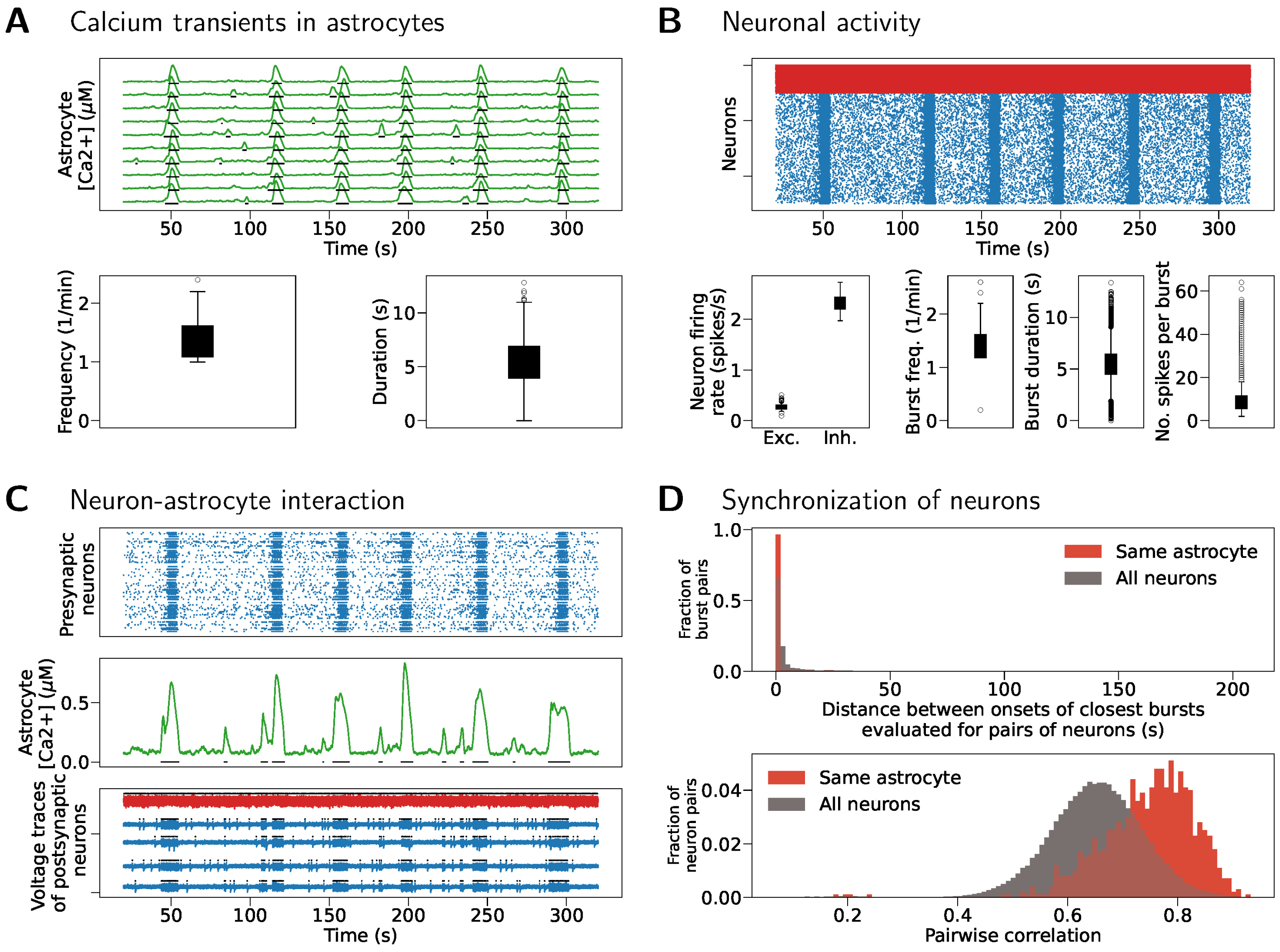


Figure 12: Article Fig 11 (`pool_type` = `"block"`, bursting) is reproduced for the maximal simulation step size set to 0.01 ms. A) Calcium transients in 10 randomly selected astrocytes (top), frequency and duration of calcium transients in all $N_{\mathrm{A}}$ astrocytes (bottom). B) Spiking activity in excitatory (blue) and inhibitory (red) neurons (top); frequency of spiking (bottom left); frequency, duration and number of spikes per single neuron burst (bottom right panels). D) Synchronization evaluated as distance between successive burst onsets (top) and as pairwise correlation (bottom).

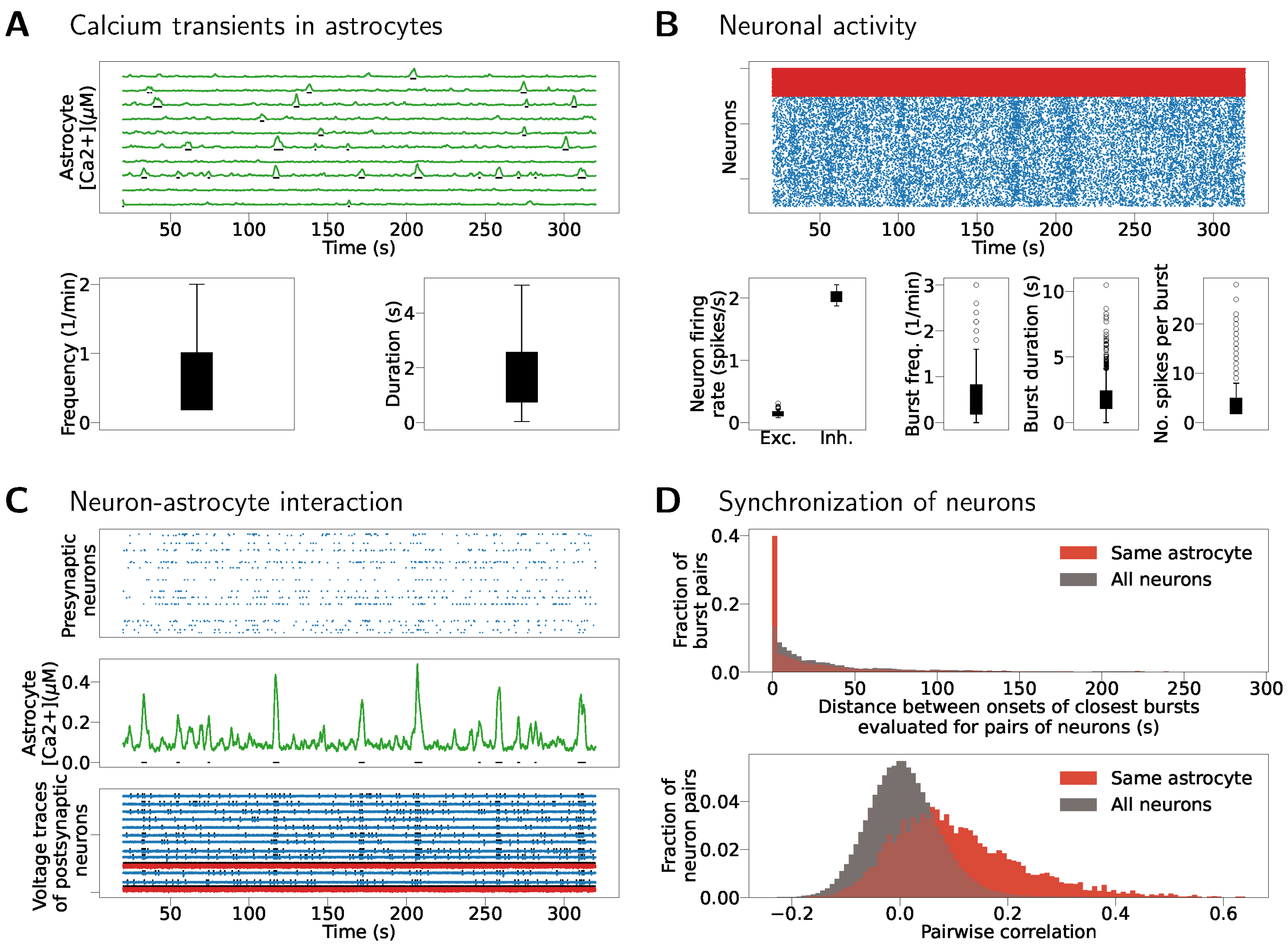


Figure 13: Article Fig 12 (`pool_type` = `"random"`, asynchronous) is reproduced for the maximal simulation step size set to 0.2 ms. A) Calcium transients in 10 randomly selected astrocytes (top), frequency and duration of calcium transients in all astrocytes (bottom). B) Spiking activity in excitatory (blue) and inhibitory (red) neurons (top); frequency of spiking (bottom left); frequency, duration and number of spikes per single neuron burst (bottom right). C) Calcium dynamics of an astrocyte (middle), its presynaptic inputs (top) and voltage traces of its postsynaptic neurons (bottom). D) Synchronization evaluated as distance between successive burst onsets (top) and as pairwise correlation (bottom).
