## Supplementary material for "Modeling neuron-astrocyte interactions in neural networks using distributed simulation": S4 Appendix

### S4 Appendix: Implementation of tripartite connectivity in the NEST Simulator

We describe here in more technical detail how tripartite connectivity generation is implemented in the NEST simulator as of version 3.8 (Graber et al., 2024).

We rename the existing NEST `ConnBuilder` base class to `BipartiteConnBuilder`. Each primary connection rule corresponds in NEST to a derived class of `BipartiteConnBuilder`, with responsibility for choosing source-target pairs and creating and parameterizing synapses according to user-provided specifications. To support third-factor connectivity, we further derive from `BipartiteConnBuilder` the classes `ThirdInBuilder` and `ThirdOutBuilder`. `ThirdInBuilder` manages the creation of connections from primary sources to third-factor nodes; since these connections are predetermined by the decisions of `ThirdOutBuilder`, no further sub-classing is allowed. `ThirdOutBuilder`, on the other hand, needs to be sub-classed for each third-factor connection rule. At present, `ThirdBernoulliWithPoolBuilder` is the only such class.

Primary and third-factor builders are combined in a new `ConnBuilder` class, which holds pointers to a builder for the primary connections which must always be present, and optionally pointers to a `ThirdInBuilder` and a `ThirdOutBuilder` which must either both be absent (normal bipartite connectivity) or both be present (tripartite connectivity). Each `BipartiteConnBuilder` holds a pointer to a `ThirdOutBuilder` (null if only bipartite). This allows the primary connection builder to trigger the check for a third-factor connection for every primary connection created by calling `ThirdOutBuilder::third_connect()`. The `ThirdOutBuilder` in turn holds a pointer to a `ThirdInBuilder`, so that for each "third out" connection created, the `ThirdOutBuilder` can call `ThirdInBuilder::register_connection()` to inform the latter about the primary source-third factor node pair that needs to be connected. The `ThirdInBuilder` maintains two closely connected data structures to manage these pairs: (1) a buffer storing tuples consisting of the global node IDs of the source and the third-factor node, and the MPI rank which manages the third-factor node, and (2) an array of counters recording how many source-target pairs need to be transmitted to each MPI rank. In order to allow for collision-free updating of these data structures during multi-threaded connectivity generation, a separate buffer and set of counters is maintained for each thread within an MPI process.

When the `Connect()` or `TripartiteConnect()` functions in NEST are called, the simulation kernel first creates a `ConnBuilder` instance with a `BipartiteConnBuilder` instance representing the primary connection rule and, for tripartite connectivity, a `ThirdInBuilder` and a `ThirdOutBuilder` instance. It then calls the `connect()` method on the primary connection builder, which creates all connections prescribed by the primary rule and triggers the `ThirdOutBuilder` as required to create the third-out connections. Once this method returns, the `connect()` method is called on the `ThirdInBuilder`. It first determines locally on each MPI rank the largest number of source-third pairs that needs to be sent to any other rank. This number is shared with all MPI ranks, so that each rank then can determine the globally largest number of pairs to be sent from any rank to any other rank, and transmission buffers are resized correspondingly and filled with local data. Using `MPI_Alltoall()`, pairs are then exchanged between ranks. Afterwards, on any given rank, all source-third pairs are available for which the rank manages the third factor node. Each rank then creates the prescribed connections in thread-parallel fashion, completing the connection process.

The presently implemented `third_factor_bernoulli_with_pool` rule is implemented by subclassing `ThirdOutBuilder` as `ThirdBernoulliWithPoolBuilder`. When the `third_connect()` method of this subclass is called with a given primary connection, it first performs a Bernoulli experiment with given probability to determine whether to create a third-factor connection. If so, the third-factor pool for the current target neuron is determined: For block-type pools, pool members are found by suitable indexing into the astrocyte population. For random pools, the pool is created when a given target neuron is connected to a third-factor element for the first time. The actual third-factor element to attach is then chosen from the pool with uniform probability, the "third out" connection created and the source-third pair registered with the `ThirdInBuilder`. The pool is stored in an associative container, so that later third-factor connections to the same target can be drawn from the same pool.

Further third-party rules can be added as new subclasses of `ThirdOutBuilder`, either in the NEST code base or in a NEST extension module. The subclass only needs to define a constructor to extract parameters for the new connection rule and override the `third_connect()` method to implement the new rule.

#### References

S. Graber, J. Mitchell, A. C. Kurth, D. Terhorst, J.-E. W. Skaar, C. M. Schöfmann, S. Kunkel, G. Trensch, N. Haug, D. Mallett, P. Y. Andriyovich, X. Otazu Porter, A. Y. Lee, and H. E. Plesser. NEST 3.8, July 2024. URL `https://doi.org/10.5281/zenodo.12624784`.
